# Single cell transcriptome profiling reveals pathogenesis of Bullous Pemphigoid

**DOI:** 10.1101/2024.06.28.601141

**Authors:** Guirong Liang, Chenjing Zhao, Qin Wei, Suying Feng, Yetao Wang

## Abstract

Bullous pemphigoid (BP) triggers profound functional changes in both non-immune and immune cells in the skin and circulation, yet the underlying mechanisms remain elusive. In this study, we conducted single-cell transcriptome analysis on donor-matched lesional and non-lesional skin, as well as blood samples from BP patients. Lesional skin non-immune cells coordinately upregulated metabolism, wound healing, immune activation, and cell migration associated pathways. Skin LAMP3^+^ DCs derived from cDC2 exhibited higher pro-inflammatory signatures than those from cDC1, and VEGFA^+^ mast cells driving BP progression, were predominantly from lesional skin. As BP patients transition from active to remission stages, blood B cell function shifts from differentiation and memory formation to heightened type 1 interferon signaling and reduced IL-4 response. Blood CX3CR1^+^ZNF683^+^ and LAG3^+^ exhausted T cells exhibited the highest TCR expansion among clones shared with skin CD8^+^T cells, suggesting they likely represent BP-reactive cells fueling skin CD8^+^T cell clonal expansion. Clinical parameters for BP severity correlated positively with blood NK cell IFN-γ production, whereas correlated negatively with NK cell AREG production. In lesional skin, NK cell-keratinocyte interactions exhibited reduced AREG-EGFR and enhanced IFNG-IFNGR1/2 signaling. NK cell-derived AREG mitigates IFN-γ-induced keratinocyte apoptosis, highlighting a crucial balance between AREG and IFN-γ in BP progression. These results reveal significant functional shifts in BP pathology within skin and blood cells and suggest new therapeutic targets for disease management.

**Graphical abstract:** 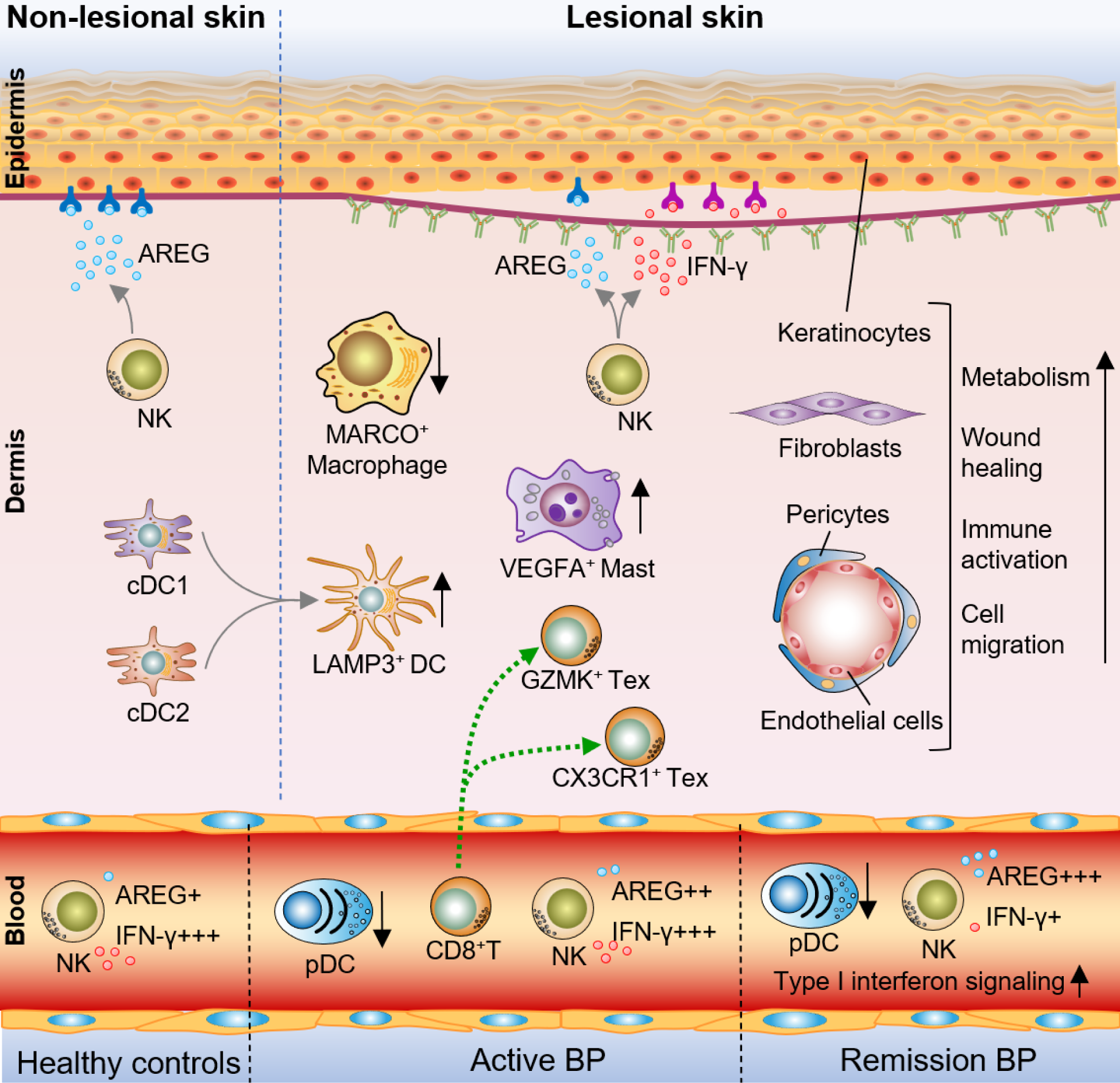

## Introduction

Bullous pemphigoid (BP) is the most common autoimmune subepidermal blistering disease, induced by antibodies against hemidesmosome anchoring proteins BP180 and BP230 (encoded by COL17A1 and DST respectively) that links basal keratinocytes to the basement membrane zone^1,2^. BP is clinically characterized by intense pruritus and bullous lesions, and usually occurs in elderly (>60 years old)^3^. Increasing evidence shows that the influence of BP is systemic, the enhanced mortality is associated with multiple comorbidities, such as hematological abnormalities, tumors, cardiovascular diseases and neurological disorders^4–9^. However, the interrelationship between BP and its complications was not clear. Systematically investigating the pathological changes in the skin and other tissues may provide new insights that benefit the clinical control of disease progression.

Besides self-reactive antibodies, multiple cell types in the skin and the blood contributed to pathogenesis of BP. For example, the elevated activity of Th2 cells, type innate lymphoid cells (ILC2s) and eosinophils contribute to blister formation and pruritus through production of IL-4, IL-5, IL-13, IL-31, MMP9, ROS and recruiting BP180 reactive IgE^1^. The absence of Treg cells lead to the generation of autoantibodies against BP230 in the BP mouse model^10^. Basophils, neutrophils and monocytes collectively determine BP progression in response to inflammatory cytokines or chemokines^11–13^. Natural killer (NK) cells enhance the inflammation of blistering skin disease by producing IFN-γ, TNF-α and releasing cytolytic granzymes^14–16^. Granzyme B cleaves BP180 and accumulates at dermal-epidermal junction, in blister fluid and in lesional skin of BP patients to promote disease progression^15,16^, however, the regulatory role of blood NK cells is important to restrict type 2 inflammation in atopic dermatitis (AD)^17^, indicating NK cells may play a paradoxical role in BP. Unbiased analysis of the cells in the skin and the peripheral blood mononuclear cells (PBMCs) should enhance the understanding of their contributions to BP pathology.

To reduce the comorbidities induced by exposure of pathogens and inflammatory irritants, the priority of BP treatment is to restore the homeostasis and integrity of lesional skin. The metabolism of keratinocytes and infiltrated immune cells is closely related with skin homeostasis^18,19^, for example, aberrant activation of MTOR leads to enhanced proliferation of keratinocytes in psoriasis^20,21^. Inhibition of glycolysis using 2-deoxy-D-glucose (2-DG) ameliorates BP-like epidermolysis bullosa acquisita (EBA) in mouse model^22^. By regulating metabolic pathways, epidermal growth factor receptor (EGFR) signaling is pivotal for the survival, proliferation and differentiation of epithelial cells in normal and disease conditions^23^. We recently showed that human natural killer (NK) cells serve as a major innate source of amphiregulin (AREG)^24^, a EGFR ligand playing important roles in maintaining the integrity of epithelium^25–27^, whereas, the function of AREG^+^NK cells and their interplay with proinflammatory IFN-γ^+^NK cells in BP remains unclear.

In this study, we conducted single-cell transcriptome analysis of paired lesional and non-lesional skin as well as donor-matched blood samples from BP patients. Our findings offer a detailed insight into the cellular dynamics in both skin and blood affected by BP, highlighting significant functional changes in immune and non-immune cells linked to the disease. These results indicate potential new targets for therapeutic intervention to enhance clinical treatment of BP.

## Results

### Experimental design and overview of single-Cell analysis of clinical BP samples

To understand the pathogenesis of BP, site and donor matched lesional (L) and non-lesional (NL) skin cells from untreated patients, PBMCs from skin samples matched BP patients in active or remission stage and from healthy donors were subjected to single cell transcriptome analysis (active BP was defined by the appearance of new skin lesions; remitted BP was defined by the healing of all lesions and the absence of new blisters or urticarial lesions for at least one month) (Figure 1A and Table S1). 113,044 cells from the skin and 38,854 cells from the blood were identified by Cell Ranger with an average depth of 45,095 mean reads/cell. After filtering by Seurat, 95,257 and 33,921 cells were identified from the skin and the blood respectively (Table S2 and Methods). Cells derived from the skin formed 12 clusters representing epidermal non-immune cells (keratinocytes, melanocytes), dermal non-immune cells (fibroblasts, endothelial cells, pericytes, vascular smooth muscle cells) and immune cells (T, B, NK, mast, dendritic cells and monophagocytes) in uniform manifold approximation and projection (UMAP) (Figures 1B and 1C). PBMCs derived from healthy donors and BP patients mainly formed 4 clusters representing T, B, NK cells and monophagocytes in UMAP (Figures 1B and 1C). Compared with non-lesional skin, the percentage of epidermal and dermal non-immune cells were not significantly altered (Figure 1D). Compared with healthy people, an increased monocytes and decreased lymphocytes percentage in the blood were detected in BP patients in both active and remission stages (Figures 1D and 1E), indicating an altered circulating immune environment in these patients.

**Figure 1.**
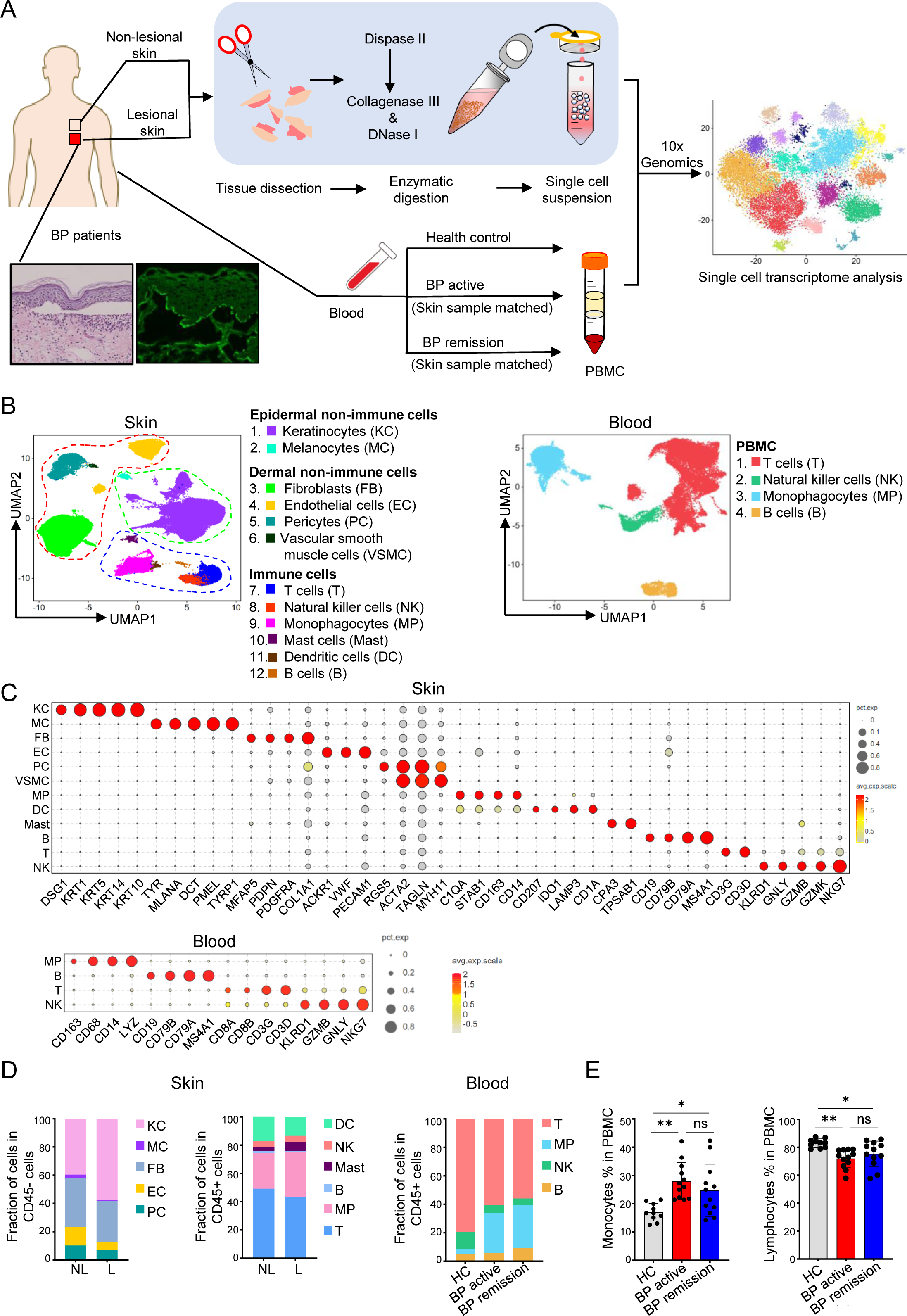
Single cell atlas of BP skin and blood samples. (A) Experimental design of this study. (B) UMAP of cells from four BP non-lesional and four lesional skin samples (left), and cells from PBMCs of two healthy donors, two active BP patients, and two remitted BP patients (right). (C) The expression of cell type specific markers for (B). (D) Histogram showing the percentage of CD45^−^ and CD45^+^ cells in BP non-lesional (n=4) and lesional skin (n=4), and the percentage of blood cells from healthy donors (n=2) and BP patients in active (n=2) and remission (n=2) stages. (E) Percentage of monocytes (left) and lymphocytes (right) in PBMCs from healthy donors (n=10) and BP patients in active (n=12) and remission (n=12) stages by complete blood count. Two-tailed unpaired t test, ns, not significant, *, p<0.05; **, p<0.01. Data are mean with s.e.m.

### Coordinated responses of non-immune cells in BP induced skin pathogenesis

Emerging evidence suggests that epidermal and dermal non-immune cells play pivotal roles in modulating immune responses in inflammatory skin diseases^28−31^. To explore their transcriptional changes in BP, keratinocytes (KCs), fibroblasts (FBs), pericytes (PCs), and endothelial cells (ECs) from BP patients were reclustered (Figure S1A-S1D and 2A). Analysis of differentially expressed genes (DEgenes) between lesional and non-lesional skin revealed different clusters within each cell type showed varying responses to BP pathology (Figure 2B). For instance, differentiated KC2 had fewer unique DE genes compared to other KC clusters, while secretory-reticular and stressed FB had more unique DE genes than other FB clusters. In lesional skin, hair follicle-associated cells (HFACs) uniquely upregulated genes enriched in WNT signaling (CTNNB1, CTNNBIP1) and cell cycle regulation (CDK4), whereas stress response (EEF1A1, UBC) and metabolism of amino acids and lipid associated genes (LARS1, MPST, OSBP, PLPP2), were uniquely downregulated in differentiated KC1 (Table S3). Arterioles uniquely upregulated genes related to notch signaling (NOTCH4, JAG1, and JAG2), while venules upregulated genes associated with TGF-β signaling (TGFB1, LTBP3) and elastic fiber formation (ELN, FBLN2) (Table S3).

**Figure 2.**
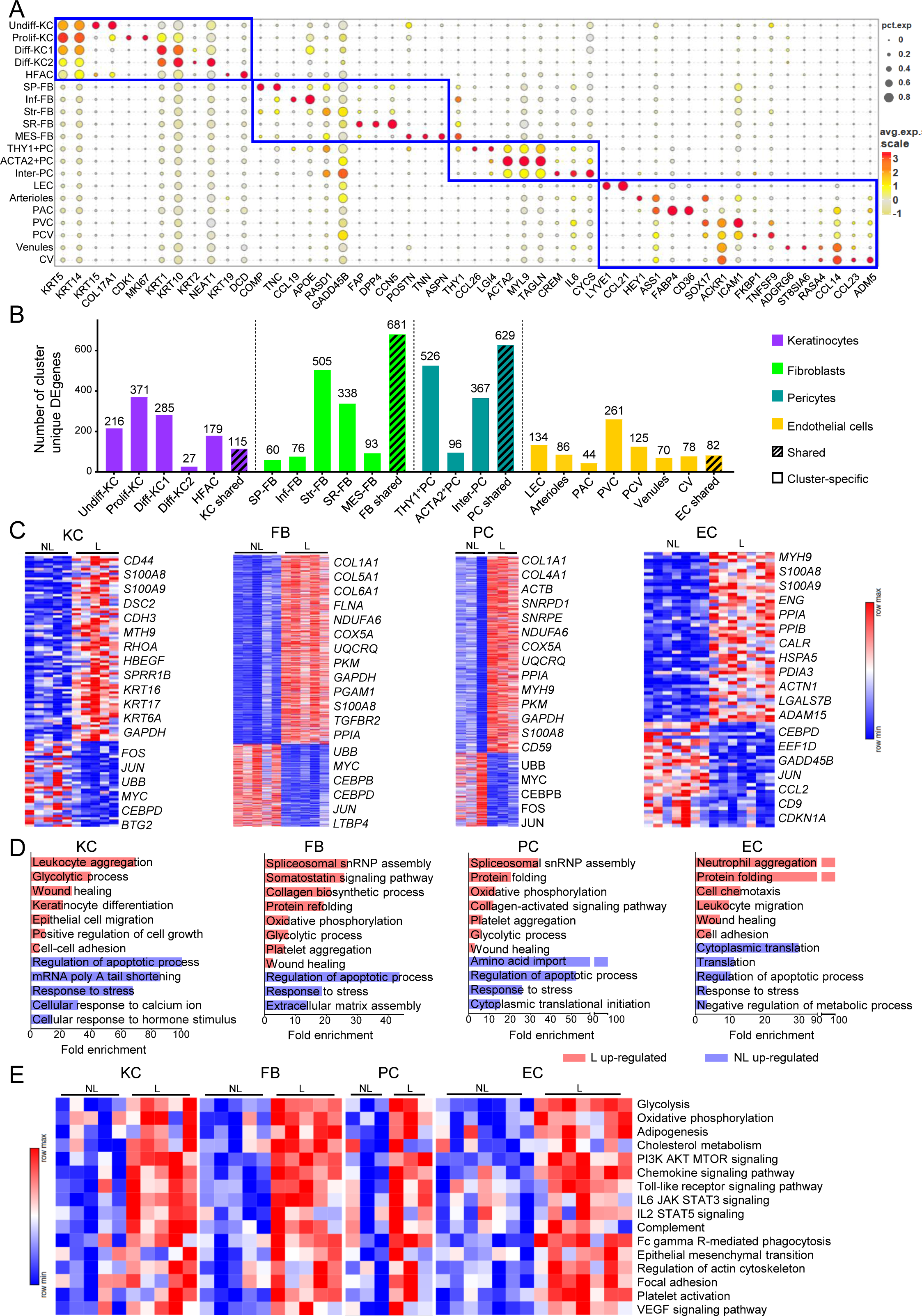
Coordinated responses of non-immune cells in BP induced skin pathogenesis. (A) The expression of cell type specific markers for clusters of keratinocytes (KC), fibroblasts (FB), pericytes (PC) and endothelial cells (EC). Undifferentiated KC (undiff-KC), Proliferating KC (Prolif-KC), Differentiated KC1 (Diff-KC1), Differentiated KC2 (Diff-KC2), Hair follicle associated cells (HFAC), Secretory-papillary FB (SP-FB), Pro-inflammatory FB (Inf-FB), Stressed FB (Str-FB), Secretory-reticular FB (SR-FB), Mesenchymal FB (MES-FB), THY1^+^ pericytes (THY1^+^PC), THY1-ACTA2^high^ pericytes (ACTA2^+^PC), Intermediate pericytes (Inter-PC), Lymphatic endothelial cells (LEC), Post-arterial capillaries (PAC), Pre-venular capillaries (PVC), Post-capillary venules (PCV), Collecting venules (CV). (B) Number of cluster unique and shared DEgenes between BP non-lesional and lesional KC, FB, PC and EC. The DEgenes was determined by Seurat FindAllMarkers (*P*<0.05, Log_2_FC>0.36). (C) Heatmap of cluster shared DEgenes between BP non-lesional and lesional KC, FB, PC and EC. (D) Enriched pathways of cluster shared up-regulated (red) and down-regulated (blue) genes of KC, FB, PC, and EC in BP lesional skin by reactome analysis (FDR<0.05). (F) GSVA of KC, FB, PC, and EC clusters in BP non-lesional and lesional skin.

Our analysis revealed remarkable shared transcriptional changes among skin non-immune cell clusters. Notably, fibroblast and pericyte clusters had substantially more shared DE genes (681 and 629, respectively) compared to keratinocyte and endothelial cell clusters (115 and 82, respectively) (Figure 2B).

In keratinocyte clusters, the shared DEgenes play critical roles in inflammation and proliferation stage of wound healing^32^, for instance, the upregulated genes in lesional skin were involved in regulating leukocyte aggregation (S100A8, S100A9, CD44), epithelial cell differentiation (SPRR1B, KRT16), growth (HBEGF, KRT17), migration (MYH9, RHOA) and adhesion (DSC2, CDH3), whereas, the downregulated genes were involved in regulation of apoptotic process (UBB, MYC), and cellular response to calcium (FOS, JUN) (Figure 2C and 2D). In fibroblast clusters, genes that function in tissue remodeling of injured skin, such as collagen synthesis (COL1A1, COL5A1), platelet aggregation (FLNA) were commonly upregulated in lesional skin. Additionally, collagen activated signaling pathway (COL1A1, COL4A1) was enriched by upregulated genes in pericytes clusters, indicating a crosstalk of these two cell types in BP lesional skin (Figure 2C and 2D). In endothelial cell clusters, genes associated with leukocyte migration (MYH9), chemotaxis (S100A8, S100A9) and wound healing (ENG, PPIA) were upregulated in lesional skin, highlighting their critical roles in regulating lymphocytes recruitment and inflammation (Figure 2C and 2D). Moreover, DEgenes associated with wound healing (PPIA, S100A8), response to stress (CEBPD, JUN), as well as glycolysis (GAPDH, PKM, PGAM1) and oxidative phosphorylation (NDUFA6, COX5A, UQCRQ) (excluding endothelial cells) were consistently detected across all skin non-immune cell types (Figure 2C and 2D).

The observed distinct and overlapping transcriptional responses among different skin non-immune cells in BP patients underscore their coordinated involvement in immune modulation, wound healing, and tissue remodeling in BP pathology. Gene Set Variation Analysis (GSVA) further elucidated these pathways, showing convergent transcriptional changes in KCs, FBs, PCs, and ECs affected by BP. In BP lesional skin, increased activity was commonly observed in pathways related to cellular metabolism, chemotaxis, immune activation, and cell migration (Figure 2E). Overall, these results offer a comprehensive insight into the transcriptional profiles of non-immune skin cells affected by BP.

### The features of myeloid cells in BP patients

Myeloid cells play a significant role in creating the inflammatory environment associated with the formation of blisters and skin lesions in BP patients. In the skin, five subsets of macrophages (THBS1^+^Mac, IER2^+^Mac, STAB1^+^Mac, MARCO^+^Mac and TREM2^+^Mac) and five subsets of dendritic cells (cDC1, cDC2, LAMP3^+^DC, cycling DC and langerhans cells) have been identified (Figure 3A and 3C). Additionally, CD14^+^ monocytes, CD16^+^ monocytes, cDC, and pDC have been identified in the blood using established markers (Figure 3B and S2A)^33,34^.

**Figure 3.**
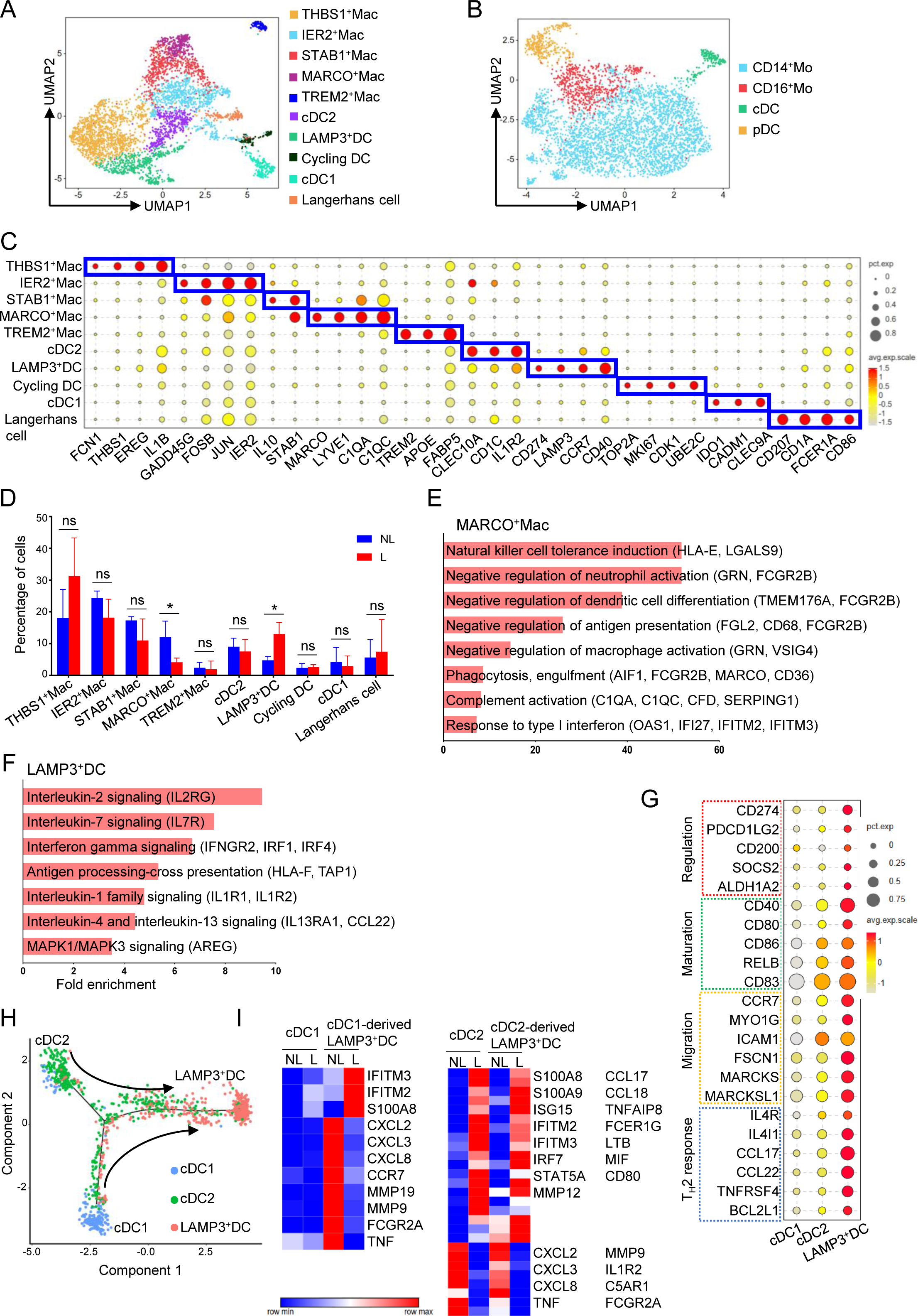
Characterization of mononuclear phagocytes in BP patients. (A, B) UMAP of myeloid cells from BP non-lesional and lesional skin (A), and from the blood of healthy donors and BP patients in active and remission stages (B). (C) Dot plot of specific markers for BP skin myeloid cell clusters. (D) Percentage of myeloid cell clusters in BP non-lesional and lesional skin. Mann-Whitney test, ns, not significant; *, p<0.05. Data are mean with s.e.m. (E, F) Enriched pathways of MARCO^+^Mac (E) and LAMP3^+^DC highly expressed genes by reactome analysis (FDR<0.05). (G) The expression of genes associated with different functions among cDC1, cDC2 and LAMP3^+^DC in the skin of BP patients (H) Pseudotime analysis of BP skin DC subsets. (I) Heatmap of inflammation-related DEgenes between BP non-lesional and lesional cDC1, cDC2, cDC1-and cDC2-derived LAMP3^+^DC.

BP lesional skin showed a decrease in MARCO^+^ macrophages compared to non-lesional skin (Figure 3D). Consistent with previous reports^35,36^, these macrophages were involved in natural killer cell tolerance induction (HLA-E, LGALS9), negative regulation of neutrophil and macrophage activation (GRN, FCGR2B, VSIG4), dendritic cell differentiation (TMEM176A, FCGR2B), antigen presentation (FGL2, CD68, FCGR2B), and phagocytosis (AIF1, FCGR2B, MARCO, CD36), and were also involved in complement activation (C1QA, C1QC, CFD, SERPING1) and response to type 1 interferon (OAS1, IFI27, IFITM2, IFITM3) (Figure 3E).

Conversely, LAMP3^+^DCs, which mature from conventional DCs^34^, were increased in lesional skin (Figure 3D). Similar to the signatures identified in tumors, skin LAMP3^+^DCs were enriched with genes associated with immunoregulation (CD274, PDCD1LG2, SOCS2), maturation (CD40, CD80, CD83), migration (MYO1G, CCR7, FSCN1), and Th2 response (IL4R, IL4I1, CCL17, CCL22) compared to cDC1 or cDC2 (Figures 3F and 3G). Furthermore, pseudotime analysis, akin to that of tumor LAMP3^+^DCs^34^, showed that skin LAMP3^+^DCs were potentially derived from both cDC1s and cDC2s (Figure 3H).

The impact of BP pathology on LAMP3^+^DCs derived from cDC1s and cDC2s was investigated next. In BP lesional skin, ISGs and S100A8/9 were commonly upregulated in both cDC1-and cDC2-derived LAMP3^+^DCs, while chemokines attracting Th2 cells/eosinophils (CCL17, CCL18), the IgE receptor (FCER1G), and the costimulatory molecule CD80 were uniquely upregulated in cDC2-derived LAMP3^+^DCs (Figure 3I). Interestingly, chemokines recruiting neutrophils (CXCL2, CXCL3, CXCL8), TNF, MMPs (MMP9, MMP19), IgG/E receptors (FCGR2A, FCER1A), and complements (C3AR1, C5AR1) were highly expressed in cDC1-and/or cDC2-derived LAMP3^+^DCs in non-lesional skin (Figure 3I). These findings indicate that cDC2-derived LAMP3^+^DCs exhibit more proinflammatory features in BP lesional skin, while LAMP3^+^DCs in non-lesional skin maintain a proinflammatory signature, suggesting an ongoing pathological process.

In BP patients, blood pDCs were decreased compared to healthy individuals, during both active and remission stages (Figure S2B and S2C). The DEgenes of blood pDCs in these stages exhibit a similar pattern compared to healthy individuals (Figure S2D). By analyzing the upregulated genes in top 10% of all DEgenes in pDCs from both stages revealed an 80% overlap in enriched pathways (Table S4), mainly involving TLR signaling (TLR8, CD14, ITGAM), IFN-γ signaling (FCGR1A, FCGR1B), antigen cross presentation (CD36, PSMB5), and neutrophil degranulation (C3AR1, S100A8, S100A9) (Figures S2E and S2F). Notably, CD14, CD36, S100A8, S100A9, FCGR1A showed even higher expression in pDCs from BP patients in remission stage compared to active stage, indicating a persistent proinflammatory signature in pDCs despite disease improvement (Figure S2D)

Four clusters of mast cells were identified in the skin of BP patients (Figures 4A and 4B). In general, genes associated with IL-2 signaling (IL2RA, IL2RG), type 2 inflammatory response (IL-4, IL-13, FCER1G), TNF signaling (TNFSF11, TNFRSF18), interferon signaling (IRF7, ISG15, IFITM3), leukocyte chemotaxis (CCL3, CCL4, CXCL8, CXCR4) and neutrophil degranulation (S100A8, S100A9), which are linked to the pathology of BP, were found to be upregulated in the lesional skin (Figures 4C and 4D). On the other hand, matrix metalloproteinases (MMP2, MMP14), which are crucial for blister formation^37^, AP-1 family members (JUN, FOS) that regulate mast cell cytokine production^38^, and IL1RL1 (ST2) signaling, which enhances the type 2 function of mast cells^39^, were upregulated in non-lesion skin (Figures 4C and 4D). These findings indicate a switching of inflammatory features of mast cells during the transition from non-lesional to lesional skin.

**Figure 4.**
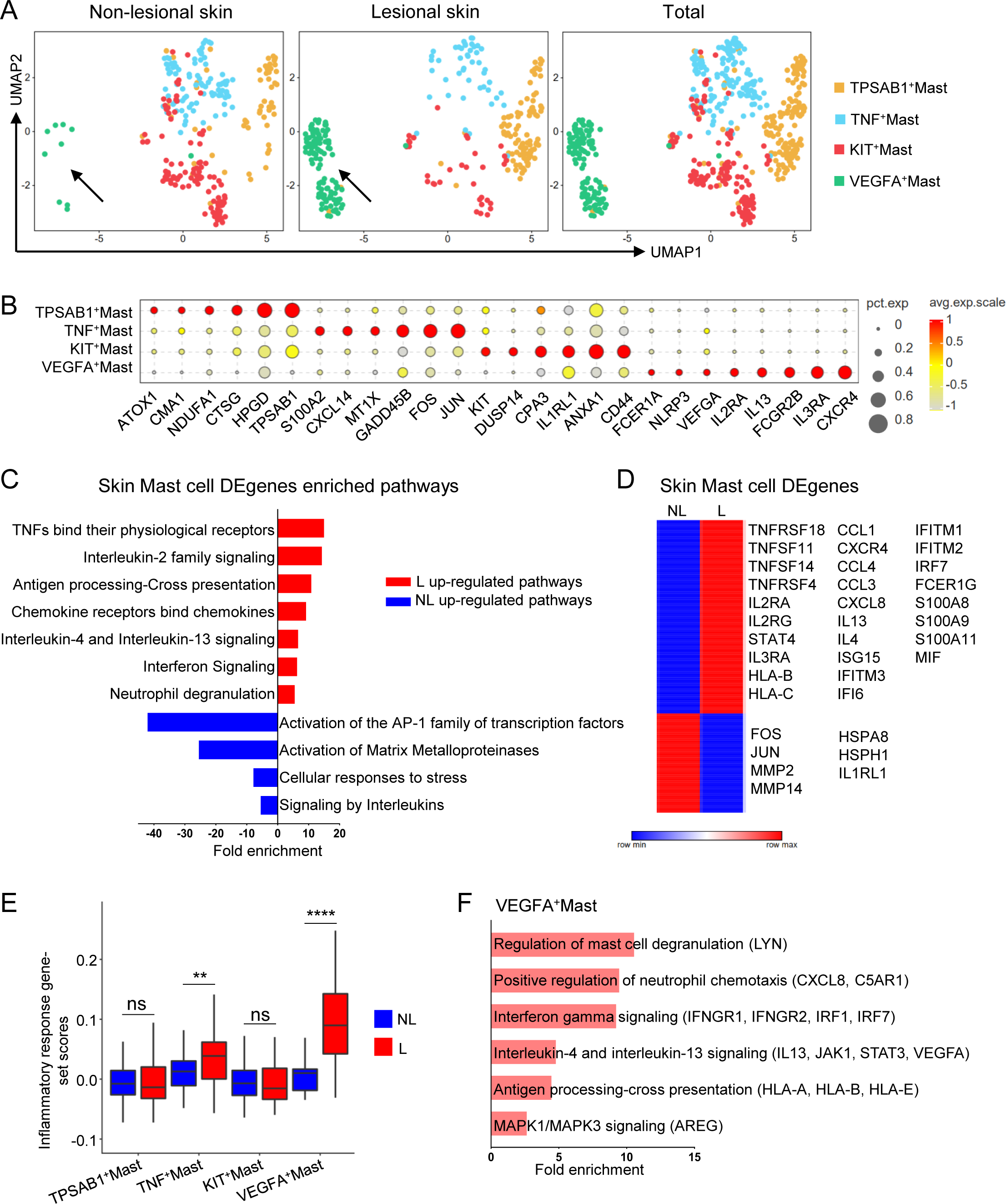
Characterization of mast cells in the skin of BP patients. (A) UMAP of non-lesional, lesional and total mast cells from the skin of BP patients. (B) Dot plot showing specific markers for mast cell clusters. (C) Enriched pathways of up-regulated (red) and down-regulated (blue) genes of mast cells in BP lesional skin by reactome analysis (FDR<0.05). (D) Heatmap of DEgenes between BP non-lesional and lesional skin mast cells. (E) Inflammatory response gene-set scores of mast cell clusters in BP non-lesional and lesional skin. Two-tailed paired t test, ns, not significant, **, p<0.01; ****, p<0.0001. Data are mean with s.e.m. (F) Enriched pathways of VEGFA^+^Mast cell highly expressed genes by reactome analysis (FDR<0.05).

TNF^+^ and VEGFA^+^ mast cells, linked to anti-tumor and pro-tumor activity respectively^33^, displayed heightened proinflammatory features in lesional skin (Figure 4E). TNF^+^ mast cells were mainly enriched with biological processes associated with translation (RPS15, RPS26, EEF1A1)) and cellular response to stress (HSPA1B, DNAJB1, FOS) (Figure S2G). Interestingly, VEGFA^+^ mast cells, particularly enriched with pathways driving BP progression, such as neutrophil chemotaxis (CXCL8, C5AR1), IL-4, IL-13, and IFN-γ signaling (IL13, JAK1, STAT3, IFNGR1, IFNGR2, VEGFA, IRF1, IRF7), and antigen presentation (HLA-A, HLA-B, HLA-E) (Figure 4F), were predominantly derived from lesional BP skin (Figure 4A), highlighting their critical roles in BP pathology.

### The BP related transcriptional features of B cells

Consistent with prior research, skin B cells were rare (Figure 1B)^40^, while blood B cells fell into four clusters: naive, TBX21^+^, IgM^+^ switched memory (SM), and SM (Figure S3A-S3B)^41,42^. We compared transcriptional features of B cells from healthy donors and BP patients in active and remission states (Figure S3C). B cells from BP patients commonly upregulated genes for NK and T cell interaction (CD48)^43^, negative feedback of B cell activation (CD52)^44^, and antigen presentation (HLA-A, HLA-B, HLA-DRB5). Active BP B cells upregulated genes for differentiation, memory, and preventing autoreactivity (CCR7, BACH2)^45,46^, while down regulating IRF7, critical for type 1 interferon induction (Figure S3C). In contrast, B cells in remission stage uniquely upregulated type 1 interferon signaling genes (BST2, S100A8, S100A9, TLR10, IRF1, STAT1)^47^, whereas genes involved in IL-4 response, IgM production, and cellular metabolism were downregulated (IL4R, IGHM, LAMTOR5) (Figure S3C). These findings delineate unique transcription profiles of B cells in BP patients, including shared upregulation of genes related to immune response, with active BP emphasizing B cell differentiation and memory, and remission showing heightened type 1 interferon signaling and reduced IL-4 response and IgM production.

### Blood LAG3^+^ and CX3CR1^+^ZNF683^+^ exhausted T cells represent BP reactive cells

It is well acknowledged that T cells play significant roles in BP progression through various mechanisms, including the production of itch mediators (IL-4, IL-13)^48^, recruitment of eosinophils, regulating of autoantibody production by B cells^49^, and modulation of inflammation and tissue damage^10^. During skin inflammation, blood T cells are a key source of the recruited T cells in the skin^50,51^. However, the detailed information regarding T cell subset clonal expansion, clone type sharing within the skin or between the skin and blood, which may have implications for potential therapeutic targets, remains unclear. To address these critical points, the transcriptome (including TCR) of paired skin and blood T cells from BP patients was examined (Figures 5A and S4A).

**Figure 5.**
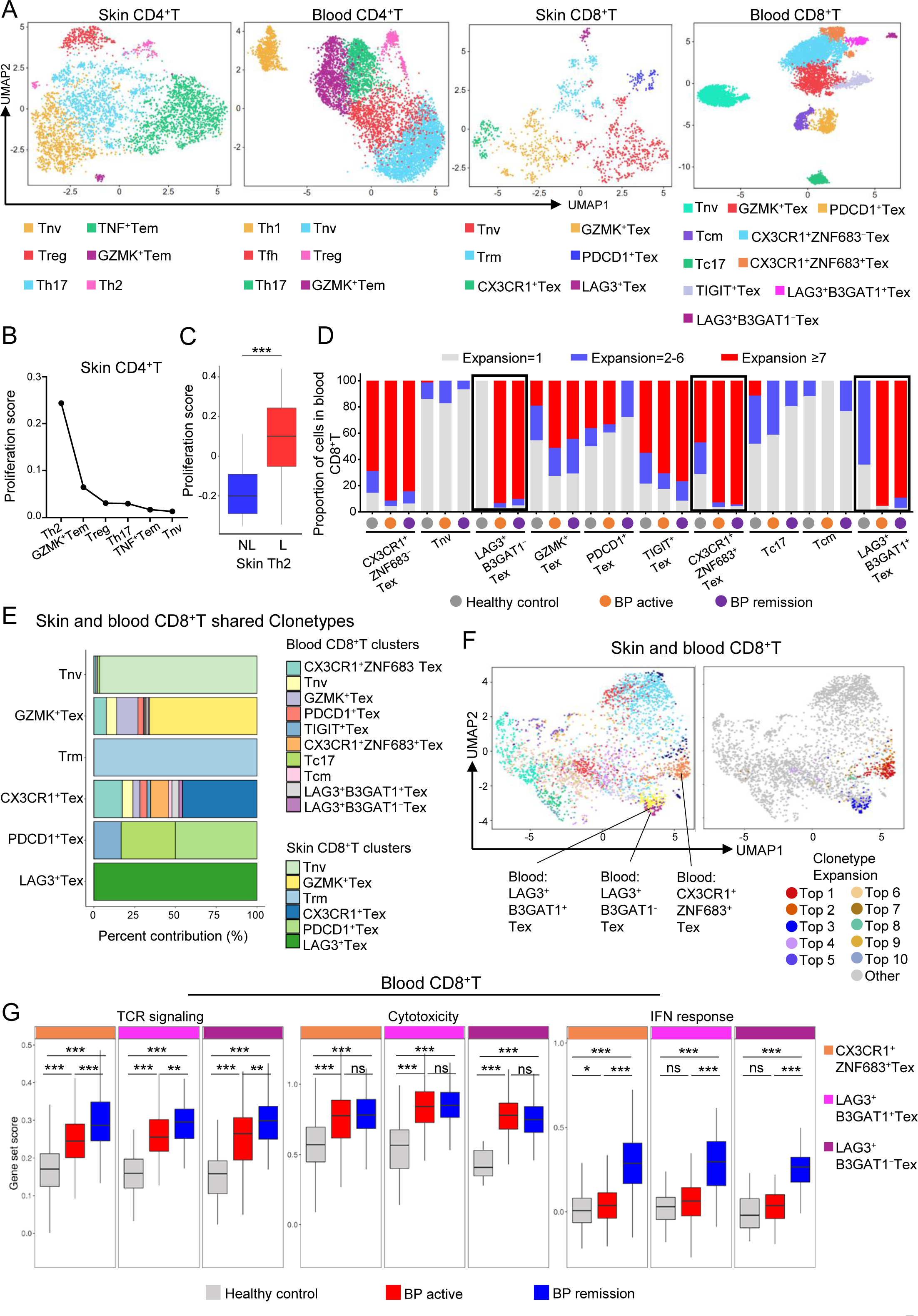
Blood LAG3^+^ and CX3CR1^+^ZNF683^+^ exhausted T cells represent BP reactive T cells. (A) UMAP of CD4^+^ and CD8^+^ T cells from BP non-lesional and lesional skin and from the blood of healthy donors and BP patients in active and remission stages. (B) The proliferation scores of BP skin CD4^+^T cell clusters. (C) Proliferation scores of Th2 cells in BP lesional and non-lesional skin. Two-tailed paired t test, ***, p<0.001. (D) Histogram showing the TCR expansion of blood CD8^+^T cell clusters in healthy donors and BP patients in active and remission stages. (E) Barplot showing the percentage of shared clone types between donor matched skin and blood CD8^+^T cells in active BP patients. (F) UMAP of mixed skin and blood CD8^+^T cells (left) and clonal expansion in shared clone types between skin and blood CD8^+^T cells (right) in active BP patients. (G) Gene set analysis of blood CX3CR1^+^ZNF683^+^ and LAG3^+^ Tex cells among healthy donors and BP patients in active and remission stages. Two-tailed unpaired t test, ns, not significant, *, p<0.05; **, p<0.01; ***, p<0.001. Data are mean with s.e.m.

Our results revealed that Th1 cells were exclusively detected in the blood, whereas Th2 cells were found only in the skin (Figures 5A and S4A). Among CD4^+^ T cells, Th2 cells exhibited the highest KI67 expression and proliferation score (Figures 5B and S4A), with higher scores in lesional skin than in non-lesional skin (Figure 5C), highlighting the pronounced activation of Th2 cells in the inflamed skin of BP patients. Except for Th1 cells, both skin and blood CD4^+^T cell clusters showed limited clonal expansion (Figures S4B-S4C), and inline with previous published reports^52^, the sharing of clone types within or between skin and blood CD4^+^T cell clusters was infrequent (Figures S5A-S5C).

In contrast to CD4^+^T cells, expanded TCRs were readily detected in blood and skin CD8^+^ T cell clusters, except for naive CD8^+^T cells in the blood (Figures 5D and S5D). LAG3^+^B3GAT1^+^, LAG3^+^B3GAT1^−^ and CX3CR1^+^ZNF683^+^ exhausted T cells in the blood exhibited higher TCR expansion in both active or recovered BP patients compared with healthy people (Figure 5D), whereas the TCR expansion in lesional and non-lesional skin CD8^+^T cells were largely comparable (Figures S5D). Shared clone types were more common in CD8^+^T cells than in CD4^+^T cells (Figures 5E, S5E and S5F). Moreover, we observed shared clone types between the paired skin and blood CD8^+^T cells in active BP patients. For example, 7.64%, 13.19% and 6.25% of clone types of skin GZMK^+^Tex cells were shared with blood CX3CR1^+^ZNF683^−^Tex, GZMK^+^Tex and Tnv cells, respectively (Figure 5E). Additionally, 10.87% and 17.39% of clone types of skin CX3CR1^+^Tex cells were shared with blood CX3CR1^+^ZNF683^+^ and CX3CR1^+^ZNF683^−^ exhausted T cell subsets, respectively (Figure 5E). Among these, blood CX3CR1^+^ZNF683^+^ and LAG3^+^ exhaust T cells exhibited the highest clonal expansion in clone types shared with skin CD8^+^T cells (Figure 5F), suggesting that these cells, with their increased fraction of expanded TCR clones in BP patients (Figure 5D), likely represent BP-reactive cells contributing to clonal expansion of skin CD8^+^T cell.

Gene set analysis revealed increased TCR activity and cytotoxicity scores in CX3CR1^+,^ZNF683^+^ and LAG3^+^ Tex cells from all BP patients (Figure 5G). Remission-stage patients exhibited higher TCR activity compared to those in the active stage or in healthy donors, while cytotoxicity scores were consistent across all BP patients (Figure 5G). A heightened IFN response was observed in both CX3CR1^+^ZNF683^+^ and LAG3^+^ Tex cells during remission, but only in CX3CR1^+^ZNF683^+^ Tex cells during the active stage (Figure 5G). These findings indicate that the features of BP-reactive CD8^+^ T cells are further influenced by the stage of BP.

### The activity of NK cells is associated with the disease progression of BP

Skin NK cells are primarily recruited from the circulation in response to inflammation and play a crucial role in the progression and resolution of skin diseases, including viral or bacterial infections and tumors^53,54^. NK cells from both lesional and non-lesional skin of BP patients, as well as from the blood of healthy individuals, BP patients in active and remission stages, were analyzed (Figure 6A). Unlike T cells, NK cells from the blood and skin were distinguished by their tissue origin (Figure 6B), confirming prior findings^53,54^.

**Figure 6.**
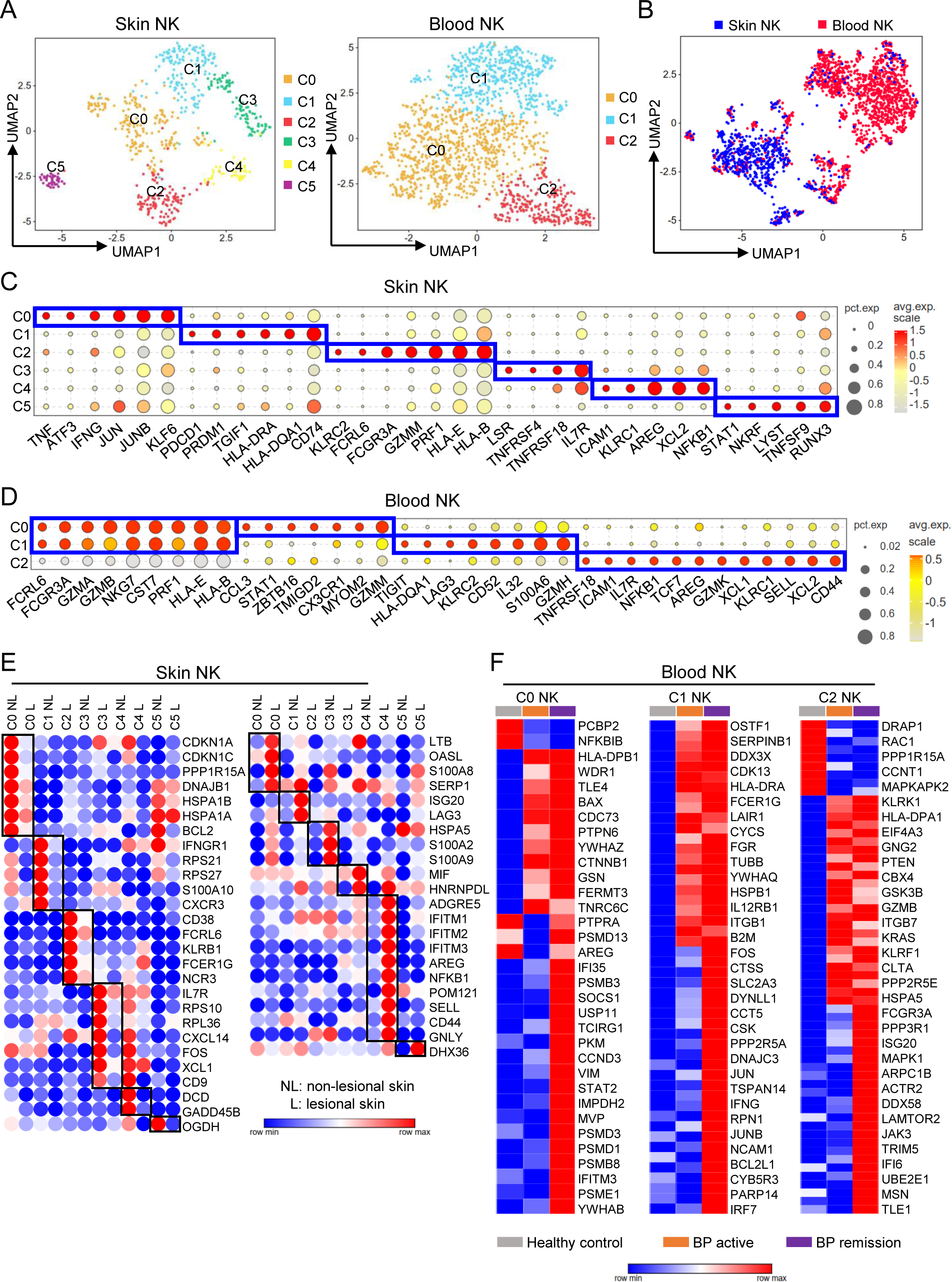
Characterization of NK cells in BP patients. (A) UMAP of NK cells from BP non-lesional and lesional skin (left) and from the blood of healthy donors and BP patients in active and remission stages (right). (B) UMAP of mixed skin (blue) and blood (red) NK cells. (C, D) Dot plots showing specific markers for skin (C) and blood (D) NK cell clusters. (E) Heatmap of cluster-specific DEgenes between BP non-lesional and lesional NK cells. (F) Heatmap of cluster-specific DEgenes among blood NK cell clusters of healthy donors and BP patients in active and remission stages.

Skin NK cells were categorized into six distinct clusters. Notably, Cluster 0 (C0) NK cells had high levels of inflammatory cytokines and AP-1 family members (TNF, IFNG, ATF3, JUN), and C2 NK cells decreased in lesional skin exhibited high expression of cytotoxicity-associated genes (FCGR3A, GZMM and PRF1), whereas C4 NK cells exhibited high expression of AREG, an EGFR ligand known to promote immune tolerance and tissue repair^25^, as well as inhibitory receptors KLRC1 (Figure 6C and S6A). Consistent with previous studies^24,55^, blood NK cells were predominantly clustered into CD56^dim^NK cells (C0 and C1) and CD56^hi^NK cells (C2), which also exhibit high expression of AREG (Figure 6D).

Transcriptional features in BP pathology in NK cells from skin and blood were analyzed. In lesional skin, cytotoxicity-associated genes were downregulated in C1 and C2 NK cells (IFNGR1, FCER1G, NCR3, KLRB1) (Figure 6E, left). Conversely, various interferon-stimulated genes involved in infection prevention, were upregulated in different NK cell clusters (OASL, ISG20, IFITMs, DHX36), and AREG, crucial for maintaining homeostasis in inflamed tissue^25^, was upregulated in C4 NK cells (Figure 6E, right), indicating a functional adaptation of NK cells in the lesional skin environment.

Compared with healthy individuals, the gene expression of blood NK cells affected by BP pathology were analyzed. Integrins (ITGB1, ITGB7), which promote epithelial residency, were upregulated in C1 and C2 NK cells during active and/or remission stages (Figure 6F). Cytotoxicity-associated genes (FCGR1G, IL12RB1, FCGR3A, GZMB, KLRF1, IFNG) were also upregulated in C1 and C2 NK cells during these stages, consistent with observations in BP-reactive CD8^+^ T cells (Figures 5G and 6F). However, ISGs were upregulated in C0 (IFI35, IFITM3), C1 (IRF7), and C2 (ISG20, DDX58, IFI6) NK cells in the remission stage (Figure 6F). Interestingly, AREG expression was downregulated in BP active patients but recovered in the remission stage in C0 NK cells, suggesting the involvement of NK cells in skin lesion repair (Figure 6F). Given that blood NK cells serve as the primary source of NK cells in inflamed skin, the interplay between the observed pro-inflammatory and homeostasis features of blood NK cells may collectively contribute to BP pathogenesis.

The flow cytometry was performed to detect the percentage, and the inflammatory (IFN-γ) and homeostatic (AREG) cytokines production of blood NK cells (Figure 7A). The results revealed a decrease in both the percentage and number of NK cells in individuals with active BP (Figures 7B and S6B). However, in recovered individuals, the amount of NK cells was restored to levels comparable to healthy controls (Figure 7B and S6B). The AREG produced by total NK cells was increased in all BP patients, while IFN-γ production was decreased in recovered patients (Figures 7C, 7D, S6C and S6D). NK cells were further categorized into AREG^+^IFN-γ^−^, AREG^+^IFN-γ^+^ and AREG^−^IFN-γ^+^ subsets based on the distinct function of the two cytokines. Our findings revealed that AREG^+^NK cells, including AREG^+^IFN-γ^−^ and AREG^+^IFN-γ^+^ NK cells, were elevated in all BP patients (Figures 7E, 7F, S6E and S6F), whereas IFN-γ^+^NK cells that did not produce AREG were reduced (Figure 7G and S6G).

**Figure 7.**
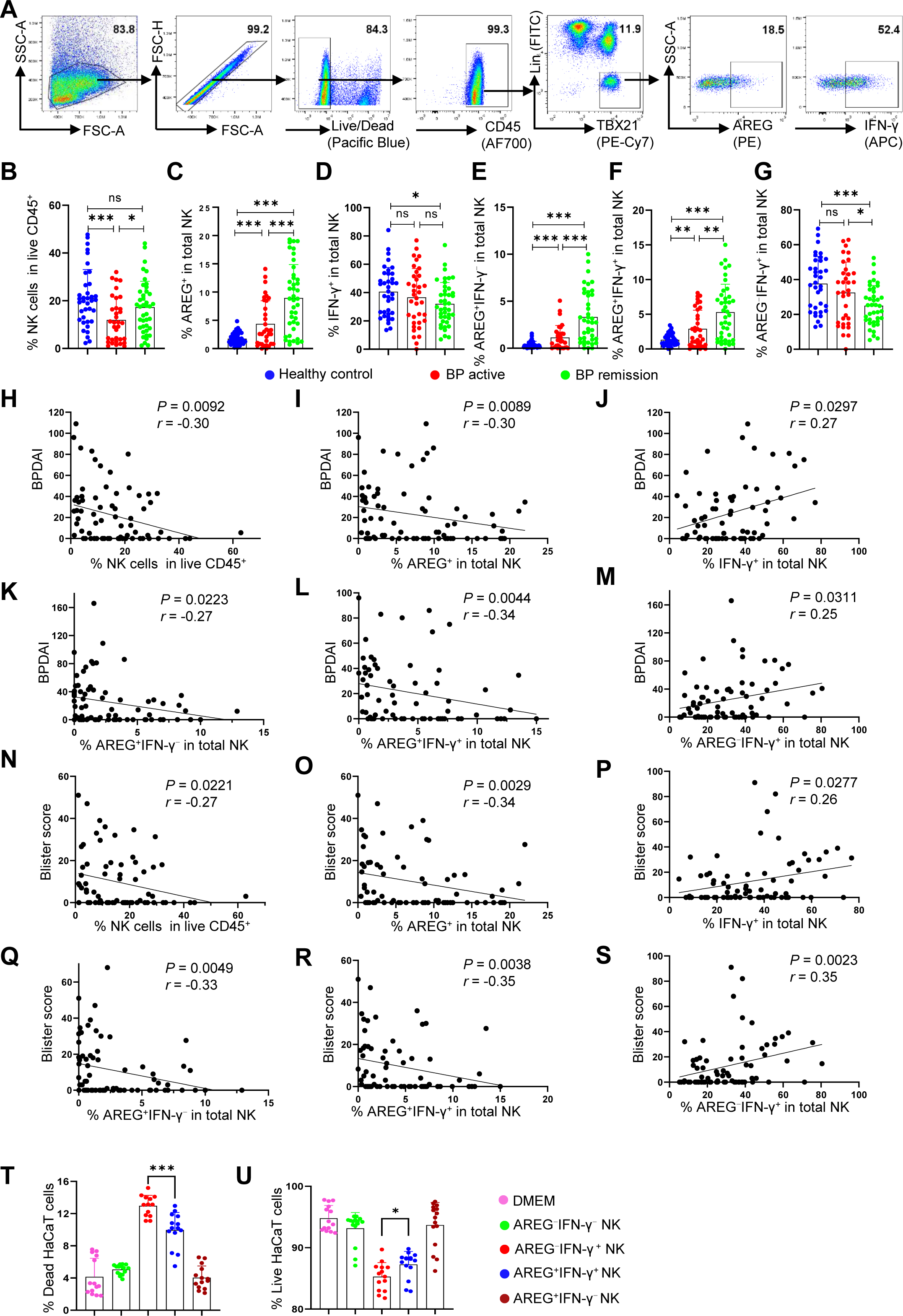
High AREG expression in BP blood NK cells correlates negatively with BP progression. (A) PBMCs were stimulated with IL-12 (10 ng/ml), IL-15 (50 ng/ml) and IL-18 (50 ng/ml) overnight, followed by incubation with protein transport inhibitor (1:1000 dilution) for 2 hrs in RPMI 1640 medium at 37°C with 5% CO_2_. The production of AREG and IFN-γ by NK cells (CD45^+^Lin^−^TBX21^+^) were detected by flow cytometry. (B-G) Histograms showing the percentage of NK cells (B), and AREG^+^ (C), IFN-γ^+^ (D), AREG^+^IFN-γ^−^ (E), AREG^+^IFN-γ^+^ (F) and AREG^−^IFN-γ^+^ (G) cells in total blood NK cells from healthy controls (n=37), BP patients in active (n=37) and remission (n=43) stages detected in (A). (H-M) Pearson correlation of the percentage of blood NK cells (H, n=74), AREG^+^ (I, n=75), IFN-γ^+^ (G, n=71), AREG^+^IFN-γ^−^ (K, n=72), AREG^+^IFN-γ^+^ (L, n=69) and AREG^−^IFN-γ^+^ (M, n=72) NK cells with BPDAI in BP patients. (N-S) Pearson correlation of the percentage of blood NK cells (N, n=72), AREG^+^ (O, n=73), IFN-γ^+^ (P, n=72), AREG^+^IFN-γ^−^ (Q, n=70), AREG^+^IFN-γ^+^ (R, n=68) and AREG^−^IFN-γ^+^ (S, n=72) NK cells with blister score in BP patients. (T-U) NK cells were enriched with magnetic beads and were cultured in NK cell MACS medium for 10 days to generate AREG^+^IFN-γ^−^ NK cells, or were further stimulated with IL-12 (10 ng/ml), IL-15 (50 ng/ml) and IL-18 (50 ng/ml) for 16 hrs to generate AREG^+^IFN-γ^+^ NK cells. The expanded NK cells were rested in MACS medium without supplement for 3 days to generate AREG^−^IFN-γ^−^NK cells, or were further stimulated with IL-12, IL-15 and IL-18 for 16 hrs to generate AREG^−^IFN-γ^+^ NK cells. All NK cells were then washed for 3 times, and were cultured in DMEM for 48 hrs to prepare the conditioned medium. HaCaT cells were cultured in conditioned medium for 48 hours, digested with TrypLE, and subjected to live/dead staining. Each dot represents a unique donor, For (B-G, T and U), two-tailed unpaired t test, ns, not significant, *, p<0.05; **, p<0.01; ***, p<0.001. Data are mean with s.e.m.

The correlation between NK cell cytokine production profiles and BP clinical indicators, including bullous pemphigoid disease area index (BPDAI) and blister score^56^, was examined. The percentage of total NK cells and AREG^+^ NK cells showed a negative correlation with BPDAI, while IFN-γ^+^NK cells exhibited a positive correlation with BP disease severity (Figures 7H-7J). Additionally, both subsets of AREG^+^NK cell, including the percentage of AREG^+^IFN-γ^−^ and AREG^+^IFN-γ^+^ NK cells were negatively correlated with BPDAI, whereas the percentage of AREG^−^IFN-γ^+^ NK cells were positively correlated with BPDAI (Figures 7K-7M), suggesting that the IFN-γ and AREG produced by NK cells may play opposite roles in BP disease progression.

The pathology of BP primarily affects keratinocytes in the basal layer of the epidermis^1,2^, where these cells are targeted by autoantibodies and further impacted by the inflammatory environment^1^. Cell chat analysis revealed that keratinocytes show the strongest ligand-receptor interactions with NK cells among all leukocytes (Figure S7A). The observation that IFNG-IFNGR1/2 signaling was enhanced, while AREG-EGFR signaling was decreased between NK cells and keratinocytes in BP lesional skin (Figure S7B), coupled with correlation analysis results (Figures 7H-7S), prompted us to investigate the biological impact of NK cell-produced AREG and IFN-γ on keratinocytes.

The survival of HaCaT epidermal keratinocytes cultured with supernatants from different NK cell cultures revealed significant differences (Figures 7T, 7U and S7C). Specifically, the IFN-γ from AREG^−^IFN-γ^+^NK cell culture supernatant induced notable cell death compared to DMEM alone or supernatant from AREG^−^IFN-γ^−^NK cell culture (Figure 7T). Conversely, HaCaT cells showed increased survival in AREG^+^IFN-γ^+^ than in AREG^−^IFN-γ^+^NK cell conditioned medium (Figure 7U), suggesting that AREG counteracts IFN-γ-induced apoptosis. The live or dead HaCaT cells percentage in AREG^+^IFN-γ^−^ conditioned medium was similar to DMEM alone (Figures 7T and 7U), indicating that the presence of AREG alone did not affect HaCaT cell survival. These experiments suggest AREG mitigates IFN-γ-induced apoptosis of keratinocytes during BP progression, potentially elucidating the negative correlation between AREG^+^NK cells and the severity of the disease.

## Discussion

Accelerating the wound healing of BP lesional skin is crucial for reducing the risk of pathogen infection and inflammatory irritants. This process requires complex collaboration among skin structural cells. Beyond the unique features impacted by BP skin pathology, we observed that skin non-immune cells exhibited a coordinated response to BP-induced damage, with altered gene expression across different stages of wound healing (Figures 2C-2E). This may be regulated by their shared enriched pathways in the lesional skin, such as upregulated glycolysis, oxidative phosphorylation, and response to stress. Targeting these shared pathways may universally impact various types of skin structural cells to facilitate the healing of skin lesions. Supporting this notion, it has been reported that inhibiting glycolysis impairs wound healing, while high glucose treatment accelerates re-epithelialization in vitro ^57^. Therefore, BP patients with metabolic disorders experiencing delayed healing of lesional skin may be further affected by additional metabolic inhibitor treatment.

The elevated activity of type 1 interferon signaling is closely associated with the susceptibility to autoimmune diseases such as systemic lupus erythematosus, Sjögren’s syndrome, and rheumatoid arthritis ^58^. Counterintuitively, we consistently observed an upregulation of type 1 interferon-associated genes in multiple blood cell populations of BP patients in the remission stage, including pDCs, B cells, CD8^+^T cells, and NK cells (Figures S2D, S3C, 5G, and 6F). However, type 1 interferon also has an anti-inflammatory mechanism due to its role in inducing immunosuppressive factors such as IL-10, SOCS1, and TTP, as well as inhibiting the expression of MMP genes^59,60^, which would alleviate inflammation and facilitate disease improvement. Thus, the biological significance of this observation on BP disease progression warrants further research.

We have uncovered changes in the composition and function of several cell populations associated with BP pathology. For instance, immunosuppressive MARCO^+^ macrophages were decreased, while LAMP3^+^ DCs and VEGFA^+^ mast cells, which are enriched with genes promoting Th2 responses and attracting eosinophils, were increased in the lesional skin (Figures 3F, 3G, and 4F). Identifying interventions that can reverse these alterations may be beneficial for alleviating BP-induced skin symptoms. Additionally, our analysis revealed that LAG3^+^ and CX3CR1^+^ZNF683^+^ exhausted T cells in the blood share the highest expanded TCRs with skin CD8^+^T cells. These cells also exhibited elevated TCR signaling and cytotoxicity scores in BP patients compared to healthy controls (Figures 5E, 5F, and 5G). Understanding their biological roles may provide new insights into the mechanisms of T cells in BP pathogenesis.

Traditionally, NK cells are recognized as proinflammatory cells that mediate tissue damage by producing IFN-γ and TNF-α^55^. However, our recent research revealed that, unlike mouse NK cells, human NK cells also serve as a major source of AREG, an EGFR ligand that plays critical roles in maintaining tissue homeostasis and immune tolerance^24^. Interestingly, clinical parameters measuring BP severity negatively correlated with NK cell AREG production but positively correlated with their IFN-γ production, suggesting opposite functions of NK cell produced IFN-γ and AREG in BP disease progression (Figures 7H-7S). This observation is consistent with the increased AREG^+^IFN-γ^−^ NK cells and decreased AREG^−^IFN-γ^+^ NK cells in remission BP patients compared to active BP patients (Figures 7E-7G). In vitro experiments further confirmed that NK cell-produced AREG counteracts IFN-γ-induced apoptosis of epithelial cells (Figures 7T and 7U). Thus, inhibiting NK cell IFN-γ production and enhancing their AREG production could be a promising therapeutic strategy for BP.

## Acknowledgments

This study was supported by CAMS Innovation Fund for Medical Sciences (CIFMS) (2024-I2M-3-005) and Hospital for Skin Diseases, Institute of Dermatology, Chinese Academy of Medical Sciences and Peking Union Medical College grant (3301030103119) to Yetao Wang. Scientific Research Project of Jiangsu Provincial Health Commission (ZD2021035) and Clinical and Translational Medicine Research Project of Chinese Academy of Medical Sciences (2023-I2M-C&T-B-112) to Suying Feng. We thank the study participants who provided skin and blood samples. All staff in the biobank of Institute of Dermatology, Chinese Academy of Medical Sciences, Jiangsu Biobank of Clinical Resources assisted with clinical sample collection. Guangzhou Genedenovo Biotechnology Co., Ltd assisted with sequencing data analysis.

## Author contributions

Yetao Wang: Conceptualization; experiment design; data analysis; resources; supervision; validation; investigation; visualization; methodology; data curation; software; writing − original draft, review and editing; project administration; funding acquisition. Suying Feng: experiment design; resources; review and editing; Guirong Liang: experiment design, performing experiments; data analysis; validation; visualization, data curation; review and editing. Chenjing Zhao: data analysis; validation. Qin Wei: performing experiments.

---

Methods.

| REAGENT or RESOURCE | SOURCE | IDENTIFIER |
| --- | --- | --- |
| **Antibodies** |  |  |
| Anti-Human CD3 | Biologend | Cat# 317306 |
| Anti-Human CD4 | Biologend | Cat# 317408 |
| Anti-Human TCR$\alpha/\beta$ | Biologend | Cat# 306706 |
| Anti-Human TCR$\gamma/\delta$ | Biologend | Cat# 331208 |
| Anti-Human CD19 | Biologend | Cat# 302206 |
| Anti-Human CD20 | Biologend | Cat# 302304 |
| Anti-Human CD22 | Biologend | Cat# 363508 |
| Anti-Human CD14 | Biologend | Cat# 325604 |
| Anti-Human CD34 | ebioscience | Cat# 11-0349-42 |
| Anti-Human FcεR1α | ebioscience | Cat# 11-5899-42 |
| Anti-Human CD1a | Biologend | Cat# 300104 |
| Anti-Human CD11c | Biologend | Cat# 301604 |
| Anti-Human CD123 | Biologend | Cat# 306014 |
| Anti-Human BDCA2 | Biologend | Cat# 354208 |
| Anti-Human TBX21 | ebioscience | Cat# 25-5825-82 |
| Anti-Human IFN-γ | ebioscience | Cat# 17-7319-82 |
| Anti-Human AREG | ebioscience | Cat# 12-5370-42 |
| Anti-Human CD45 | ebioscience | Cat# 56-9459-42 |
| **Biological Samples** |  |  |
| Skins | Biobank of Institute of Dermatology, Chinese Academy of Medical Sciences, Jiangsu Biobank of Clinical Resources (BM2015004) |  |
| PBMCs | Biobank of Institute of Dermatology, Chinese Academy of Medical Sciences, Jiangsu Biobank of Clinical Resources (BM2015004) |  |
| Blood for NK cells purification | Jiangsu province blood center | https://www.jsblood.com.cn/ |

| **Chemicals, Peptides, and Recombinant Proteins** |  |  |
| --- | --- | --- |
| IL-2 | Novoprotein | Cat# C013 |
| IL-12 | Novoprotein | Cat# CI58 |
| IL-15 | Novoprotein | Cat# C016 |
| IL-18 | Novoprotein | Cat# CH29 |
| Protein transport inhibitor | ebioscience | Cat# 00-4980-03 |
| TRIzol reagent | Invitrogen | Cat# 15596026CN |
| Fixable Viability Dye efluor 780 | Invitrogen | Cat# 65-0865-18 |
| **Critical Commercial Assays** |  |  |
| EasySep Human NK Cell Isolation Kit | STEMCELL | Cat# 17955 |
| NK MACS Medium human | MACS | Cat# 130-114-429 |
| Chromium NextGEM Single Cell 5' HT Kit v2 | 10 X Genomics | Cat# PN-1000356 |
| NEBNext Ultra RNA Library Prep Kit for Illumina | New England Biolabs | Cat# 7530 |
| **Deposited Data** |  |  |
|  | NCBI Sequence Read Archive |  |
| **Software and Algorithms** |  |  |
| FlowJo | FlowJo, LLC | https://www.flowjo.com/ |
| GraphPad Prism | GraphPad Software, Inc | https://www.graphpad.com/scientific-software/prism/ |
| Cell Ranger | 10x Genomics | https://support.10xgenomics.com/single-cell-gene-expression/software/pipelines/latest/what-is-cell-ranger |
| R Statistical Computing Software | The R Foundation | https://www.r-project.org/ |
| GSVA | Hänzelmann, S. et al.<sup>61</sup> | https://www.bioconductor.org/packages/release/bioc/html/GSVA.html |
| Seurat 3.0 | Stuart T. et al.<sup>62</sup> | http://satijalab.org/seurat/ |

## Resource availability

For additional details and requests regarding resources and reagents, please contact the lead person, Yetao Wang.

## Materials availability

This study did not produce novel or distinctive reagents.

## Data availability

scRNA-Seq data of non-lesional and lesional skin of bullous pemphigoid (BP) and PBMCs from healthy controls and BP patients, and bulk RNA-Seq data of PBMCs from healthy controls and BP patients, are available in the NCBI Sequence Read Archive (SRA), BioProject: PRJNA1124631

## Clinical samples

Samples of both non-lesional and lesional skin from BP patients, as well as PBMCs from both BP patients and healthy controls, were obtained from the biobank of the Institute of Dermatology, Chinese Academy of Medical Sciences, Jiangsu Biobank of Clinical Resources (BM2015004). Diagnostic criteria of BP: (1) intensely pruritic blisters of the skin, (2) positive linear IgG and/or C3c staining along the basement membrane zone by direct immunofluorescence, (3) positive IgG staining on the epidermal side of 1M-NaCl-split normal human skin by indirect immunofluorescence, (4) positive IgG enzyme-linked immunosorbent assay of BP180 NC16A and/or BP230 (MBL, Nagoya, Japan, >9 U/ml). BP was diagnosed if the first criterion and at least two of the last three criteria were met. Healthy blood samples used for NK cell purification were obtained from the Jiangsu Province Blood Center. All participants provided written informed consent following approval by the ethics committee of the Institute of Dermatology, Chinese Academy of Medical Sciences and Peking Union Medical College. Detailed clinical characteristics for each BP patient and healthy control are provided in Table S1.

## Human skin cell preparation

To isolate single cells from both non-lesional and lesional skin of BP patients for scRNA-Seq, skin biopsies were processed using the instructions provided with the whole skin dissociation kit (130-101-540, MACS). Initially, fat tissue was removed by scraping, and the biopsies were then cut into small pieces (< 4 mm^2^). Each piece underwent digestion in a buffer containing 435 μl buffer L, 12.5 μl enzyme P, 50 μl enzyme D, and 2.5 μl enzyme A on a 37°C shaker for 3 hrs. The digested skin tissue was subsequently diluted in ice-cold DMEM containing 2% BSA, filtered through a 70 μm strainer, and the flow-through was centrifuged at 300 x g for 10 min at 4°C. The isolated skin cells were finally resuspended in a MACS buffer (0.5% BSA + 2 mmol EDTA+1L PBS).

## Peripheral blood mononuclear cells (PBMCs) isolation

Each human peripheral blood leukopak was initially washed with 80 ml of serum-free RPMI 1640 (Gibco), followed by layering onto lymphoprep (STEMCELL, 07851) and centrifugation at 500 x g for 30 min at room temperature. After three washes with MACS buffer, the isolated PBMCs were either used immediately or cryopreserved in FBS containing 10% DMSO.

## Flow cytometry

PBMCs were initially stained with fixable viability dye efluor 780 (Invitrogen, 65-0865-18). For surface staining, cells were then incubated with antibodies in MACS buffer at 4°C for 30 minutes in the dark. For intracellular staining, cells were fixed and permeabilized using the Foxp3 staining kit (eBioscience, 00-5523-00), followed by staining for cytokines or TBX21 with antibodies in permeabilization buffer at 4°C for 30 minutes in the dark. The cells were ready for flow cytometry analysis after washing with MACS buffer.

## scRNA-Seq library preparation

For scRNA-Seq library preparation, we utilized the Chromium NextGEM Single Cell 5’ HT Kit v2 (10x Genomics, PN-1000356). Skin cells and PBMCs were washed and resuspended in 1x PBS containing 0.05% BSA. Cell count and viability were assessed using trypan blue staining under a microscope, with cell concentration adjusted to 1,000-2,000 cells/μl (viability > 80%). The single-cell suspension was loaded onto the Chromium Controller (10x Genomics) to encapsulate single cells into gel beads in emulsion (GEMs). The quality of amplified cDNA and the final sequencing library were assessed using the Agilent 2100 Expert (Agilent Technologies). Sequencing depth was maintained at approximately 30,000 mean reads per cell for skin cells and 70,000 mean reads per cell for PBMCs (Table S2). The libraries were sequenced using the Illumina NovaSeq 6000 platform.

## scRNA-Seq data processing

113,044 skin cells and 38,854 PBMCs were identified by Cell Ranger (10x Genomics, version 5.0.0) with an average depth of 45,095 mean reads/cell (Table S2). A total of 95,527 cells (for skin) and 33,921 cells (for PBMCs) passed the filtering criteria using Seurat (Version 3.0) and were subsequently used for downstream analysis. The filtering criteria included: (1) ensuring nFeature ranged from 200 to 5900; (2) restricting UMIs to less than 49,000; and (3) ensuring UMIs from mitochondrial genes constituted less than 35% of the total UMIs. Cells derived from the skin were re-clustered into 12 unique clusters (res=0.5) and PBMCs derived from healthy donors and BP patients were mainly re-clustered into 4 clusters (res=0.5) for further analysis.

To identify genes highly expressed in a specific cluster, we compared their expression values with those of other clusters using the FindMarkers function in Seurat, employing the Model-based Analysis of Single-cell Transcriptomics (MAST) test. The average expression counts for each cluster were computed using the AverageExpression function.

## Gene sets score analysis

The gene set scores for cell clusters were calculated using the AddModuleScore function in Seurat, with significance determined by either a Wilcoxon rank-sum test or a two-tailed unpaired t-test. Table S5 listed the signature genes for gene set score analysis.

## Pseudotime analysis of BP skin DC subsets

To investigate the developmental pathways of LAMP3^+^ DCs, we utilized the Monocle algorithm (version 2.14.0) and focused on the top 400 signature genes identified by the differentialGeneTest function, The differentiation trajectory of DCs was inferred using Monocle with default parameters following dimension reduction and cell ordering.

## Bulk RNA-Seq library preparation

Total RNA was isolated using the Trizol reagent kit (Invitrogen, Carlsbad, CA, USA) following the manufacturer’s instructions. RNA integrity was evaluated with an Agilent 2100 Bioanalyzer and confirmed by RNase-free agarose gel electrophoresis. Subsequently, eukaryotic mRNA was enriched using Oligo(dT) beads. The enriched mRNA was then fragmented using a fragmentation buffer and reverse transcribed into cDNA using the NEBNext Ultra RNA Library Prep Kit for Illumina (NEB #7530, New England Biolabs, Ipswich, MA, USA). The resulting double-stranded cDNA fragments underwent end repair, addition of an A base, and ligation to Illumina sequencing adapters. After purification of the ligation reaction using AMPure XP Beads (1.0X), the cDNA fragments were PCR amplified. The prepared cDNA library was sequenced using Illumina Novaseq6000.

---

**Figure S1.**
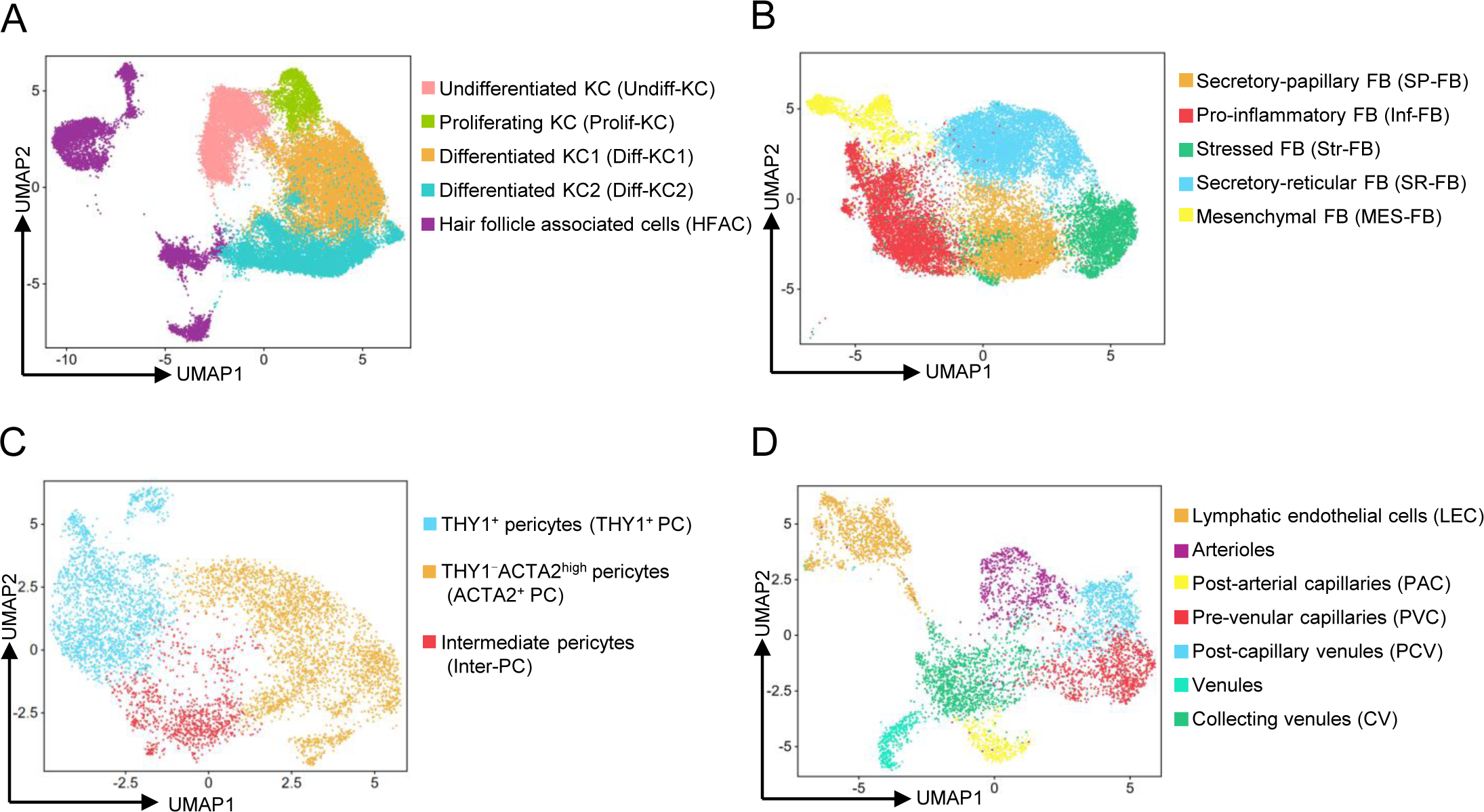
UMAP of skin non-immune cells of BP patients. (A-D) UMAP of keratinocytes (A), fibroblasts (B), pericytes (C) and endothelial cells (D) from BP non-lesional and lesional skin.

**Figure S2.**
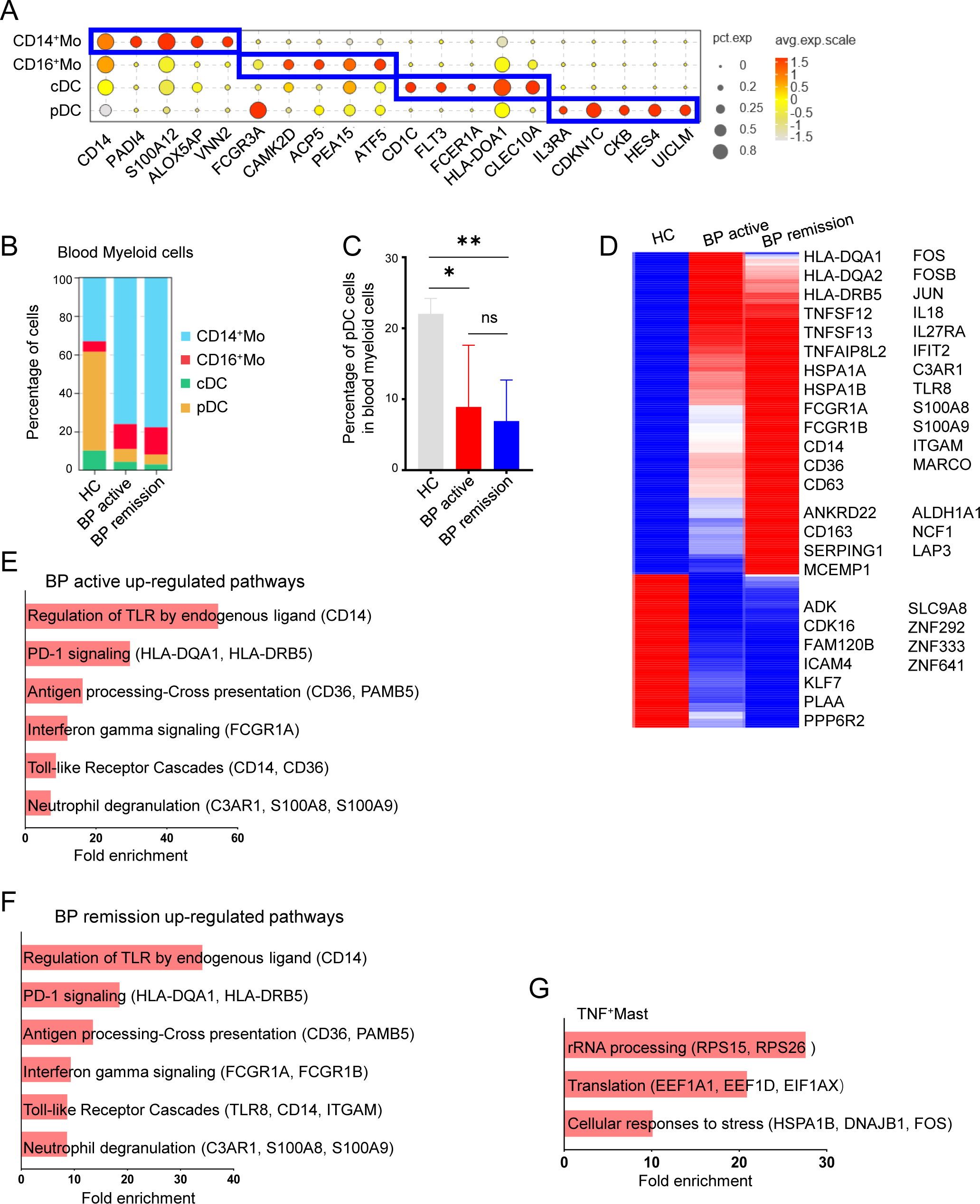
Characterization of mononuclear phagocytes in BP patients. (A) Dot plot showing specific markers for blood myeloid cell clusters from healthy donors and BP patients in active and remission stages. (B) Histogram showing the percentage of blood myeloid cell clusters from healthy donors (n=2) and BP patients in active (n=2) and remission (n=2) stages. (C) Percentage of blood pDC cells from healthy donors (n=5) and BP patients in active (n=5) and remission (n=5) stages obtained from bulk RNA-seq data using CYBERSORTx. Mann-Whitney test, ns, not significant, *, p<0.05; **, p<0.01. Data are mean with s.e.m. (D) Heatmap of DEgenes of blood pDCs among healthy donors and BP patients in active and remission stages. (E, F) Enriched pathways of up-regulated genes in top 10% of all DEgenes by reactome analysis (FDR<0.05) in blood pDCs from BP patients in active (E) or remission (F) stages vs healthy donors. (G) Enriched pathways of TNF^+^Mast cell highly expressed genes by reactome analysis (FDR<0.05).

**Figure S3.**
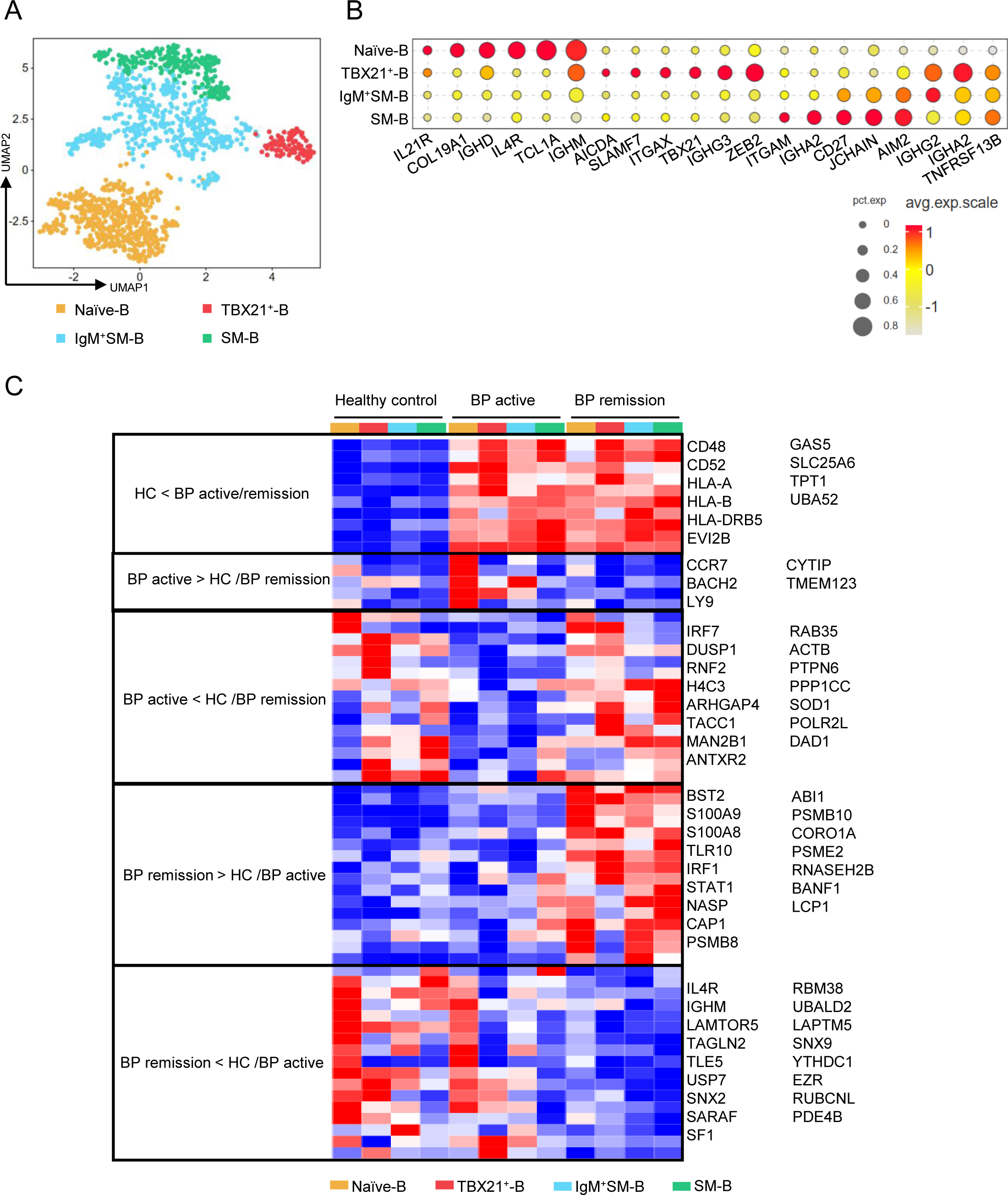
The BP related transcriptional features of B cells. (A) UMAP of B cells from the blood of healthy donors and BP patients in active and remission stages. (B) Dot plot showing specific markers for blood B cell clusters. (C) Heatmap of DEgenes of blood B cell clusters among healthy donors and BP patients in active and remission stages.

**Figure S4.**
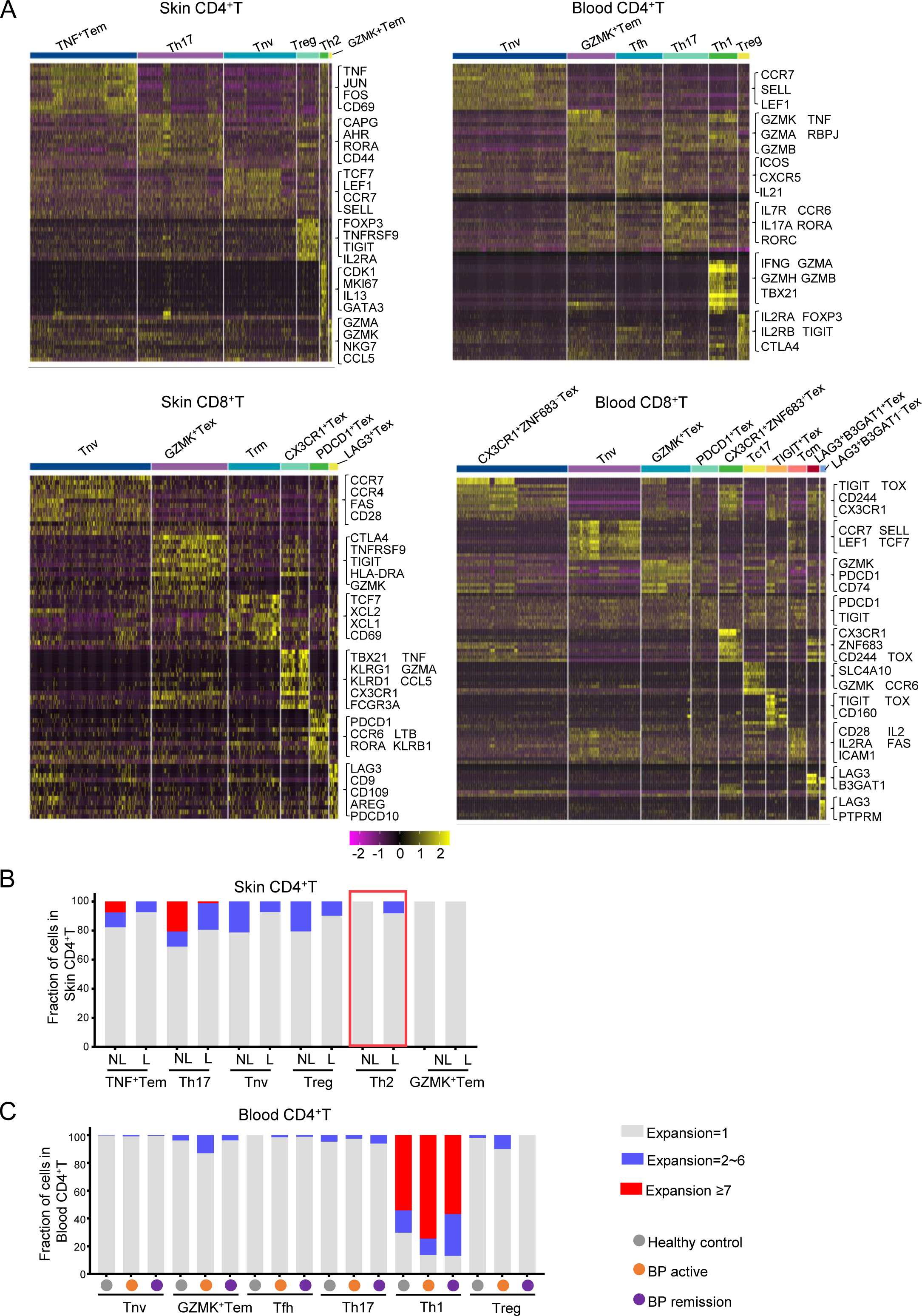
Characterization of T cells in BP patients. (A) Heatmaps showing specific markers for CD4^+^ and CD8^+^ T cell clusters from BP non-lesional and lesional skin and from the blood of healthy donors and BP patients in active and remission stages. (B, C) Histogram showing the TCR expansion of CD4^+^T cell clusters in BP non-lesional and lesional skin (B), and in the blood of healthy donors and BP patients in active and remission stages (C).

**Figure S5.**
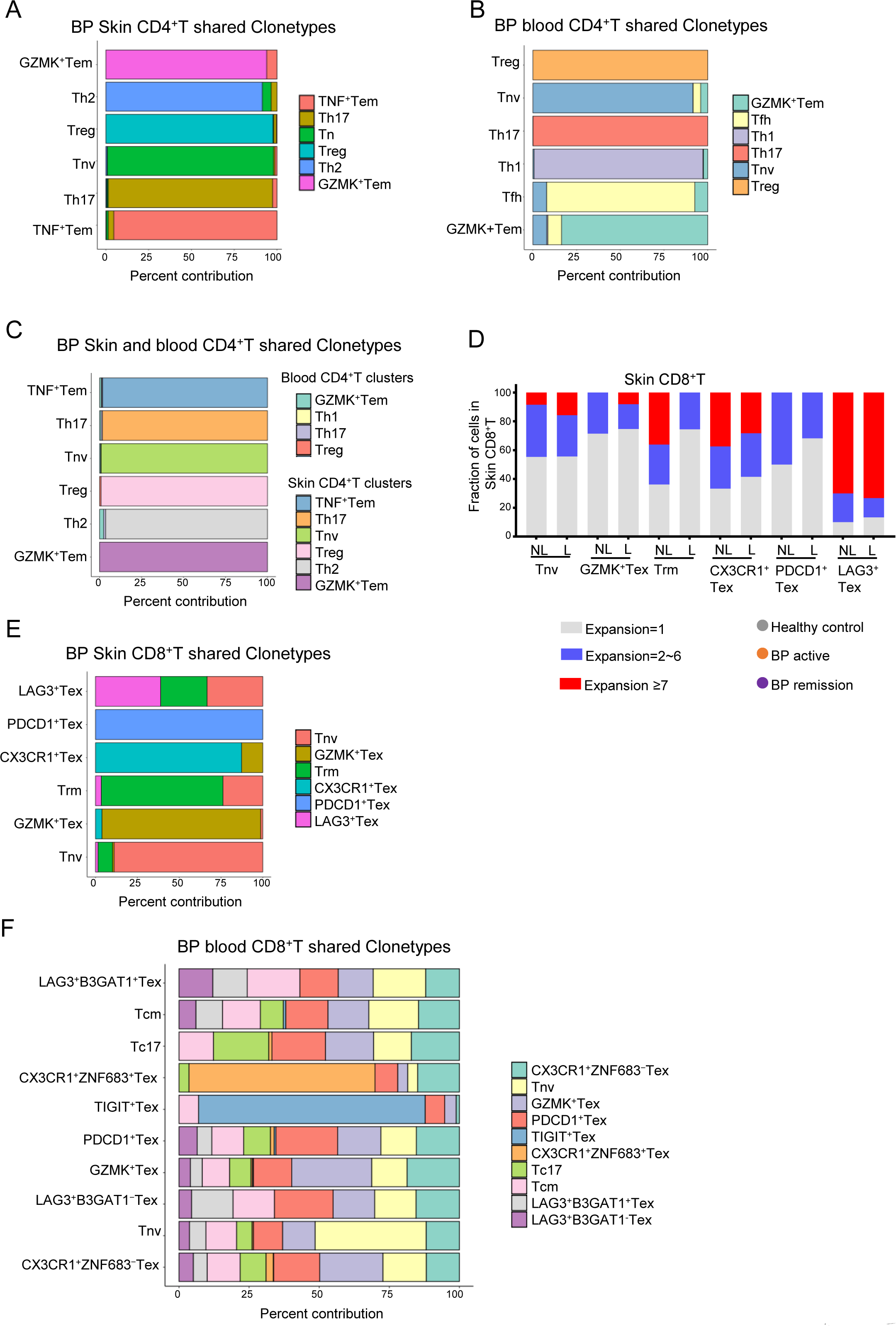
Characterization of T cells in BP patients. (A-C) Barplot showing the percentage of shared clone types in CD4^+^T cell clusters from BP non-lesional and lesional skin (A), blood of healthy donors and BP patients in active and remission stages (B), and between donor matched skin and blood of active BP patients (C). (D) Histogram showing the TCR expansion of CD8^+^T cell clusters in BP non-lesional and lesional skin. (E, F) Barplot showing the percentage of shared clone types in CD8^+^T cell clusters from BP non-lesional and lesional skin (E), and blood of healthy controls and BP patients in active and remission stages (F).

**Figure S6.**
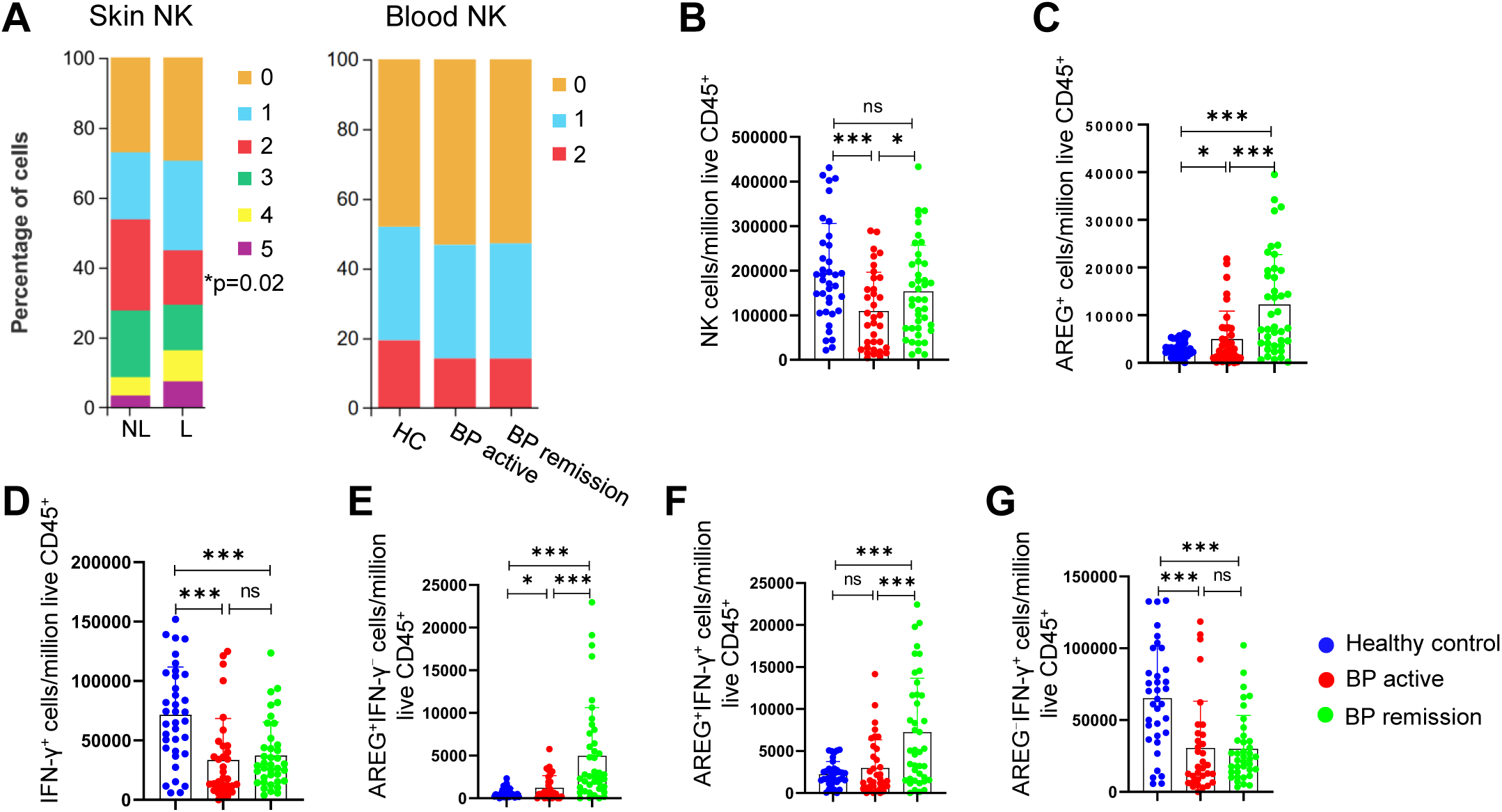
High AREG expression in BP blood NK cells correlates negatively with BP progression. (A) Histograms showing the percentage of NK cell clusters from BP non-lesional and lesional skin (left, n=4), and from the blood of healthy donors (n=2) and BP patients in active (n=2) and remission stages (n=2) (right). Mann-Whitney test, *, p<0.05. (B-G) Histograms showing the number of NK cells (B), and AREG^+^ (C), IFN-γ^+^ (D), AREG^+^IFN-γ^−^ (E), AREG^+^IFN-γ^+^ (F) and AREG^−^IFN-γ^+^ (G) NK cells in one million live CD45^+^ blood cells from healthy donors (n=37), BP patients in active (n=37) and remission (n=43) stages detected in (Figure 7A). Each dot represents a unique donor, two-tailed unpaired t test, ns, not significant, *, p<0.05; **, p<0.01; ***, p<0.001. Data are mean with s.e.m.

**Figure S7.**
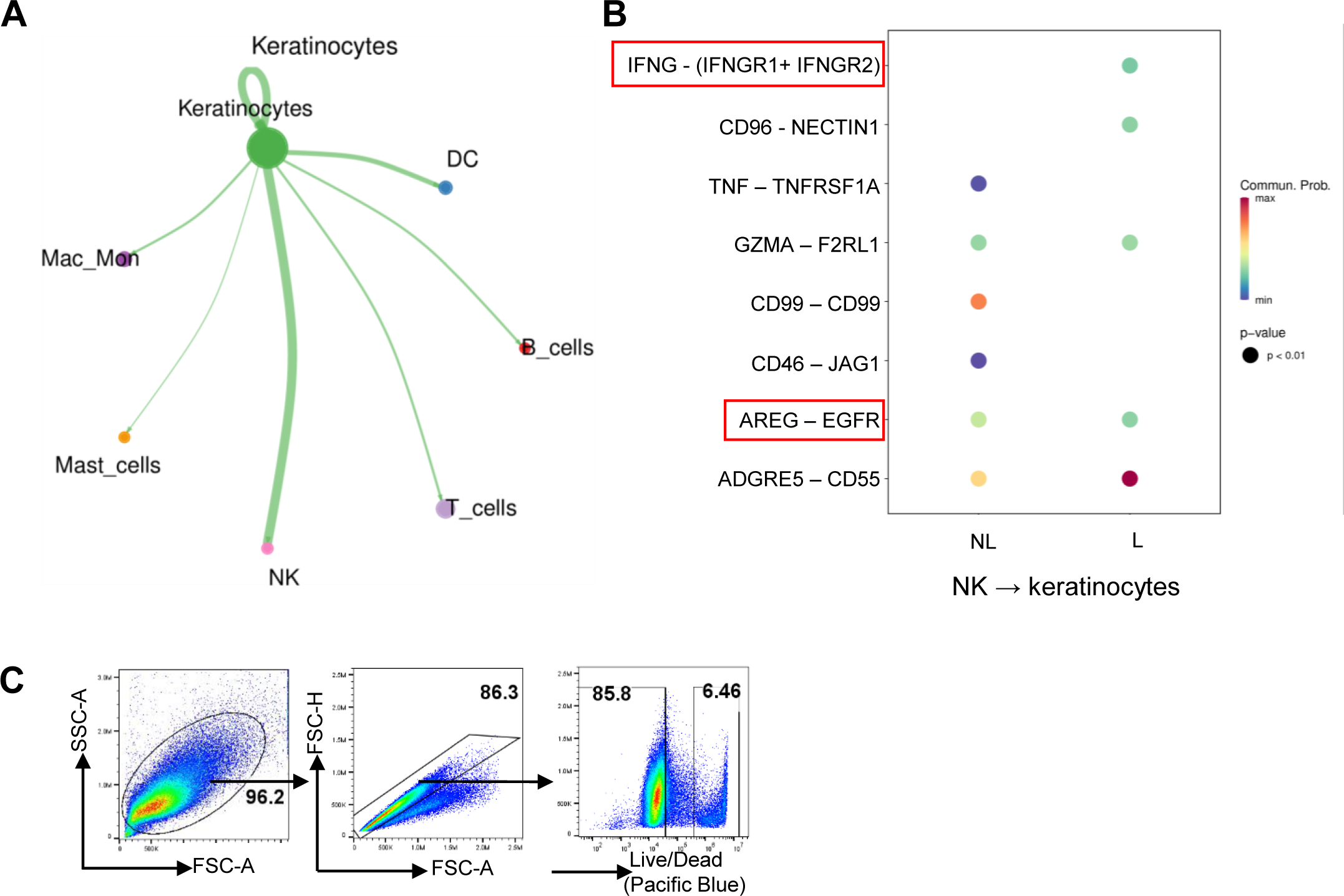
High AREG expression in BP blood NK cells correlates negatively with BP progression. (A) Network diagram of numbers of receptor-ligand pairs between keratinocytes and other immune cells in non-lesion and lesion of BP patients. The size of the peripheral circles indicates the number of receptor pairs contained within each subpopulation, the thickness of the lines represents the number of interacting receptor-ligand pairs between cells. (B) Bubble plot showing the probability of ligand-receptor pairs between NK cells and keratinocytes. The size of the bubbles indicates the p-value, with smaller p-values resulting in larger bubbles. The color indicates the probability of communication. (C) Flow cytometry detection of live and dead HaCaT cells in Figures 7T and 7U.

