## supplemental tables for "Single cell transcriptome profiling reveals pathogenesis of Bullous Pemphigoid"

| Table S1 \| Bullous pemphigoid samples for scRNA-Seq, bulk RNA-seq and flow cytometry | | | | | | | |
|---|---|---|---|---|---|---|---|
| Sample ID# | Experiment | Dagnosis | Sample type | Sex | Age/year | BPDAI | Blister score |
| BP 1 | scRNA-seq and flow cytometry | Bullous pemphigoid | Skin and blood | Female | 83 | 65.3 | 36 |
| BP 2 | scRNA-seq and flow cytometry | Bullous pemphigoid | Skin and blood | Female | 37 | 42 | 5 |
| BP 3 | scRNA-seq and flow cytometry | Bullous pemphigoid | Skin | Female | 27 | 42.4 | 16.9 |
| BP 4 | scRNA-seq and flow cytometry | Bullous pemphigoid | Skin | Male | 58 | 75 | 39 |
| BP 5 | bulk RNA-seq and flow cytometry | Bullous pemphigoid | Blood | Male | 63 | 83 | 53 |
| BP 6 | bulk RNA-seq and flow cytometry | Bullous pemphigoid | Blood | Male | 64 | 77 | 67 |
| BP 7 | bulk RNA-seq and flow cytometry | Bullous pemphigoid | Blood | Female | 71 | 46 | 16 |
| BP 8 | bulk RNA-seq and flow cytometry | Bullous pemphigoid | Blood | Male | 48 | 59 | 31 |
| BP 9 | bulk RNA-seq and flow cytometry | Bullous pemphigoid | Blood | Male | 68 | 66 | 38 |
| BP 10 | flow cytometry | Bullous pemphigoid | Blood | Female | 63 | 109 | 68 |
| BP 11 | flow cytometry | Bullous pemphigoid | Blood | Female | 72 | 40 | 5 |
| BP 12 | flow cytometry | Bullous pemphigoid | Blood | Male | 87 | 63 | 32 |
| BP 13 | flow cytometry | Bullous pemphigoid | Blood | Male | 50 | 166 | 91 |
| BP 14 | flow cytometry | Bullous pemphigoid | Blood | Male | 24 | 83 | 33 |
| BP 15 | flow cytometry | Bullous pemphigoid | Blood | Male | 51 | 31 | 1 |
| BP 16 | flow cytometry | Bullous pemphigoid | Blood | Female | 69 | 42.9 | 18 |
| BP 17 | flow cytometry | Bullous pemphigoid | Blood | Female | 47 | 69 | 36 |
| BP 18 | flow cytometry | Bullous pemphigoid | Blood | Male | 24 | 28.3 | 11.3 |
| BP 19 | flow cytometry | Bullous pemphigoid | Blood | Male | 62 | 40.9 | 14.6 |
| BP 20 | flow cytometry | Bullous pemphigoid | Blood | Female | 73 | 49 | 19 |
| BP 21 | flow cytometry | Bullous pemphigoid | Blood | Male | 62 | 39 | 7 |
| BP 22 | flow cytometry | Bullous pemphigoid | Blood | Male | 24 | 26 | 12 |
| BP 23 | flow cytometry | Bullous pemphigoid | Blood | Female | 81 | 34.3 | 31.3 |
| BP 24 | flow cytometry | Bullous pemphigoid | Blood | Female | 80 | 96 | 51 |
| BP 25 | flow cytometry | Bullous pemphigoid | Blood | Female | 76 | 18.8 | 16.8 |
| BP 26 | flow cytometry | Bullous pemphigoid | Blood | Female | 58 | 36.3 | 8.3 |
| BP 27 | flow cytometry | Bullous pemphigoid | Blood | Female | 80 | 48.2 | 34.6 |
| BP 28 | flow cytometry | Bullous pemphigoid | Blood | Female | 69 | 42.9 | 18 |
| BP 29 | flow cytometry | Bullous pemphigoid | Blood | Female | 80 | 96 | 51 |
| BP 30 | flow cytometry | Bullous pemphigoid | Blood | Male | 82 | 43 | 26 |
| BP 31 | flow cytometry | Bullous pemphigoid | Blood | Female | 58 | 36.3 | 18.3 |
| BP 32 | flow cytometry | Bullous pemphigoid | Blood | Male | 68 | 12.6 | 11 |
| BP 33 | flow cytometry | Bullous pemphigoid | Blood | Male | 70 | 47 | 47 |
| BP 34 | flow cytometry | Bullous pemphigoid | Blood | Female | 27 | 42.4 | 16.9 |
| BP 35 | flow cytometry | Bullous pemphigoid | Blood | Female | 61 | 17.2 | 17.2 |
| BP 36 | flow cytometry | Bullous pemphigoid | Blood | Male | 68 | 20.6 | 13.3 |
| BP 37 | flow cytometry | Bullous pemphigoid | Blood | Male | 85 | 5.6 | 3 |
| BP 38 | flow cytometry | Bullous pemphigoid | Blood | Male | 58 | 75 | 39 |
| BP 39 | flow cytometry | Bullous pemphigoid | Blood | Female | 80 | 81 | 30 |
| BP 40 | flow cytometry | Bullous pemphigoid | Blood | Male | 41 | 3 | 3 |
| BP 41 | flow cytometry | Bullous pemphigoid | Blood | Male | 58 | 34.6 | 27.6 |
| BP 42 | flow cytometry | Bullous pemphigoid | Blood | Male | 78 | 80.2 | 21.6 |
| BP 43 | flow cytometry | Bullous pemphigoid | Blood | Male | 68 | 86 | 82 |
| BP 44 | flow cytometry | Bullous pemphigoid | Blood | Male | 58 | 38.6 | 29.6 |
| BP 45 | flow cytometry | Bullous pemphigoid | Blood | Male | 89 | 14 | 13 |
| BP 46 | flow cytometry | Bullous pemphigoid | Blood | Male | 70 | 29.6 | 13.6 |

| | | | | | | | |
|---|---|---|---|---|---|---|---|
| BP 47 | flow cytometry | Bullous pemphigoid | Blood | Male | 59 | 19.2 | 19.2 |
| BP 48 | flow cytometry | Bullous pemphigoid | Blood | Female | 63 | 26 | 9 |
| BP 49 | flow cytometry | Bullous pemphigoid | Blood | Male | 59 | 6 | 1 |
| BP 50 | flow cytometry | Bullous pemphigoid | Blood | Female | 72 | 0 | 0 |
| BP 51 | flow cytometry | Bullous pemphigoid | Blood | Female | 63 | 7 | 6 |
| BP 52 | flow cytometry | Bullous pemphigoid | Blood | Female | 72 | 23 | 3 |
| BP 53 | flow cytometry | Bullous pemphigoid | Blood | Female | 72 | 12 | 0 |
| BP 54 | flow cytometry | Bullous pemphigoid | Blood | Male | 59 | 14.6 | 1 |
| BP 55 | flow cytometry | Bullous pemphigoid | Blood | Female | 69 | 6 | 0 |
| BP 56 | flow cytometry | Bullous pemphigoid | Blood | Male | 59 | 0 | 0 |
| BP 57 | flow cytometry | Bullous pemphigoid | Blood | Female | 39 | 0 | 0 |
| BP 58 | flow cytometry | Bullous pemphigoid | Blood | Female | 63 | 0 | 0 |
| BP 59 | flow cytometry | Bullous pemphigoid | Blood | Male | 59 | 0 | 0 |
| BP 60 | flow cytometry | Bullous pemphigoid | Blood | Female | 72 | 28.2 | 4 |
| BP 61 | flow cytometry | Bullous pemphigoid | Blood | Male | 51 | 0 | 0 |
| BP 62 | flow cytometry | Bullous pemphigoid | Blood | Male | 63 | 0 | 0 |
| BP 63 | flow cytometry | Bullous pemphigoid | Blood | Male | 49 | 1.3 | 0 |
| BP 64 | flow cytometry | Bullous pemphigoid | Blood | Male | 59 | 0 | 0 |
| BP 65 | flow cytometry | Bullous pemphigoid | Blood | Male | 78 | 0 | 0 |
| BP 66 | flow cytometry | Bullous pemphigoid | Blood | Male | 69 | 0 | 0 |
| BP 67 | flow cytometry | Bullous pemphigoid | Blood | Female | 73 | 0 | 0 |
| BP 68 | flow cytometry | Bullous pemphigoid | Blood | Male | 62 | 0 | 0 |
| BP 69 | flow cytometry | Bullous pemphigoid | Blood | Male | 24 | 0 | 0 |
| BP 70 | flow cytometry | Bullous pemphigoid | Blood | Female | 63 | 26 | 9 |
| BP 71 | flow cytometry | Bullous pemphigoid | Blood | Female | 59 | 6 | 1 |
| BP 72 | flow cytometry | Bullous pemphigoid | Blood | Male | 59 | 0 | 0 |
| BP 73 | flow cytometry | Bullous pemphigoid | Blood | Male | 48 | 0 | 0 |
| BP 74 | flow cytometry | Bullous pemphigoid | Blood | Female | 63 | 0 | 0 |
| BP 75 | flow cytometry | Bullous pemphigoid | Blood | Female | 83 | 0 | 0 |
| BP 76 | flow cytometry | Bullous pemphigoid | Blood | Male | 51 | 0 | 0 |
| BP 77 | flow cytometry | Bullous pemphigoid | Blood | Female | 73 | 0 | 0 |
| BP 78 | flow cytometry | Bullous pemphigoid | Blood | Female | 21 | 26.6 | 26.6 |
| BP 79 | flow cytometry | Bullous pemphigoid | Blood | Female | 58 | 5 | 5 |
| BP 80 | flow cytometry | Bullous pemphigoid | Blood | Male | 24 | 0 | 0 |
| BP 81 | flow cytometry | Bullous pemphigoid | Blood | Male | 78 | 3 | 0 |
| BP 82 | flow cytometry | Bullous pemphigoid | Blood | Female | 83 | 0 | 0 |
| BP 83 | flow cytometry | Bullous pemphigoid | Blood | Female | 74 | 0 | 0 |
| BP 84 | flow cytometry | Bullous pemphigoid | Blood | Female | 66 | 0 | 0 |
| BP 85 | flow cytometry | Bullous pemphigoid | Blood | Female | 38 | 3 | 0 |
| BP 86 | flow cytometry | Bullous pemphigoid | Blood | Male | 24 | 0 | 0 |
| BP 87 | flow cytometry | Bullous pemphigoid | Blood | Male | 60 | 0 | 0 |
| BP 88 | flow cytometry | Bullous pemphigoid | Blood | Male | 68 | 0 | 0 |
| BP 89 | flow cytometry | Bullous pemphigoid | Blood | Female | 28 | 0 | 0 |
| HC 1 | scRNA-seq | Healthy control | Blood | Female | 71 | NA | NA |
| HC 2 | scRNA-seq | Healthy control | Blood | Female | 40 | NA | NA |
| HC 3 | bulk RNA-seq | Healthy control | Blood | Male | 65 | NA | NA |
| HC 4 | bulk RNA-seq | Healthy control | Blood | Male | 67 | NA | NA |
| HC 5 | bulk RNA-seq | Healthy control | Blood | Female | 67 | NA | NA |
| HC 6 | bulk RNA-seq | Healthy control | Blood | Male | 49 | NA | NA |
| HC 7 | bulk RNA-seq | Healthy control | Blood | Male | 65 | NA | NA |
| HC 8 | flow cytometry | Healthy control | Blood | Male | 87 | NA | NA |
| HC 9 | flow cytometry | Healthy control | Blood | Female | 83 | NA | NA |
| HC 10 | flow cytometry | Healthy control | Blood | Female | 82 | NA | NA |
| HC 11 | flow cytometry | Healthy control | Blood | Male | 82 | NA | NA |
| HC 12 | flow cytometry | Healthy control | Blood | Female | 78 | NA | NA |
| HC 13 | flow cytometry | Healthy control | Blood | Male | 75 | NA | NA |
| HC 14 | flow cytometry | Healthy control | Blood | Male | 70 | NA | NA |
| HC 15 | flow cytometry | Healthy control | Blood | Female | 67 | NA | NA |
| HC 16 | flow cytometry | Healthy control | Blood | Male | 65 | NA | NA |
| HC 17 | flow cytometry | Healthy control | Blood | Male | 67 | NA | NA |
| HC 18 | flow cytometry | Healthy control | Blood | Male | 65 | NA | NA |
| HC 19 | flow cytometry | Healthy control | Blood | Female | 61 | NA | NA |
| HC 20 | flow cytometry | Healthy control | Blood | Female | 60 | NA | NA |
| HC 21 | flow cytometry | Healthy control | Blood | Female | 48 | NA | NA |

| | | | | | | | |
|---|---|---|---|---|---|---|---|
| HC 22 | flow cytometry | Healthy control | Blood | Female | 39 | NA | NA |
| HC 23 | flow cytometry | Healthy control | Blood | Female | 27 | NA | NA |
| HC 24 | flow cytometry | Healthy control | Blood | Female | 78 | NA | NA |
| HC 25 | flow cytometry | Healthy control | Blood | Female | 73 | NA | NA |
| HC 26 | flow cytometry | Healthy control | Blood | Female | 72 | NA | NA |
| HC 27 | flow cytometry | Healthy control | Blood | Male | 67 | NA | NA |
| HC 28 | flow cytometry | Healthy control | Blood | Male | 64 | NA | NA |
| HC 29 | flow cytometry | Healthy control | Blood | Female | 64 | NA | NA |
| HC 30 | flow cytometry | Healthy control | Blood | Male | 63 | NA | NA |
| HC 31 | flow cytometry | Healthy control | Blood | Female | 63 | NA | NA |
| HC 32 | flow cytometry | Healthy control | Blood | Female | 62 | NA | NA |
| HC 33 | flow cytometry | Healthy control | Blood | Female | 60 | NA | NA |
| HC 34 | flow cytometry | Healthy control | Blood | Male | 56 | NA | NA |
| HC 35 | flow cytometry | Healthy control | Blood | Male | 52 | NA | NA |
| HC 36 | flow cytometry | Healthy control | Blood | Female | 49 | NA | NA |
| HC 37 | flow cytometry | Healthy control | Blood | Male | 49 | NA | NA |
| HC 38 | flow cytometry | Healthy control | Blood | Male | 45 | NA | NA |
| HC 39 | flow cytometry | Healthy control | Blood | Male | 42 | NA | NA |
| HC 40 | flow cytometry | Healthy control | Blood | Male | 35 | NA | NA |
| HC 41 | flow cytometry | Healthy control | Blood | Female | 30 | NA | NA |
| HC 42 | flow cytometry | Healthy control | Blood | Female | 28 | NA | NA |
| HC 43 | flow cytometry | Healthy control | Blood | Male | 26 | NA | NA |
| HC 44 | flow cytometry | Healthy control | Blood | Female | 23 | NA | NA |

Note: BP bullous pemphigoid; HC, healthy control; BPDAI, bullous pemphigoid disease area index; NA, not applicable.

| Table S2 \| Quality control for scRNA-Seq | | | |
|---|---|---|---|
| **Sample** | **Estimated Number of Cells** | **Fraction Reads in Cells** | **Mean Reads per Cell** |
| BP1-L | 18,172 | 91.10% | 21,184 |
| BP2-L | 11,456 | 83.00% | 31,936 |
| BP3-L | 10,187 | 94.40% | 35,522 |
| BP4-L | 12,361 | 93.40% | 29,287 |
| BP1-NL | 15,949 | 82.20% | 27,282 |
| BP2-NL | 16,176 | 84.60% | 26,154 |
| BP3-NL | 14,680 | 93.00% | 24,779 |
| BP4-NL | 14,063 | 95.20% | 25,711 |
| BP1 active | 11,026 | 96.50% | 38,573 |
| BP2 active | 1,870 | 86.70% | 107,747 |
| BP1 remission | 8,509 | 95.10% | 51,026 |
| BP2 remission | 2,078 | 94.00% | 95,209 |
| HC1 PBMC | 7,480 | 91.30% | 60,476 |
| HC2 PBMC | 7,891 | 89.70% | 56,450 |

| Sample | Median Genes per Cell | Total Genes Detected | Median UMI Counts per Cell |
|---|---|---|---|
| BP1-L | 1,852 | 27,334 | 4,739 |
| BP2-L | 1,404 | 27,541 | 3,097 |
| BP3-L | 1,196 | 26,746 | 3,076 |
| BP4-L | 1,507 | 24,510 | 3,836 |
| BP1-NL | 1,644 | 28,005 | 3,869 |
| BP2-NL | 1,385 | 28,370 | 2,881 |
| BP3-NL | 1,304 | 27,499 | 3,502 |
| BP4-NL | 1,431 | 25,890 | 3,744 |
| BP1 active | 1,844 | 23,231 | 5,249 |
| BP2 active | 994 | 20,151 | 4,363 |
| BP1 remission | 2,064 | 23,168 | 5,959 |
| BP2 remission | 1,984 | 20,506 | 5,837 |
| HC1 PBMC | 1,211 | 20,801 | 2,466 |
| HC2 PBMC | 1,161 | 21,112 | 2,590 |

| Sample | Reads Mapped Confidently to Genome | Reads Mapped Confidently to Intergenic Regions | Reads Mapped Confidently to Intronic Regions |
|---|---|---|---|
| BP1-L | 84.30% | 3.50% | 4.40% |
| BP2-L | 69.20% | 4.20% | 5.20% |
| BP3-L | 80.00% | 3.40% | 3.80% |
| BP4-L | 75.90% | 2.70% | 3.60% |
| BP1-NL | 79.00% | 5.00% | 6.10% |
| BP2-NL | 83.90% | 4.60% | 5.60% |
| BP3-NL | 84.70% | 3.10% | 4.10% |
| BP4-NL | 73.80% | 3.20% | 4.20% |
| BP1 active | 81.90% | 2.30% | 5.70% |
| BP2 active | 57.60% | 3.40% | 7.10% |
| BP1 remission | 82.40% | 2.60% | 6.20% |
| BP2 remission | 79.20% | 2.90% | 7.90% |
| HC1 PBMC | 65.80% | 4.20% | 7.00% |
| HC2 PBMC | 66.70% | 4.20% | 6.10% |

| Sample | Reads Mapped Confidently to Exonic Regions | Reads Mapped Confidently to Transcriptome |
|---|---|---|
| BP1-L | 76.30% | 67.90% |
| BP2-L | 59.80% | 54.40% |
| BP3-L | 72.80% | 63.00% |
| BP4-L | 69.60% | 63.60% |
| BP1-NL | 67.90% | 60.50% |
| BP2-NL | 73.60% | 67.20% |
| BP3-NL | 77.50% | 67.80% |
| BP4-NL | 66.40% | 61.20% |
| BP1 active | 73.90% | 67.50% |
| BP2 active | 47.10% | 43.20% |
| BP1 remission | 73.70% | 67.00% |
| BP2 remission | 68.50% | 61.10% |
| HC1 PBMC | 54.60% | 48.50% |
| HC2 PBMC | 56.30% | 50.50% |

| Table S3 \| Cluster unique and shared DEgenes of KCs, FBs, PCs and ECs | | | | |
|---|---|---|---|---|
| **Cluster unique upregulated DEgenes of keratinocytes in BP lesional skin** | | | | |
| Undiff-KC | Prolif-KC | Diff-KC1 | Diff-KC2 | HFAC |
| OS9 | TRIM25 | SERPINB6 | NPM1 | IGFBP7 |
| LACTB | GTF2H5 | ENSG0000012 4593 | SLC25A3 | MRPS6 |
| IMP3 | ORAI1 | ANKRD22 | PABPC1 | MRPL14 |
| KHSRP | CLPTM1L | UFD1 | COX4I1 | POLR2F |
| ABLIM1 | BRI3BP | PDHA1 | PRDX1 | TMSB4X |
| SELENOS | SLX9 | HSF1 | CDKN1A | TNC |
| TIMM23 | TMEM256 | ATP2A2 | HINT1 | MRPL3 |
| NAP1L4 | OTUB1 | TSR1 | | SRP72 |
| TMEM238 | BRMS1 | STX6 | | NSFL1C |
| PSME1 | GRINA | CKAP4 | | MRPL24 |
| GPC1 | BMP4 | DESI1 | | ARF5 |
| DNAJC7 | CDKN2D | EHD4 | | LDHB |
| KPNB1 | SCAND1 | NUDT8 | | PTBP1 |
| SPPL3 | HRAS | SNX5 | | DRAP1 |
| UBAP1 | PPP2R5E | CMIP | | MKNK2 |
| COL27A1 | CTDSP2 | IRF2BP2 | | CRIP2 |
| ODC1 | MOB1A | SIAH2 | | BUD23 |
| CXADR | PPP1R35 | PSMC1 | | NAA38 |
| MTCH1 | JPT2 | GALK1 | | PPP2R1A |
| ACBD3 | CDC34 | ARID5B | | MRPL54 |
| SERTAD2 | EPRS1 | COPZ1 | | TRMT112 |
| SMIM29 | TBCA | ITGB1BP1 | | LSM4 |
| ZNF385A | PHF19 | PTK6 | | ATP5MC1 |
| STARD5 | RASSF7 | RAB35 | | NAA20 |
| CRYBG1 | POSTN | LMAN1 | | COA3 |
| DSP | NCOA4 | FAM83C | | NNMT |
| NECTIN2 | CDC25B | CLTA | | ECHS1 |
| OPTN | FXYD3 | MLX | | APRT |
| GSK3B | GLRX5 | EIF2B2 | | DNAJC15 |
| LUZP1 | RRM2 | EIF4G3 | | IFITM2 |
| PRRC2B | TLE1 | STRN3 | | SNHG16 |
| CENPX | BCL2L12 | PRNP | | UBALD2 |
| NDUFAF4 | FTSJ1 | CHIC2 | | CTNNB1 |
| SPEN | DERL1 | FAM83A | | CTNNBIP1 |
| SPINT1 | COPRS | BZW1 | | BANF1 |
| HSPA6 | CCNC | PTRH1 | | YWHAZ |
| OXSR1 | ALDH16A1 | CSNK1E | | FKBP4 |
| KDM2A | TP53RK | PACSIN3 | | SERF2 |
| BRD7 | BIRC5 | TAX1BP3 | | PUF60 |
| AK4 | MRPL37 | SQOR | | MBD2 |
| ANKLE2 | NUDT5 | TMEM45B | | DNPH1 |
| ETV3 | SIGMAR1 | RSU1 | | H2AZ2 |
| CRB3 | MIB2 | NAPG | | CCT2 |
| PHF23 | EMC10 | IDH3A | | DNAJA2 |
| SZRD1 | NOTCH1 | TUT7 | | ATG3 |
| TPPP3 | MDFI | FBLIM1 | | SPARC |
| DENND2B | RPS19BP1 | IFRD2 | | EDF1 |
| AAMP | CCND3 | GOT1 | | G3BP1 |
| HEBP2 | PAM16 | IGSF8 | | SNRPD1 |
| CSNK1D | ALG3 | ASPH | | IMPDH2 |
| TNFRSF19 | CIAPIN1 | LEO1 | | CDK4 |
| RNASEK | RNPEP | SLC10A6 | | PSMD6 |

| | | | | |
|---|---|---|---|---|
| MPG | CDC123 | ENSG00000282034 | | HSPA4 |
| DDR1 | INF2 | NAGK | | SNRPD3 |
| USP47 | POLR3K | TGFA | | LGALS1 |
| SGMS1 | RNASEH2C | HMOX2 | | CALML5 |
| CDK12 | GRK6 | MRPS25 | | GTF3A |
| PLEC | SMC1A | LLGL2 | | SNRPB |
| ANKRD11 | KCTD1 | APOBEC3A | | HMGA1 |
| GATA3 | TMA7 | FAM110C | | TOMM5 |
| GNB4 | MCTS1 | MVD | | PHPT1 |
| SERTAD3 | RPS10-NUDT3 | EPPK1 | | COMT |
| PICALM | GET3 | SRP19 | | PRDX4 |
| EIF4E2 | PPP2R5A | IPO7 | | MRPL23 |
| CCDC137 | LMNB1 | MAPKAPK3 | | MAP2K2 |
| PTPRF | TPD52 | DAPP1 | | GNAI3 |
| EVPL | KMT5A | KCNK6 | | GCSH |
| GJB5 | TBCD | DBNDD2 | | PTMS |
| TRIP10 | MRPS33 | NANS | | RPS2 |
| YY1 | RCC1 | DHRS1 | | METTL26 |
| LSM1 | NARF | ATP6V1D | | MRPS35 |
| STAT3 | MRPL33 | PSMD14 | | CYBA |
| IMP4 | ZNF750 | SDCBP2 | | STAU1 |
| TMEM259 | PRPF19 | ALDH3B2 | | HSPA5 |
| TAP1 | CAVIN1 | ATP6V1B2 | | PSMC5 |
| TFAP2A | CDC20 | FBXL6 | | IGFBP4 |
| DDT | EI24 | MED15 | | ATF4 |
| SMAP1 | STARD7 | CLK3 | | PSMA4 |
| ADAMTS1 | TACC3 | METTL9 | | ATP6V1F |
| SRPRA | VSIR | ACP1 | | ELOB |
| EWSR1 | MRPL11 | DNAJC1 | | |
| LAMC2 | MT1M | STRN | | |
| F11R | CISH | MYL12A | | |
| SRSF4 | MRPL58 | PELI1 | | |
| IQGAP1 | NT5C | SORD | | |
| UBE2Z | CDKN3 | KCMF1 | | |
| NXF1 | SAC3D1 | BNIP3 | | |
| FAT2 | ALDH9A1 | DCTPP1 | | |
| ATG101 | UBE2C | MIR222HG | | |
| ATP2B4 | GIT1 | DUSP22 | | |
| DNAJB6 | NDUFA11 | ZPR1 | | |
| FBXO11 | SNX17 | PCBD1 | | |
| KIAA1522 | NUBP2 | PPME1 | | |
| TAF15 | SLC25A11 | SOWAHC | | |
| MIER1 | ZBTB7B | BLVRB | | |
| ITPRIP | CCDC25 | LINC01605 | | |
| KDM5B | DPM3 | IL6ST | | |
| SPPL2A | ECH1 | SDHB | | |
| SRRT | DTYMK | AGO2 | | |
| FGD6 | ATG4B | TUBGCP2 | | |
| PRKDC | DNAJC19 | CASZ1 | | |
| PPP1R14C | MYBL2 | MBD1 | | |
| SF3B2 | WIPI2 | YARS1 | | |
| USP22 | NDUFS5 | OSMR | | |
| MAPK6 | CLPTM1 | USP53 | | |
| RNF213 | PLA2G4F | PPP1R11 | | |
| PLK3 | RWDD1 | ARHGEF1 | | |
| TM9SF3 | SKA2 | SGPP2 | | |

| | | |
|---|---|---|
| PPP1R18 | HNRNPUL1 | TACSTD2 |
| NPTN | PARP1 | TNIP1 |
| YY1AP1 | LRP10 | SNHG7 |
| STAP2 | PPP5C | STIM1 |
| PSMB9 | PRPF31 | EMC4 |
| UBE2A | NSMCE1 | NPDC1 |
| PAFAH1B1 | MRPL15 | ATP1A1 |
| NRG1 | GGH | TMEM41A |
| SETD5 | DDX49 | TWF2 |
| NAA50 | HMGB3 | SLMAP |
| PSMF1 | DNM2 | DAPK3 |
| RBCK1 | MRPL34 | SUCO |
| IDH3G | SURF2 | PLA2G3 |
| ZNF267 | CCNB1 | VPS37B |
| ADAM15 | SREBF1 | PRKCD |
| HCAR3 | FIS1 | LITAF |
| ATXN2L | OGDH | DHRS7 |
| MTLN | SMS | EID1 |
| ATN1 | FUOM | MYO1E |
| DYNC1H1 | H19 | SDHC |
| PRXL2A | SMARCA4 | LINC00958 |
| ANXA4 | TARS1 | PTPN1 |
| ARF6 | TBL3 | POLE4 |
| DDX39A | PAK4 | UBE2V1 |
| SH3GLB1 | PGP | MAP4K4 |
| RIOK3 | FAAP20 | NBEAL2 |
| PCBP1 | B4GALT2 | TPD52L2 |
| NCBP2AS2 | PHB2 | PRELID3B |
| ACTN4 | MAP3K14 | EPHX3 |
| RPL37A | DGKZ | CASP4 |
| RABGGTB | CCNA2 | TRAPPC3 |
| CMPK1 | EMC8 | FBXO9 |
| PLEKHM2 | SLC16A14 | DPH1 |
| BAG3 | ARID1A | ARL8A |
| TNKS1BP1 | VARS1 | FARSB |
| TRIM28 | TSSC4 | KDELR1 |
| EIF2S3 | MRPS18C | PLIN2 |
| ANXA8L1 | HDHD5 | CDKN2B |
| COX17 | MGAT1 | PGM2 |
| PRRC2C | ANKRD9 | RALGPS2 |
| EMP3 | MRPS5 | SLC25A25 |
| | UQCC3 | TMPRSS4 |
| | TROAP | NIPAL4 |
| | NDUFAF3 | MFSD10 |
| | NDUFA2 | PSMG1 |
| | TRABD | CYP51A1 |
| | YIF1B | KATNBL1 |
| | VMA21 | UBE2D1 |
| | NMRAL1 | SIRT7 |
| | TTLL12 | METRNL |
| | NPM3 | ERP44 |
| | NDUFA3 | STK24 |
| | H1-0 | HSPB8 |
| | UBXN6 | RAB22A |
| | GAS2L1 | SLC4A7 |
| | MRPS23 | CYB5R2 |
| | MPDU1 | PLSCR3 |
| | JAGN1 | PDZD11 |

| | | |
|---|---|---|
| | DDX56 | MT-CO1 |
| | ADAR | GNAS |
| | CENPB | PLA2G4E |
| | COA4 | RRP36 |
| | CCM2 | C16orf91 |
| | PNKP | PSMD1 |
| | RUSC1 | |
| | MKKS | |
| | RABL6 | |
| | STX10 | |
| | TRAPPC4 | |
| | ENSG00000255639 | |
| | HDAC3 | |
| | FEN1 | |
| | HDDC2 | |
| | LGALS3BP | |
| | BCL9L | |
| | PTOV1 | |
| | KPNA2 | |
| | MRPS28 | |
| | FAM83D | |
| | DNTTIP1 | |
| | TLE3 | |
| | UPF1 | |
| | AGAP3 | |
| | RUVBL1 | |
| | FIBP | |
| | MBD3 | |
| | ACOT7 | |
| | ACAA1 | |
| | HCFC1 | |
| | PTGES | |
| | ADAMTS4 | |
| | DDX41 | |
| | TSPAN4 | |
| | BCL11B | |
| | SEMA3F | |
| | HDAC1 | |
| | COX16 | |
| | PIN1 | |
| | CEP55 | |
| | KEAP1 | |
| | KCNK7 | |
| | SSRP1 | |
| | BLCAP | |
| | EFCAB14 | |
| | TOB2 | |
| | CHCHD3 | |
| | ZNF511 | |
| | MRPL55 | |
| | FAM102A | |
| | URM1 | |
| | ENDOG | |
| | TIMMDC1 | |
| | PIEZO1 | |
| | AGTRAP | |

| Cluster unique downregulated DEgenes of keratinocytes in BP lesional skin | | | | |
|---|---|---|---|---|
| Undiff-KC | Prolif-KC | Diff-KC1 | Diff-KC2 | HFAC |
| JAG2 | ARL6IP6 | EIF3L | MT-ND1 | SCPEP1 |
| PITPNB | NQO2 | RPL4 | HNRNPDL | PURA |
| KRT31 | SMARCA2 | LSS | HNRNPH1 | NR4A2 |
| CD81 | TRMT10C | ZC2HC1A | IFFO2 | TMEM87A |
| CXCL2 | FAF1 | NXT1 | SF1 | CD9 |
| NDUFS4 | SLC7A1 | RPS14 | HSPE1 | S100A13 |
| ALKBH7 | VCL | MCL1 | DDX21 | LMBRD1 |
| TNFSF9 | PLEKHA1 | SS18L2 | NDEL1 | TFCP2L1 |
| LAMA3 | UPF3B | HINT2 | TCP1 | SOX9 |
| PRDX5 | SPATS2L | RPS7 | DDX3X | KLF9 |
| PITHD1 | RBPJ | DBP | MT-ND2 | IER2 |
| RNF227 | BCL10 | PRKD2 | HNRNPU | PCM1 |
| LAMTOR3 | BTAF1 | NMU | KDM6B | AOPEP |
| UXT | HIVEP2 | PIM1 | NECTIN4 | CITED4 |
| HEXIM1 | ESF1 | LARS1 | RTN4 | STOM |
| PGRMC1 | ITGB1 | ESPN | TRA2B | GSN |
| MORF4L2 | SIAH1 | SIN3A | FGFR3 | ITGB8 |
| CKB | SUN1 | ULK3 | MT-ATP8 | MAGED2 |
| ADH5 | OXR1 | UBC | TAF1D | TAX1BP1 |
| EIF4A3 | PIP5K1A | GAS6 | GOLGA4 | CSRP1 |
| TOMM20 | BNIP3L | POLR1F | | FOXC1 |
| NENF | USP34 | CAPNS2 | | OXA1L |
| IER5L | ENSG00000283674 | RPL36 | | CYSTM1 |
| H2AX | SCCPDH | KLF13 | | DPP7 |
| H2AZ1 | NCK1 | NUCKS1 | | PIP |
| FOXQ1 | ENSG00000287906 | MAP3K2 | | RBP1 |
| H3-3B | EML4 | PIM3 | | IGFL2-AS1 |
| MGST3 | C1D | CAMSAP3 | | NDUFA5 |
| CDCA7L | PPP1CC | MPST | | SPARCL1 |
| RASD1 | NRM | EEF1A1 | | ETV6 |
| ATP6V0C | TATDN1 | LAMTOR4 | | IFI16 |
| NASP | MYADM | RPL37 | | SCGB1D2 |
| MAT2A | ZCCHC8 | ELF1 | | SCGB2A1 |
| NEDD9 | CENPK | KDM5A | | PLPP1 |
| ANKRD37 | IP6K2 | ENSG00000261189 | | XBP1 |
| CYCS | RAMAC | SELENOP | | TIMP3 |
| ABHD5 | CDK6 | RPS13 | | C15orf61 |
| ARL4D | PAFAH1B3 | SMARCC2 | | TEX264 |
| MAPRE1 | C11orf1 | RPL22L1 | | MTUS1 |
| SRSF7 | PPP1R12A | RPS25 | | PCMTD1 |
| COPZ2 | GART | PTGS1 | | GJB6 |
| DCXR | CENPU | DUT | | PCNP |
| CKS1B | MARCHF5 | VPS36 | | ATP1B1 |
| ATRAID | SAV1 | MED4 | | ZRANB2 |
| TCEA1 | RAB9A | SCEL | | CCDC47 |
| EIF1 | EMP1 | ENSG00000275993 | | FCGRT |
| LAMB4 | COPS8 | RPS9 | | SEMA3C |
| H1-10 | NOTCH2NLA | NOTCH3 | | MIR205HG |
| POU3F1 | NUTM2B-AS1 | TSEN34 | | SRSF3 |
| EIF5 | DIP2B | GSTA4 | | TTC3 |
| NFKBIA | TCERG1 | WNK1 | | DDAH2 |

| | | | | |
|---|---|---|---|---|
| PEBP1 | BACH1 | RPL19 | | CRNDE |
| CTNNAL1 | RCN2 | NFKBIB | | SAT1 |
| CAMTA1 | GNL1 | RPL36A | | TCF25 |
| JDP2 | AZI2 | PPTC7 | | RUNX1 |
| ELOC | CHORDC1 | PLEKHG5 | | NDRG2 |
| HCAR2 | RIF1 | TOM1L2 | | RBIS |
| POLR2J | C3orf14 | BBC3 | | LTBP4 |
| HTRA1 | PUM3 | NFIX | | FST |
| CDH13 | SNX2 | RTF1 | | SDC4 |
| RGS2 | WDR43 | CBR1 | | NFIA |
| ADIPOR1 | NUP98 | ENSG00000263620 | | ACSL3 |
| MAFB | WDFY2 | MYCBP2 | | LAMP2 |
| ACTG1 | SETD2 | OGA | | NFIC |
| TLE4 | ORC6 | RB1CC1 | | XIST |
| SKP1 | UBR5 | SSB | | NUDT4 |
| SELENOK | MLLT10 | MZT2B | | NET1 |
| | TRIP11 | BICDL1 | | SINHCAF |
| | OGT | ZNF652 | | SOCS3 |
| | HECTD1 | UBE2E3 | | PPP4R2 |
| | CFAP20 | RPL10A | | SCGB1B2P |
| | TBRG1 | COMMD6 | | DTWD1 |
| | PDCL3 | DGCR6L | | ANKRD10 |
| | ABI1 | PLPP2 | | PPFIBP1 |
| | RELB | PIK3C2G | | KRT18 |
| | FNIP1 | HES4 | | CCNI |
| | CDC5L | ENSG00000261215 | | MBNL2 |
| | CEBPZ | DEK | | FOXP1 |
| | ANLN | WTAP | | DANCR |
| | ITPR3 | SYTL1 | | SERINC3 |
| | ACAT1 | SMC3 | | RNPC3 |
| | RECQL | MRPL40 | | RAD21 |
| | PSIP1 | SNW1 | | CHPT1 |
| | ARHGEF5 | UFC1 | | TPD52L1 |
| | ZBTB43 | RPS28 | | EPHX1 |
| | MAST4 | DDB1 | | PYGB |
| | SCML1 | DHX36 | | SLC12A2 |
| | ZNF24 | SUPT16H | | NFIB |
| | RNF11 | VAMP2 | | CDA |
| | ZFC3H1 | TECR | | AVPI1 |
| | SELENOI | RTN3 | | LAPTM4A |
| | SPATS2 | IER3 | | CFD |
| | RAB5A | TPR | | ADIRF |
| | RAB3IP | SMNDC1 | | ZKSCAN1 |
| | PIP4P1 | SCP2 | | ADM |
| | MORC3 | SMIM7 | | LRATD1 |
| | NRBF2 | HS3ST6 | | NDFIP1 |
| | MBP | EPN2 | | SCAF11 |
| | ZNF800 | RARG | | ELF3 |
| | PPRC1 | CTDSPL | | |
| | MCM7 | KRT2 | | |
| | GNL2 | TACC2 | | |
| | MED19 | ATP5IF1 | | |
| | SSBP2 | LY6G6C | | |
| | CHMP2B | DSC1 | | |
| | STX12 | G3BP2 | | |
| | ETNK1 | DPY30 | | |

| | | |
|---|---|---|
| | NFKBID | EEF2K |
| | SMIM14 | OSBP |
| | DIAPH3 | GTF3C1 |
| | NAA15 | |
| | TNKS2 | |
| | FBXL5 | |
| | ABCE1 | |
| | MED28 | |
| | NIPBL | |
| | CCDC6 | |
| | MIS18BP1 | |
| | RBM5 | |
| | HYI | |
| | KRR1 | |
| | RASSF1 | |
| | MGME1 | |
| | RBMX | |
| | SMC2 | |
| | MRPL50 | |
| | SPAG9 | |
| | BGN | |
| | THUMPD3-AS1 | |
| | ZNHIT3 | |
| | ZNF131 | |
| | RACGAP1 | |
| | LIG1 | |
| | HELLS | |
| | CDCA7 | |
| | CCDC186 | |
| | CEP57 | |
| | PAXBP1 | |
| | CYB5R1 | |
| | PRPF38B | |
| | C8orf88 | |
| | FUBP1 | |
| | PHACTR4 | |
| | TSPAN13 | |
| | C1orf52 | |
| | MASTL | |
| | ASPM | |
| | MAP4K5 | |
| | STAG1 | |

| **Cluster shared DEgenes of keratinocytes** | |
|---|---|
| Up in BP lesional skin | Down in BP lesional skin |
| S100A16 | IRF1 |
| TPM4 | JUNB |
| KRT6C | TRA2A |
| CSTB | SQSTM1 |
| FLNB | N4BP2L2 |
| RAN | MYC |
| ATP5F1B | DCD |
| NDRG1 | MIR23AHG |
| KRT6B | JUND |
| MYL12B | SLC38A2 |
| CD59 | ZFP36 |

| | |
|---|---|
| SERPINB4 | ATF3 |
| S100A2 | CXCL14 |
| ARPC5L | PNISR |
| KRT16 | FOSB |
| KRT14 | PER1 |
| KTN1 | NR4A1 |
| COX8A | NFKBIZ |
| HBEGF | SYNE2 |
| CDH3 | MT1X |
| MAP7D1 | DUSP1 |
| S100A9 | INTS6 |
| PPIB | GPBP1 |
| MPZL2 | CEBPD |
| IFITM3 | UBB |
| EHF | FOS |
| SH3BGRL3 | CCNL1 |
| CA2 | NR1D1 |
| PGAM1 | EFNA1 |
| PPIA | RND3 |
| PFN1 | DNAJA1 |
| FABP5 | ANKRD12 |
| KRT5 | RPL34 |
| TAGLN2 | BTG2 |
| SDC1 | JUN |
| SPRR1B | MT-ND3 |
| PGK1 | |
| YWHAQ | |
| CFL1 | |
| CHCHD2 | |
| CSNK1A1 | |
| JUP | |
| TMSB10 | |
| KRT17 | |
| LYPD3 | |
| ALDOA | |
| SBSN | |
| KRT6A | |
| LY6D | |
| PKM | |
| MYH9 | |
| S100A14 | |
| RAC1 | |
| PHLDA2 | |
| EIF5A | |
| P4HB | |
| NDUFA4L2 | |
| RHOA | |
| LGALS7B | |
| ENO1 | |
| TXN | |
| S100A8 | |
| GAPDH | |
| MIF | |
| S100A7 | |
| TPI1 | |
| ATP5MC3 | |
| CST3 | |
| AQP3 | |

| |
|---|
| SFN |
| CD44 |
| S100A10 |
| GADD45A |
| KLF3 |
| DSC2 |
| YWHAB |
| S100A11 |
| GSTP1 |

| Cluster unique upregulated DEgenes of fibroblasts in BP lesional skin | | | | |
|---|---|---|---|---|
| SP-FB | Inf-FB | Str-FB | SR-FB | MES-FB |
| SERPINB6 | PDE4DIP | PTDSS1 | SUMO3 | OS9 |
| MGMT | NUDT22 | MDFIC | DPYD | FARP1 |
| PTRHD1 | EMC3 | HPF1 | PHLDA1 | COX14 |
| REEP5 | LOXL1 | TMEM167B | KCTD10 | KDELR3 |
| MTDH | PHF5A | TNFRSF1B | EEF1B2 | DESI2 |
| SMIM29 | COPB1 | ZDHHC24 | XRCC6 | PCDH18 |
| ISG15 | RPL7 | EXOSC3 | YBX3 | KRT6B |
| TMEM248 | ERGIC3 | AHI1 | ATP6AP2 | TRPS1 |
| CAVIN1 | ANP32B | HSF1 | AUTS2 | GPX8 |
| RGS3 | MICALL2 | KHSRP | TMEM214 | FYN |
| SUCLG1 | SMIM3 | SLC38A5 | MLF2 | ANAPC5 |
| SNRPD2 | A2M | FAM136A | STOM | HACD3 |
| DUSP23 | LINC01140 | DYRK4 | UGDH | RSPO4 |
| ETFB | FBLIM1 | TUSC2 | S1PR2 | CDC26 |
| LOXL2 | RHEB | RRP1 | NOP58 | PRRC2B |
| BST2 | POLR1H | CHSY1 | CSNK1D | NORAD |
| MZT2B | UFM1 | NABP2 | PSMB8 | CRABP1 |
| COX4I1 | CMTM6 | EIF3J | CNP | HEBP2 |
| IL24 | ORMDL1 | NAP1L4 | IGSF10 | ARL2BP |
| MVB12A | RTCB | ACO2 | ZDHHC9 | TUSC1 |
| WIPI1 | NUBP2 | DLGAP4 | POGLUT3 | MRPS18B |
| BEST1 | CXXC5 | CMTM7 | POLR2A | GLG1 |
| RAB2A | HDAC2 | METAP2 | PYGL | ANKRD11 |
| POLD4 | C1orf198 | ELK3 | UBE2K | GNB4 |
| CST3 | CDC42EP4 | AGGF1 | RNPS1 | PRDX2 |
| COX7A1 | DARS1 | UBA5 | SRSF1 | LMF2 |
| MRPL23 | PRPF31 | RPIA | GPCPD1 | VPS29 |
| SMIM7 | HSPB6 | PEX16 | KLC1 | BCL7C |
| TMEM160 | EWSR1 | GTF2F1 | DYNLL1 | COPA |
| CSNK2B | RASSF8 | MYBBP1A | PRXL2C | ECH1 |
| STMP1 | ERI3 | UBE2G1 | UBA2 | VPS28 |
| FADS1 | COX7B | ARHGAP1 | NOP56 | YIPF3 |
| | MGAT1 | PCED1A | NAP1L1 | NDN |
| | CLSTN3 | MAGOHB | P4HA1 | PRCP |
| | PRR16 | YRDC | SF3A3 | ATP6AP1 |
| | RBP5 | FMNL2 | CGREF1 | PTGER3 |
| | PSMD9 | CDC34 | COPS3 | CTNNA1 |
| | SEC22B | BABAM2 | OGDH | PTK7 |
| | UFC1 | TUSC3 | MYCBP2 | MRPL34 |
| | RHOA | MFHAS1 | TNPO1 | GDI1 |
| | CPSF6 | PTPRA | SNED1 | NID1 |
| | CPXM2 | TP53I11 | PTRH2 | TCEA1 |
| | ICAM2 | SLC30A9 | CAMK2D | NID2 |
| | PDLIM7 | EPSTI1 | HSD17B12 | RRAGA |
| | LAGE3 | GOLT1B | PKDCC | NFIC |

| | | | | |
|---|---|---|---|---|
| | DDX39A | PGAP6 | EGR1 | SEC31A |
| | UBD | ADAMTSL4 | PPP1R18 | CCDC107 |
| | PTP4A2 | NCOR2 | GJA1 | ZCRB1 |
| | PCMT1 | BLOC1S4 | IFNAR1 | CBX5 |
| | ENY2 | TMEM223 | APBB1IP | NRP2 |
| | C5orf15 | MRPS18A | HIGD1A | LRPAP1 |
| | URM1 | COMMD8 | HNRNPA1 | RAB1B |
| | NOL7 | RABEPK | SMARCA5 | CUEDC2 |
| | FADS3 | MFSD12 | EIF3A | SFRP1 |
| | | NAGLU | SGK1 | SNX6 |
| | | OGFR | HNRNPU | RNF187 |
| | | TMUB1 | DTWD1 | AKR7A2 |
| | | PLXND1 | ZMYM4 | ANXA11 |
| | | RALB | SLC2A4RG | DYNC1H1 |
| | | PLEKHB2 | PDGFD | TSR2 |
| | | NIPA2 | SLC16A7 | ZNF22 |
| | | HEXA | DDB1 | RTF2 |
| | | MARK3 | INTS10 | GSTK1 |
| | | RBFA | UBAP2L | POLR2J |
| | | TP53RK | FUS | NDUFS2 |
| | | PLPPR2 | PRKAR1A | TSPAN4 |
| | | C1QTNF6 | OSR1 | CPNE1 |
| | | NUDT5 | THBS3 | TGFB1I1 |
| | | NOPCHAP1 | EXT2 | CZIB |
| | | SCO2 | HNRNPF | NDUFB10 |
| | | ABCF2 | ABCC9 | GLI3 |
| | | USP3 | ENTPD1 | AMOTL2 |
| | | QSOX1 | EIF2S3 | |
| | | GAR1 | AP2B1 | |
| | | DIMT1 | PLEKHG1 | |
| | | SLC35F6 | MRPL16 | |
| | | SMIM10L1 | MCFD2 | |
| | | GSK3B | EIF1AX | |
| | | VOPP1 | ENSG00000258017 | |
| | | NSD1 | | |
| | | ENSG00000274213 | | |
| | | GART | | |
| | | SLC39A13 | | |
| | | BYSL | | |
| | | PGM1 | | |
| | | WDR18 | | |
| | | CIAPIN1 | | |
| | | RNPEP | | |
| | | TM4SF1 | | |
| | | RAB7A | | |
| | | POLR3K | | |
| | | DNAJB12 | | |
| | | SIRPA | | |
| | | FKBP14 | | |
| | | JUND | | |
| | | DDX1 | | |
| | | TFRC | | |
| | | TRMT61A | | |
| | | AK4 | | |
| | | IDH3A | | |
| | | RBM28 | | |

| | | |
|---|---|---|
| | | IMPDH1 |
| | | API5 |
| | | GFER |
| | | ADPRS |
| | | LYAR |
| | | THAP4 |
| | | AP1S1 |
| | | MED8 |
| | | PFDN6 |
| | | RING1 |
| | | RNH1 |
| | | DKC1 |
| | | FAM210A |
| | | DPP7 |
| | | TMX3 |
| | | FAHD1 |
| | | NOP14 |
| | | MRPS33 |
| | | ENOPH1 |
| | | CCDC86 |
| | | PRPF19 |
| | | NAGK |
| | | THAP7 |
| | | CTCF |
| | | ISCA1 |
| | | ZMAT2 |
| | | MPHOSPH10 |
| | | HMOX2 |
| | | ATP2B1-AS1 |
| | | CDC37L1 |
| | | CDK12 |
| | | DCTN1 |
| | | PPFIA1 |
| | | ZMPSTE24 |
| | | EXOSC5 |
| | | POP5 |
| | | STARD7 |
| | | THOC6 |
| | | TRAF7 |
| | | TCIRG1 |
| | | LYPLA1 |
| | | DFFA |
| | | MDK |
| | | LRRC41 |
| | | CRAT |
| | | PDZRN3 |
| | | ABRACL |
| | | DHX30 |
| | | CISH |
| | | MAPKAPK3 |
| | | APEH |
| | | MRPL58 |
| | | MYO9B |
| | | DBNDD2 |
| | | RRP9 |
| | | SPAG7 |
| | | GNPNAT1 |
| | | SF1 |

| | | |
|---|---|---|
| | | CA12 |
| | | AIFM2 |
| | | SCARA3 |
| | | WDR46 |
| | | RIPOR1 |
| | | ATP6V1D |
| | | ITGB3 |
| | | SLC38A6 |
| | | ANKRD17 |
| | | SQLE |
| | | RRS1 |
| | | MRPL9 |
| | | SEH1L |
| | | ADAM9 |
| | | TMEM9B |
| | | BIN1 |
| | | SOAT1 |
| | | DCAF7 |
| | | TXNDC9 |
| | | MED15 |
| | | MRTFA |
| | | DRG2 |
| | | RBM42 |
| | | RWDD1 |
| | | PAK1IP1 |
| | | SLC25A24 |
| | | SEMA3C |
| | | MRAS |
| | | GGT5 |
| | | ATF5 |
| | | PPIH |
| | | B4GALT5 |
| | | PPP5C |
| | | C14orf119 |
| | | EIF2AK1 |
| | | POLE3 |
| | | DDX54 |
| | | MICU2 |
| | | TMEM131 |
| | | CMC1 |
| | | ENSG00000280571 |
| | | SERPINB2 |
| | | DDX49 |
| | | RDH11 |
| | | FASN |
| | | CLIC6 |
| | | DDA1 |
| | | SURF2 |
| | | SLCO3A1 |
| | | TEAD4 |
| | | EPB41L2 |
| | | LARP1 |
| | | SREBF1 |
| | | DOCK7 |
| | | SPTAN1 |
| | | SLC25A28 |
| | | QDPR |

| | | |
|---|---|---|
| | | UGP2 |
| | | ATP2B4 |
| | | AFF4 |
| | | MIIP |
| | | AFDN |
| | | NSUN2 |
| | | HDGFL3 |
| | | ZNF703 |
| | | IRAK1 |
| | | B4GALT2 |
| | | AFG3L2 |
| | | SDC4 |
| | | KPNA3 |
| | | CISD2 |
| | | CMSS1 |
| | | MRPS17 |
| | | BRIX1 |
| | | GLMP |
| | | NELFE |
| | | HMGXB3 |
| | | NABP1 |
| | | EMC8 |
| | | GNL2 |
| | | GLRX2 |
| | | NRIP1 |
| | | PPP1R11 |
| | | PROS1 |
| | | CHMP1A |
| | | P3H4 |
| | | PANX1 |
| | | PLK3 |
| | | HDHD5 |
| | | CS |
| | | BCKDK |
| | | STX12 |
| | | AKT1S1 |
| | | ZFAND3 |
| | | TUBG1 |
| | | PGM3 |
| | | PHLDA3 |
| | | DUS3L |
| | | HERC2 |
| | | APP |
| | | GLTP |
| | | PES1 |
| | | NCBP2 |
| | | RIOK1 |
| | | COMMD5 |
| | | CUEDC1 |
| | | RFC2 |
| | | TRABD |
| | | U2AF2 |
| | | AGT |
| | | VMA21 |
| | | EMILIN2 |
| | | SLC25A1 |
| | | NUP58 |
| | | CDK10 |

| | | |
|---|---|---|
| | | GNL3 |
| | | GASK1B |
| | | NPM3 |
| | | NTPCR |
| | | PPFIBP1 |
| | | APOL1 |
| | | SAMM50 |
| | | HGS |
| | | NECAP2 |
| | | NDUFV3 |
| | | NBN |
| | | FH |
| | | ITPRIPL2 |
| | | MLST8 |
| | | UBTF |
| | | EGLN3 |
| | | CCDC6 |
| | | DDX56 |
| | | MYO1E |
| | | ACTL6A |
| | | GTF2H3 |
| | | WDR5 |
| | | HYAL2 |
| | | ADSS2 |
| | | PNKP |
| | | MCC |
| | | PTBP3 |
| | | DSEL |
| | | PTPN1 |
| | | ERAL1 |
| | | ZNF267 |
| | | MRPL19 |
| | | TRMT1 |
| | | TMEM115 |
| | | ECPAS |
| | | DOK5 |
| | | ATXN2L |
| | | MRPS11 |
| | | ENSG00000255639 |
| | | TOR1A |
| | | PCYT1A |
| | | PPP2CB |
| | | NELFCD |
| | | GOLGA7 |
| | | RASA3 |
| | | ST6GALNAC4 |
| | | DMAC1 |
| | | GSS |
| | | CNDP2 |
| | | SUPT6H |
| | | SGTA |
| | | RBM6 |
| | | POLR1E |
| | | ENSG00000280071 |
| | | SLC39A4 |
| | | TMEM33 |

| | | |
|---|---|---|
| | | THRAP3 |
| | | MRPS28 |
| | | DPH1 |
| | | STK25 |
| | | NOCT |
| | | CCNQ |
| | | FARSB |
| | | PNO1 |
| | | SLC50A1 |
| | | GRWD1 |
| | | RUVBL1 |
| | | PIP4K2A |
| | | MBD3 |
| | | ACAA1 |
| | | GSK3A |
| | | GIPC1 |
| | | NSMF |
| | | ADAMTS4 |
| | | TMTC1 |
| | | PMM2 |
| | | RPF2 |
| | | DDX41 |
| | | WDR77 |
| | | NCBP2AS2 |
| | | RPL38 |
| | | RPS6KA4 |
| | | KARS1 |
| | | UTP11 |
| | | ADPGK |
| | | SMPD1 |
| | | SNX8 |
| | | PDIA5 |
| | | RHOT2 |
| | | ADAMTS2 |
| | | NARS1 |
| | | DENND5A |
| | | TYW3 |
| | | TRIB1 |
| | | KEAP1 |
| | | ABI2 |
| | | SLC4A7 |
| | | USB1 |
| | | ZNF326 |
| | | LRRC47 |
| | | MTHFD1L |
| | | UBQLN2 |
| | | MRPS10 |
| | | CAPN2 |
| | | DR1 |
| | | GNPDA1 |
| | | F3 |
| | | NUDCD1 |
| | | BAZ1B |
| | | HIC1 |
| | | METTL1 |
| | | NUDT16 |
| | | TRUB2 |
| | | DNM1L |

| | | | | |
|---|---|---|---|---|
| | | SCNM1 | | |
| | | TRIM27 | | |
| | | GPX3 | | |
| | | PIEZO1 | | |
| | | CIBAR1 | | |
| | | C16orf91 | | |
| | | TSN | | |
| | | IL17RA | | |
| | | EML3 | | |
| | | SRPK2 | | |
| | | EEF1E1 | | |
| **Cluster unique downregulated DEgenes of fibroblasts in BP lesional skin** | | | | |
| SP-FB | Inf-FB | Str-FB | SR-FB | MES-FB |
| DUSP4 | UPF3A | PAF1 | TRIO | IGF1 |
| DNTTIP2 | APOC1 | MKLN1 | ADGRA2 | COCH |
| HMGB1 | GSTM5 | FAM120AOS | OSBPL9 | EIF3H |
| CXCL2 | ABHD14B | CEP350 | NQO2 | MYADM |
| FOXO3 | NR2F2 | KIF5B | SMARCA2 | PLXDC1 |
| NXT1 | SELENBP1 | PMM1 | KMT2A | VAPA |
| ELL2 | FAS | C11orf58 | LRIG3 | RBM3 |
| PRNP | DDX24 | TMEM87A | PCOLCE2 | TNN |
| ZNF331 | NDRG2 | TSG101 | VPS13C | RPS25 |
| AQP1 | C7 | RALBP1 | MICAL1 | ENHO |
| FOSB | CAMK2N1 | SH3KBP1 | ABLIM1 | SNHG6 |
| HNRNPA2B1 | CRLF1 | TNFSF9 | GAS7 | STIP1 |
| MATR3 | KIAA0930 | MIA3 | NIBAN1 | PLPP5 |
| RAB1A | BRI3 | SPART | SDK1 | YWHAE |
| MEG3 | TNXB | PHF20 | APPL2 | DKK2 |
| SLC19A2 | IL11RA | AP1S2 | NDUFA1 | MCUB |
| FOS | NPC2 | RUNX1T1 | RERE | CSDE1 |
| AXL | MAP1LC3A | UTP23 | EHD4 | FNBP1 |
| RASL11A | CYGB | SLC20A1 | AKAP13 | LINC00632 |
| ZFP36L1 | NEGR1 | TATDN1 | ZNF292 | ASPN |
| RTN4 | CH25H | PPP1CB | AFAP1L1 | DNAJC4 |
| CSRP2 | CCNL2 | KRCC1 | S100A4 | |
| NPTX2 | | CUTA | CREB5 | |
| CMPK1 | | HEXIM1 | DKK1 | |
| REL | | PMPCB | CD24 | |
| PNP | | FAM204A | JAZF1 | |
| GNAS | | UQCRC2 | ZMIZ1 | |
| RNF149 | | CCND3 | HRCT1 | |
| | | PLPP3 | HMCN1 | |
| | | NUCKS1 | CAPG | |
| | | PPIL4 | SLC7A8 | |
| | | ERGIC2 | GAA | |
| | | MRPL33 | UBAP1 | |
| | | CALM1 | BCL2L1 | |
| | | BFAR | SUN1 | |
| | | RCN2 | PTGFR | |
| | | AKIRIN2 | PALM | |
| | | CRTAP | ENSG00000255495 | |
| | | RAB21 | ATP2B1 | |
| | | ERVK3-1 | TMEM14A | |
| | | AIMP1 | LYST | |
| | | TMBIM6 | NCK1 | |
| | | ITM2C | DENND4C | |

| | | | |
|---|---|---|---|
| | | PGRMC2 | RHOQ |
| | | MEAF6 | ARHGAP12 |
| | | MED4 | FOXO1 |
| | | RNF115 | FKBP7 |
| | | MPC2 | CAPS |
| | | GPR108 | KANK2 |
| | | SREK1 | BAIAP2 |
| | | SRP9 | TPCN1 |
| | | CEBPZ | CAMK1D |
| | | ABHD5 | CFH |
| | | SEPHS2 | RENBP |
| | | ZNF638 | ITIH5 |
| | | MED13L | MAPT |
| | | PCF11 | NUMA1 |
| | | WDR13 | PKN2 |
| | | NXF1 | SEC22C |
| | | RBM22 | ALDH3A2 |
| | | IAH1 | PIN4 |
| | | IVNS1ABP | NDUFB4 |
| | | IK | CPVL |
| | | ZNF800 | GDF15 |
| | | CDC73 | EPS8 |
| | | PUM1 | VIM-AS1 |
| | | ENSG00000289341 | BMPR2 |
| | | ZNF622 | ZDHHC1 |
| | | TBC1D22A-DT | DDR2 |
| | | SGCE | KMT2C |
| | | C18orf32 | MPST |
| | | MED19 | FGD5-AS1 |
| | | NET1 | TBC1D2B |
| | | WBP11 | LAMTOR4 |
| | | MAP3K20 | FEZ1 |
| | | MRPL40 | RPL37 |
| | | TFAM | CCPG1 |
| | | EID1 | KITLG |
| | | TMX4 | DNM1 |
| | | NDUFB5 | TFDP2 |
| | | KRR1 | UTRN |
| | | CCAR1 | CPB1 |
| | | TANK | C1QTNF2 |
| | | SUPT4H1 | PAQR3 |
| | | HSPB11 | ANTXR2 |
| | | SRP14 | HIGD2A |
| | | ABL1 | HSPB2 |
| | | PPCS | ATP5MG |
| | | HNRNPH2 | AAMDC |
| | | RIOK3 | GPRC5A |
| | | ATP6V1C1 | CTSD |
| | | RSBN1L | LRRC2 |
| | | FUBP1 | GBE1 |
| | | TIAL1 | CYP27A1 |
| | | EFCAB14 | COL16A1 |
| | | SFRP2 | SUN2 |
| | | ATP5IF1 | CCSER2 |
| | | ITGAE | NCOA1 |
| | | PPP2CA | RPAIN |

| | | | |
|---|---|---|---|
| | | G3BP2 | MTSS1 |
| | | DPY30 | RRAGD |
| | | HMGN2 | EBF1 |
| | | ARL5A | MPLKIP |
| | | BTF3 | RHBDF1 |
| | | PNPLA8 | KLHL24 |
| | | UBE2E1 | CD70 |
| | | | SLC38A10 |
| | | | CYP4B1 |
| | | | MIR100HG |
| | | | TPRG1L |
| | | | DPM3 |
| | | | ST3GAL5 |
| | | | STAM |
| | | | HGSNAT |
| | | | SEMA3D |
| | | | ABI1 |
| | | | FNIP1 |
| | | | SGCA |
| | | | CDC5L |
| | | | AGAP1 |
| | | | TLN2 |
| | | | PCYOX1 |
| | | | ACAT1 |
| | | | YPEL5 |
| | | | DYNLT3 |
| | | | PKD1 |
| | | | DNASE2 |
| | | | RNMT |
| | | | SSC5D |
| | | | PCSK1N |
| | | | SLC17A5 |
| | | | C12orf75 |
| | | | SSPN |
| | | | NTN1 |
| | | | ANGPTL5 |
| | | | OMD |
| | | | AFF1 |
| | | | MARVELD1 |
| | | | RFNG |
| | | | VGLL4 |
| | | | TSPAN8 |
| | | | CBR3 |
| | | | ENSG00000263620 |
| | | | CTDSP1 |
| | | | SYNE2 |
| | | | PLAAT3 |
| | | | SMS |
| | | | KIDINS220 |
| | | | C1orf21 |
| | | | BOC |
| | | | PRG4 |
| | | | SNX21 |
| | | | SLPI |
| | | | LINC00963 |
| | | | HSD17B11 |
| | | | AGTR1 |

| | | | |
|---|---|---|---|
| | | | H2AJ |
| | | | PSD3 |
| | | | HSD17B14 |
| | | | RB1CC1 |
| | | | CILP |
| | | | RORA |
| | | | GFRA1 |
| | | | CACUL1 |
| | | | MEGF6 |
| | | | SP5 |
| | | | SNX29 |
| | | | PAMR1 |
| | | | JUP |
| | | | PRUNE2 |
| | | | SPOCK1 |
| | | | GLT8D1 |
| | | | MCOLN3 |
| | | | PET100 |
| | | | TMEM243 |
| | | | MIR99AHG |
| | | | BLOC1S1 |
| | | | KAT6B |
| | | | SVBP |
| | | | HPS1 |
| | | | FN3K |
| | | | LINC01315 |
| | | | PHYH |
| | | | RAB32 |
| | | | SMIM14 |
| | | | SFXN3 |
| | | | PBRM1 |
| | | | HPGD |
| | | | IDS |
| | | | TNRC6B |
| | | | FGFR1 |
| | | | CPQ |
| | | | LGMN |
| | | | CYP20A1 |
| | | | EAPP |
| | | | MBNL2 |
| | | | BBLN |
| | | | PBX1 |
| | | | TULP3 |
| | | | DEXI |
| | | | MMP24OS |
| | | | ZDHHC7 |
| | | | IST1 |
| | | | MYH10 |
| | | | FAM180B |
| | | | ABCA9 |
| | | | MARCHF2 |
| | | | ARHGEF12 |
| | | | HNMT |
| | | | CNPPD1 |
| | | | SNTA1 |
| | | | ANGPTL2 |
| | | | SNAP23 |
| | | | ABHD12 |

| | | | |
|---|---|---|---|
| | | | THRA |
| | | | CARD19 |
| | | | FKBP9 |
| | | | PDLIM3 |
| | | | UST |
| | | | KLHDC2 |
| | | | GLIPR2 |
| | | | PINK1 |
| | | | TRAK2 |
| | | | PKN1 |
| | | | ANGPTL1 |
| | | | SH3GLB2 |
| | | | EPB41L3 |
| | | | PIP4P2 |
| | | | SESTD1 |
| | | | AMDHD2 |
| | | | CCBE1 |
| | | | MYOF |
| | | | LIMS2 |
| | | | CMTM3 |
| | | | AKR1C2 |
| | | | KLF3 |
| | | | STX7 |
| | | | OLFML1 |
| | | | IRX5 |
| | | | ILK |
| | | | HSPB8 |
| | | | OSBPL1A |
| | | | PMEPA1 |
| | | | EIF4EBP3 |
| | | | PDLIM5 |
| | | | MTURN |
| | | | ECM2 |
| | | | PPP2R2A |
| | | | ZNF358 |
| | | | FAM102A |
| | | | C1QTNF7 |
| | | | GOLIM4 |
| | | | CCNDBP1 |
| | | | NIT2 |
| | | | SMIM20 |
| | | | CCDC50 |
| | | | LRRC58 |
| | | | LSM8 |
| | | | CCND1 |

| **Cluster shared DEgenes of fibroblasts** | |
|---|---|
| Up in BP lesional skin | Down in BP lesional skin |
| XRCC5 | DCD |
| LIMA1 | SCGB2A2 |
| CDC37 | MUCL1 |
| CCT3 | PDK4 |
| GNG5 | TXNIP |
| SNU13 | MAFF |
| MRPL51 | DNAJA1 |
| NRDC | TSC22D3 |
| MT-ND4L | CACYBP |

| | |
|---|---|
| S100A10 | CLK1 |
| EIF3B | ZFAND2A |
| LDHB | HBP1 |
| SSBP4 | MRPL18 |
| CCDC85B | HSPH1 |
| ALYREF | BAG3 |
| CDC42EP5 | UBL3 |
| GSPT1 | CRYAB |
| CNIH1 | OSR2 |
| FDPS | CKB |
| AK2 | DNAJB4 |
| SET | H2AZ1 |
| SSR1 | ID1 |
| PSMD6 | LDAF1 |
| GNB2 | PPP1R10 |
| PAXX | MAP1LC3B |
| ATP5F1D | MGP |
| SPCS2 | OSER1 |
| NDUFS3 | MAFB |
| G3BP1 | RSRC2 |
| RPLP2 | TSC22D1 |
| SSBP1 | GADD45B |
| HNRNPA3 | HSPE1 |
| RAN | NEU1 |
| RAB5C | ELN |
| PSMA4 | SNHG5 |
| MAP4 | DDIT3 |
| KPNB1 | BTG1 |
| PSMC5 | ARL6IP1 |
| RPL13A | CHIC2 |
| HNRNPAB | RASD1 |
| ILF3 | IER5L |
| B4GALT1 | CITED2 |
| RPS2 | EPB41L4A-AS1 |
| PRMT1 | STMN1 |
| SNRPE | CAV1 |
| PLIN3 | PPP1R15A |
| DNAJC15 | SNHG8 |
| TUBA1B | RNASE4 |
| HMGN1 | RSRP1 |
| VCP | DNAJB6 |
| ETFA | GEM |
| LY6E | TCP1 |
| SRI | ZFAS1 |
| MXRA5 | ZNF106 |
| ADRM1 | HSP90AB1 |
| PSMD13 | MYLIP |
| POLR2E | DNAJB1 |
| UGCG | HSPA1B |
| PTGES | NUDT4 |
| SSU72 | RSL24D1 |
| LMNA | HNRNPH3 |
| SRSF9 | UBE2S |
| KLF2 | RHOB |
| TUFM | HSPA1A |
| EDF1 | UBE2B |
| BANF1 | TIMP3 |

| | |
|---|---|
| PA2G4 | DDX6 |
| UBE2L3 | DANCR |
| MPZL1 | SERTAD1 |
| ARCN1 | CEBPB |
| UBE2M | UBB |
| CYC1 | ARID4B |
| PTMS | CDKN1A |
| PPM1G | UBC |
| POLR2G | TNFAIP6 |
| SNRPD3 | IRF1 |
| LRRC59 | TFPI |
| EVA1B | EIF4A2 |
| RPS17 | SLC38A2 |
| CAPRIN1 | SELENOP |
| TRIM47 | CCNL1 |
| CIAO2B | PER1 |
| PRRX1 | BTG2 |
| MYL12B | ZFAND5 |
| SNRPF | RPS27 |
| PSENEN | PNRC1 |
| COL5A2 | MAP1B |
| KRTCAP2 | BRD2 |
| IMPDH2 | JUN |
| COPB2 | ADIRF |
| DRAP1 | ANKRD12 |
| CLTC | SAP18 |
| TUBB | GPBP1 |
| SDF4 | SELENOK |
| TWIST2 | RPL30 |
| EBPL | RNF13 |
| DENR | SRSF3 |
| ATP5F1A | CCNI |
| EIF3I | ID3 |
| RER1 | ID2 |
| SNX17 | VEGFB |
| SURF4 | CD9 |
| ARF3 | COPS2 |
| RAD23B | PID1 |
| CCT7 | GPNMB |
| PSMC3 | SYF2 |
| PSMA3 | JUNB |
| NDUFB2 | ST13 |
| MT-ATP8 | EIF3E |
| METRNL | PNRC2 |
| SERPINH1 | RPL35A |
| IFI35 | VAT1 |
| TIMM10 | ARL4D |
| CTDSP2 | SLC3A2 |
| PSMD3 | CHASERR |
| EPRS1 | SRSF5 |
| STING1 | EIF1B |
| PUF60 | GLUL |
| ANAPC11 | KLF4 |
| ACTN4 | KLF9 |
| NOP10 | GSN |
| PRKCSH | ANKRD10 |
| MRPS34 | EIF1 |
| LMAN1 | SNHG29 |

| | |
|---|---|
| STOML2 | ARID5B |
| BGN | BNIP3L |
| ITGAV | WSB1 |
| PSMA6 | C12orf57 |
| LSM7 | MYC |
| RSU1 | RPL5 |
| PSMB2 | JMJD1C |
| SPCS1 | FHL1 |
| COL5A1 | RPL34 |
| RAB5IF | CIRBP |
| MRPS15 | RPS13 |
| LAMTOR5 | INTS6 |
| PPDPF | DDX5 |
| NDUFS5 | HERPUD1 |
| EIF5A | ELF2 |
| STT3A | SERINC1 |
| RPS26 | DST |
| CTSL | LTBP4 |
| SDHC | EPC1 |
| CCT5 | SBDS |
| EIF2S2 | CYBRD1 |
| MYL6 | RPL39 |
| APRT | S100A13 |
| TPM3 | DDIT4 |
| ATP5MC3 | BTG3 |
| RANBP1 | GAS6 |
| STAT3 | CD55 |
| DHRS4 | EPHX1 |
| TSPAN3 | CDKN1C |
| CANX | RPS15A |
| ERCC1 | NRN1 |
| PPIA | RBBP6 |
| COX8A | N4BP2L2 |
| POLR2F | DDAH2 |
| PSMD7 | RPL32 |
| POLD2 | RPL29 |
| YWHAB | TAF7 |
| PARK7 | CRNDE |
| SRPX | KMT2E |
| TMED10 | BDH2 |
| HNRNPR | CAMLG |
| FSTL1 | APOD |
| PSMB1 | RPL41 |
| SNRPD1 | CSRNP1 |
| MSN | RPL11 |
| SH3BGRL3 | TUBA1A |
| CLPTM1 | NR3C1 |
| TMED3 | NOP53 |
| GRHPR | SNHG7 |
| GSTO1 | SOX4 |
| PSMC4 | RPS3A |
| SERF2 | THBS2 |
| PPT1 | SPSB3 |
| COX5A | NACA |
| IFT57 | SOD1 |
| AP2M1 | ZBTB20 |
| MRPL37 | STK24 |
| AHCY | RPL9 |

| | |
|---|---|
| RBX1 | POLR1D |
| PDCD5 | DAZAP2 |
| EIF4G1 | LOX |
| PRKDC | RNF130 |
| COPZ1 | CEBPD |
| MT1E | PRELP |
| XRN2 | MAN1A1 |
| MRPL41 | KRT1 |
| MRPL4 | HLA-E |
| TOMM5 | CD302 |
| PSMD8 | NDFIP1 |
| TBCB | ETS2 |
| HSBP1 | PCBP2 |
| NDUFAB1 | RPS4X |
| CALU | TGIF1 |
| REXO2 | ANAPC16 |
| PSMB5 | EIF3L |
| TMEM165 | ANXA4 |
| TXN2 | RND3 |
| DDOST | UBXN1 |
| NUCB2 | RPL10 |
| MT-CO1 | H1-0 |
| ANPEP | OAZ1 |
| ALDH9A1 | |
| COX16 | |
| PSMD4 | |
| CAPNS1 | |
| RNASEH2C | |
| NDUFB3 | |
| PARVA | |
| GALK1 | |
| MRPL14 | |
| NDUFAF8 | |
| LAPTM4B | |
| PPA1 | |
| EFEMP1 | |
| MRPL11 | |
| AURKAIP1 | |
| EIF6 | |
| UQCRQ | |
| TIMM13 | |
| PDAP1 | |
| VDAC1 | |
| RUNX1 | |
| WDR1 | |
| NDUFA6 | |
| NME2 | |
| TPI1 | |
| CAP1 | |
| DNAJB11 | |
| CALR | |
| IMP4 | |
| ACTR2 | |
| ACTR3 | |
| AEBP1 | |
| PPP4C | |
| GADD45GIP1 | |
| DPM2 | |

| |
|---|
| AP2S1 |
| HBB |
| COMMD1 |
| DNPH1 |
| CNIH4 |
| OSTC |
| NAA10 |
| RBMS1 |
| SEC61G |
| COPS6 |
| TUBB6 |
| PRRX2 |
| FSTL3 |
| SPON1 |
| NME4 |
| ISOC2 |
| CCT6A |
| TMEM208 |
| MDH2 |
| NSFL1C |
| CIAO1 |
| PKM |
| LITAF |
| SIGMAR1 |
| CNN2 |
| AGTRAP |
| GAPDH |
| ETHE1 |
| MTLN |
| PSMD2 |
| GTF2I |
| LDHA |
| GPI |
| HSP90B1 |
| TRAPPC5 |
| PSMA5 |
| CYBA |
| ACTB |
| MRPL55 |
| GARS1 |
| MRPS16 |
| PSMB6 |
| CD63 |
| VAMP5 |
| CALM3 |
| SEC13 |
| NHP2 |
| RAB13 |
| CYBC1 |
| FKBP10 |
| SSNA1 |
| PSMB4 |
| PSMB10 |
| SAR1A |
| PLEC |
| FAM174C |
| TM9SF1 |
| HM13 |

| |
|---|
| COL6A3 |
| TXNDC15 |
| FIBP |
| LGALS1 |
| C1QBP |
| FAAP20 |
| PFN1 |
| PSMA7 |
| EFEMP2 |
| MT-ND6 |
| TUBA1C |
| TAF10 |
| S100A11 |
| MRPL20 |
| BAX |
| SPCS3 |
| HSPA5 |
| ZNRD2 |
| C1orf122 |
| PGAM1 |
| MAD2L2 |
| MRPL52 |
| CCDC124 |
| PPP1R14B |
| VCL |
| COL6A2 |
| ECHS1 |
| COL6A1 |
| NDUFS6 |
| ATP5MC1 |
| TWF1 |
| LSM12 |
| TXNL4A |
| PRDX4 |
| RPN2 |
| FLOT1 |
| SRA1 |
| COPS7A |
| SSR3 |
| FBN1 |
| SEC61B |
| LMAN2 |
| MRPS7 |
| VTI1B |
| CTSA |
| CYP51A1 |
| NDUFS8 |
| SEC61A1 |
| TKT |
| FGF7 |
| TRIM44 |
| POLR2L |
| TMEM147 |
| FKBP1A |
| PSMB9 |
| P3H1 |
| CFL1 |
| NUTF2 |

| |
|---|
| MRPL3 |
| SSR4 |
| SRPRA |
| COA3 |
| TPM4 |
| LAMTOR2 |
| JOSD2 |
| RPN1 |
| FKBP2 |
| TIMM8B |
| PSME2 |
| ARHGDIA |
| TNFRSF1A |
| PRELID1 |
| TXNDC5 |
| TRAPPC1 |
| CISD3 |
| VCAN |
| CHID1 |
| SNRPC |
| MACROH2A1 |
| PDIA6 |
| TMEM141 |
| LTBR |
| HDDC2 |
| PPIC |
| PLOD3 |
| PSMB3 |
| ENO1 |
| MYDGF |
| PHPT1 |
| HSD17B10 |
| IL1R1 |
| PDIA3 |
| BOLA3 |
| SND1 |
| BICC1 |
| PPP1CA |
| SLC39A8 |
| PPIB |
| GTF3C6 |
| COL1A1 |
| CKAP4 |
| YIF1B |
| SHC1 |
| CTSB |
| EIF4EBP1 |
| MLEC |
| IFITM3 |
| PLOD1 |
| CRELD2 |
| FLNA |
| KDELR2 |
| TGFBR2 |
| ARPC5L |
| RRBP1 |
| YIF1A |
| JPT1 |

| |
|---|
| PHB |
| TMEM176A |
| MANF |
| RCN1 |
| LY6D |
| TMED9 |
| MIF |
| MRPL27 |
| CD248 |
| SRPRB |
| P4HB |
| MRPL24 |
| KRT5 |
| S100A16 |
| MVP |
| CD320 |
| PLAAT4 |
| SERPINE2 |
| PCOLCE |
| RARRES2 |
| PDIA4 |
| TXNDC17 |
| ELOVL1 |
| HIF1A |
| ZNF593 |
| SRM |
| GGCT |
| LSM4 |
| NUCB1 |
| IKBIP |
| ARPC1B |
| FAM20A |
| RCN3 |
| TYMP |
| KRT17 |
| FKBP11 |
| KRT14 |
| SDF2L1 |
| IFITM2 |
| FUOM |
| TIMP1 |
| UQCC2 |
| C1R |
| EMILIN1 |
| NME1 |
| OSMR |
| IL32 |
| FGL2 |
| IGFBP4 |
| THY1 |
| PDPN |
| S100A2 |
| TNC |
| KRT16 |
| KRT6A |
| S100A9 |
| S100A8 |

| Cluster unique upregulated DEgenes of pericytes in BP lesional skin | | |
|---|---|---|
| THY1$^{+}$ PC | ACTA2$^{+}$ PC | Inter-PC |
| OS9 | PHLDA1 | MT-ND4 |
| NUDT22 | NDUFV1 | MT-ND1 |
| FARP1 | CDK2AP2 | MT-ATP6 |
| CRK | KRT6B | UBE2R2 |
| SUMO3 | COL12A1 | GPRC5C |
| SSR2 | MLF2 | DDX18 |
| PTRHD1 | FRMD4A | CD164 |
| PSMB10 | KLF13 | TFG |
| CDC42EP2 | SLC25A4 | IMP3 |
| MESD | KLF2 | PURA |
| EIF2AK4 | ENDOD1 | DHX9 |
| GNAI2 | B2M | CD81 |
| PARVB | MAP3K7CL | SP100 |
| GTF2H5 | TUSC1 | VCL |
| GYG1 | AK1 | SPCS2 |
| RCN3 | NENF | SNRPA1 |
| COL5A3 | MYO1B | KLF6 |
| MMP14 | CARHSP1 | PHF5A |
| CKAP4 | TRAM1 | PSMA1 |
| IMMT | CRIP1 | RAB20 |
| NUCB2 | SUSD2 | AKAP13 |
| CAPN1 | SOD3 | SPCS1 |
| PWP1 | ZFHX3 | HMGB1 |
| LAMC1 | LDHA | BCCIP |
| SLX9 | LHFPL6 | THBD |
| DLGAP4 | H2AJ | SERBP1 |
| CD320 | CLTC | SF3B6 |
| TMEM141 | ACTG2 | DGUOK |
| CHCHD5 | CSPG4 | REEP5 |
| SH3PXD2A | KCNMB1 | TRA2A |
| PCDH18 | PHLDA3 | GPATCH4 |
| SLC44A2 | MAP1B | ANAPC5 |
| SRP72 | ATP1A1 | HBB |
| PPIF | S100A14 | MMADHC |
| ARHGAP1 | NTN4 | GNB1 |
| ABL2 | IDH3G | SF3B4 |
| PCED1A | TXN | HNRNPR |
| MRPS2 | PPP2CB | BCL2L1 |
| PDHB | DMPK | RPL7 |
| DAP3 | EPHX1 | VDAC3 |
| GALK1 | SLC25A5 | MKNK2 |
| BACE2 | FOXS1 | QTRT1 |
| FMOD | PTTG1IP | RPL35 |
| ECEL1 | NOL3 | CLTA |
| CTSK | PAK2 | GPAT2 |
| COPZ1 | EIF4G2 | POLR2K |
| POSTN | | POLR2I |
| NDUFA8 | | TPM1 |
| THY1 | | KRAS |
| DPM2 | | VDAC2 |
| GLRX5 | | KTN1 |
| WARS1 | | CSNK1E |
| ERGIC3 | | STOM |
| ARFRP1 | | LRRFIP2 |
| CSNK2A1 | | GHITM |

| | | |
|---|---|---|
| CTDNEP1 | | SSBP1 |
| PLXND1 | | UQCRC2 |
| RRAS | | PAM16 |
| PLEKHB2 | | RPS20 |
| IFI6 | | NIFK |
| TNIP2 | | NDUFB4 |
| MARCKS | | ARL6IP5 |
| FAM174C | | CENPX |
| LARS1 | | NDUFAF4 |
| SIGMAR1 | | KDM2A |
| UXT | | ZC3H15 |
| ARRDC4 | | NDUFB1 |
| JOSD2 | | ADH5 |
| NECTIN2 | | LSM5 |
| SMIM10L1 | | CYSTM1 |
| SEC61A1 | | AKAP12 |
| IFNGR2 | | COPS8 |
| EFEMP2 | | SMTN |
| BCAR1 | | COX4I2 |
| ARHGAP17 | | EIF4H |
| TAX1BP3 | | STRAP |
| MSRB2 | | GPR4 |
| LINC01140 | | EI24 |
| CDC123 | | DDX46 |
| IKBIP | | RNF7 |
| CSRP1 | | GABARAPL2 |
| GPAA1 | | SNRPA |
| TAPBP | | UFM1 |
| ANKLE2 | | PRPF8 |
| MCTS1 | | STARD7 |
| RHOG | | ANKRD11 |
| MAP7D1 | | ATP5MC2 |
| TPPP3 | | SNHG16 |
| MTX1 | | RAB11A |
| DNMT1 | | RAB11B |
| PIM3 | | DUT |
| GET3 | | TMBIM1 |
| AAMP | | POLR2H |
| OLFML2B | | NT5C |
| ATP5F1A | | C4orf3 |
| ELAVL1 | | SF1 |
| COL5A2 | | EDNRB |
| ATP1B3 | | VPS29 |
| TXNDC15 | | NSA2 |
| CRELD2 | | NR2F2 |
| COL14A1 | | RELL1 |
| DNAJB11 | | NRBP1 |
| ELOVL1 | | BCL7C |
| RNASEK | | NDUFA11 |
| PAPSS2 | | XRN2 |
| B3GAT3 | | RNPS1 |
| HK1 | | NONO |
| ABCF1 | | BCAP31 |
| C4orf48 | | MT-ND4L |
| RFTN1 | | SEPTIN2 |
| PDLIM2 | | RASD1 |
| CREB3 | | ATP6V0C |
| LASP1 | | COX6C |

| | | |
|---|---|---|
| DCTN1 | | MRTFA |
| SMAP2 | | CCDC47 |
| UBE2L6 | | HDAC2 |
| CHN1 | | RWDD1 |
| M6PR | | RPL36A |
| PYCARD | | SRP9 |
| NINJ1 | | EIF5B |
| RCN1 | | DEGS1 |
| CMTM6 | | RPP21 |
| CTSD | | GTF2I |
| RPL22L1 | | SUMO1 |
| REM1 | | DDX24 |
| LYPLA1 | | KCTD20 |
| IPO7 | | ATP5PF |
| GPX1 | | XRCC5 |
| SPCS3 | | SYNPO2 |
| VSIR | | SNX3 |
| EFHD2 | | RPS29 |
| ITM2C | | EWSR1 |
| EIF3I | | EIF2S1 |
| POLR2G | | EMD |
| EIF4E2 | | NDUFC1 |
| SGIP1 | | IQGAP1 |
| EGFLAM | | FIS1 |
| DCTD | | SPTAN1 |
| LTBP3 | | DNAJA2 |
| DBNDD2 | | S100A6 |
| NANS | | AFF4 |
| CD4 | | IL6ST |
| CD82 | | TCEA1 |
| MPV17 | | C11orf98 |
| GSDMD | | RTRAF |
| VTI1B | | SRRT |
| NUBP2 | | SSB |
| MSC | | SF3B2 |
| LSM1 | | HNRNPD |
| SLC25A11 | | SDHD |
| USE1 | | NFIC |
| CD40 | | IRAG1 |
| C1S | | SEC31A |
| TMEM70 | | TERF2IP |
| METTL9 | | ETF1 |
| LAMP1 | | MT-ND2 |
| RELB | | RCSD1 |
| SMARCB1 | | VMP1 |
| CHMP3 | | MT-ND5 |
| SLC25A6 | | ARHGEF17 |
| CTTN | | HES4 |
| PPIC | | EMC4 |
| WNT6 | | MEA1 |
| FAM110B | | CAST |
| NDN | | BOP1 |
| C1QTNF5 | | NDUFA2 |
| MAPRE1 | | ADAMTS9 |
| PARP1 | | AXL |
| PLIN3 | | HSPA4 |
| DUSP23 | | PSMG2 |
| MRPL13 | | EIF3A |

| | | |
|---|---|---|
| ETFB | | RPS27L |
| NSMCE1 | | WTAP |
| FGF7 | | PAFAH1B1 |
| C14orf119 | | SOCS1 |
| SDC2 | | VASN |
| MRPL15 | | EID1 |
| MCUR1 | | SRRM2 |
| LOXL2 | | ERH |
| MRPL28 | | SARS1 |
| WDR13 | | RTN4 |
| COPZ2 | | RAB2A |
| SRPX | | SNRPB |
| CCL19 | | FYTTD1 |
| DNM2 | | ZBTB7A |
| IER3IP1 | | PITPNC1 |
| TEAD4 | | RASSF1 |
| MTCH2 | | SYNGR2 |
| LIMS1 | | DLC1 |
| CAMK1 | | C9orf78 |
| MAD2L2 | | SUPT4H1 |
| ATG3 | | DYNC1H1 |
| PEF1 | | NSD3 |
| MPZL1 | | WDR74 |
| TMEM50A | | PGF |
| ATRAID | | PRKAR1A |
| SRPRB | | ATP6V1G1 |
| SDHB | | THRAP3 |
| GUCY1B1 | | URI1 |
| COL5A1 | | COX7A1 |
| BASP1 | | PAIP1 |
| MT1F | | PSMD12 |
| BATF3 | | AQP3 |
| TRIP6 | | OCIAD1 |
| SPPL2A | | RTF2 |
| MZT2A | | BSG |
| TMED3 | | DDX39A |
| PIGT | | TRIR |
| BAD | | EMP2 |
| OAF | | HINT1 |
| POR | | RPL38 |
| CHPF | | ARFGAP3 |
| NELFE | | HNRNPF |
| AKR1A1 | | BCAM |
| ANPEP | | NDUFA10 |
| DNPEP | | SELENOF |
| INAFM1 | | PTMA |
| NABP1 | | COX20 |
| USP22 | | CMPK1 |
| PROS1 | | ADIRF |
| SGCB | | GLUD1 |
| MAPKAPK2 | | EIF3M |
| UBA1 | | MAGOH |
| NR1H2 | | NDUFB10 |
| MRPS18C | | WASL |
| TPGS1 | | C1orf52 |
| ANTKMT | | BTF3L4 |
| HLA-F | | NME3 |
| CCL26 | | PCMT1 |

| | | |
|---|---|---|
| TNIP1 | | ACADVL |
| METRN | | BDP1 |
| PDLIM4 | | NOL7 |
| SERP1 | | ARL5A |
| CDIPT | | UBE2D2 |
| PET100 | | HNRNPA3 |
| PFKL | | GSPT1 |
| OSTF1 | | |
| COL6A3 | | |
| CHID1 | | |
| MVB12A | | |
| RDH5 | | |
| UQCC3 | | |
| AOC3 | | |
| VPS25 | | |
| BLOC1S1 | | |
| B3GNT2 | | |
| CLSTN3 | | |
| TMEM176A | | |
| NPDC1 | | |
| TMEM219 | | |
| RBP5 | | |
| FERMT2 | | |
| UBE2N | | |
| CDC42EP5 | | |
| SSR1 | | |
| PSENEN | | |
| PSMB9 | | |
| ARCN1 | | |
| ISOC2 | | |
| PSMD9 | | |
| EPN1 | | |
| RAB32 | | |
| NDUFA4 | | |
| DAPK3 | | |
| CD276 | | |
| SLC25A1 | | |
| GLRX3 | | |
| SYNGR1 | | |
| LRPAP1 | | |
| OGFOD3 | | |
| SLC2A4RG | | |
| NECAP2 | | |
| DECR1 | | |
| NAA50 | | |
| SNX6 | | |
| IL6 | | |
| LAPTM4B | | |
| ITPRIPL2 | | |
| ACBD6 | | |
| MLST8 | | |
| TMEM167A | | |
| HAS2 | | |
| RNF187 | | |
| DBNL | | |
| PSMD3 | | |
| RBCK1 | | |
| MRPL47 | | |

| |
|---|
| RTL8B |
| STIP1 |
| CDC42 |
| MKKS |
| AKR7A2 |
| MRTO4 |
| TMEM208 |
| PSMG3 |
| ITPA |
| PEMT |
| EIF3K |
| GPM6B |
| TRAPPC4 |
| BMP1 |
| COPS7A |
| CAVIN2 |
| UROD |
| NIBAN2 |
| SHMT2 |
| TPD52L2 |
| IFI35 |
| PLTP |
| SLC39A1 |
| DMAC1 |
| NCOA7 |
| SEMA4C |
| CTSA |
| FPGS |
| CIAO1 |
| TRAP1 |
| IL32 |
| FKBP9 |
| CCL21 |
| PPP1R7 |
| AP2A1 |
| SND1 |
| RAB24 |
| TIMM50 |
| DPH3 |
| DAZAP1 |
| COMMD4 |
| FIBP |
| CLEC11A |
| APCDD1 |
| ERLEC1 |
| WDR61 |
| PARVA |
| SNAI2 |
| FAM114A1 |
| PTGES |
| BHLHE40 |
| ETHE1 |
| MFSD10 |
| TRPC6 |
| FBXW5 |
| RPL26L1 |
| CYP7B1 |
| ACTR1A |

| | | |
|---|---|---|
| FLYWCH2 | | |
| TGFB1I1 | | |
| PRKCSH | | |
| COL23A1 | | |
| TXNDC5 | | |
| ADAMTS12 | | |
| NFU1 | | |
| MYOF | | |
| CDH13 | | |
| COX16 | | |
| RELA | | |
| NEURL1B | | |
| TRIM44 | | |
| RGS10 | | |
| METRNL | | |
| ADAMTS2 | | |
| ARL3 | | |
| ERP44 | | |
| SYAP1 | | |
| DEF8 | | |
| PDIA4 | | |
| MRPS35 | | |
| ITGAV | | |
| PLSCR3 | | |
| TMEM176B | | |
| PLOD1 | | |
| EMC6 | | |
| ZNF511 | | |
| ITGAE | | |
| PDZD11 | | |
| HYOU1 | | |
| MRPL16 | | |
| TRADD | | |
| SFT2D1 | | |
| PPT1 | | |
| SEPTIN11 | | |
| FKBP10 | | |
| URM1 | | |
| EHD1 | | |
| SCARB2 | | |
| MCFD2 | | |
| CERCAM | | |
| PEPD | | |
| ARL1 | | |
| RRP36 | | |
| AKT1 | | |
| RALA | | |
| PSMD1 | | |
| FSTL1 | | |
| **Cluster unique downregulated Degenes of pericytes in BP lesional skin** | | |
| $THY1^+$ PC | $ACTA2^+$ PC | Inter-PC |
| IFNGR1 | PDE4DIP | PAF1 |
| EIF4B | OSBPL9 | CLU |
| LACTB | AKAP1 | DNAJB9 |
| SORBS3 | WTIP | CCNT1 |
| CXCL3 | NIBAN1 | RPL6 |
| ISCU | MAP3K8 | ALKBH7 |

| | | |
|---|---|---|
| KRT1 | TMEM14A | LRIF1 |
| APPL1 | HSDL2 | DEDD2 |
| FAM241A | RPS21 | MOB2 |
| S100A13 | FHL5 | NXT1 |
| MPHOSPH8 | CDS2 | ZNF302 |
| FKBP5 | MPC1 | ILF2 |
| ID1 | PPP1CB | CDKN2AIP |
| IGFBP2 | CRIM1 | IGFBP5 |
| STARD13 | PLS3 | ZMYM5 |
| PCM1 | PPP1R12A | CCT4 |
| SEPTIN10 | NAMPT | ID2 |
| PTEN | TBX2-AS1 | RBM25 |
| KANK2 | SORT1 | PPP1R15B |
| SLC1A5 | RBM39 | TMEM59 |
| SMIM3 | PJA2 | NUPR1 |
| RBM3 | FBXO32 | HEXIM1 |
| TM4SF1 | PICALM | RND1 |
| REV3L | CASQ2 | FDFT1 |
| TCIM | PRKG1 | ACKR3 |
| SAV1 | SORBS1 | AMD1 |
| SAT2 | ZRANB2 | GSTM3 |
| TRNP1 | SELENBP1 | MAGED2 |
| CD46 | HNRNPC | ZNF181 |
| PIP | CABIN1 | TOB1 |
| TBC1D1 | JMJD1C | HBEGF |
| RPL39 | CNRIP1 | ID3 |
| MOCS1 | C12orf75 | ZNF331 |
| CXCL14 | SNHG6 | NARF |
| HMOX2 | SDCBP | RPS3 |
| NDUFA5 | CARMN | CBX4 |
| HIPK3 | PHLDB2 | CALM1 |
| TMEM204 | UGP2 | H2BC11 |
| SCGB1D2 | NFIA | FAM117A |
| AHNAK | MBNL1 | FXR1 |
| CLEC3B | SSBP2 | TUBA1A |
| PALLD | IDS | HOOK2 |
| XBP1 | ATRX | RPL18 |
| AIMP1 | TACC1 | MT1M |
| ESD | FUS | ZC3H12A |
| PSMC6 | THRA | RPS9 |
| ITM2B | TWIST1 | EEF1D |
| SPRY2 | TTLL7 | TRIP10 |
| RPAIN | CAMK2G | ANP32E |
| EBF1 | ZBTB38 | PTTG1 |
| PCNP | | YY1 |
| NUFIP2 | | RPL19 |
| NGDN | | MGST3 |
| TCEAL3 | | GPCPD1 |
| CALCOCO2 | | CHTOP |
| PABPC1 | | ARF4 |
| F10 | | SLC25A33 |
| PER1 | | COQ10B |
| VAT1 | | PRXL2C |
| CPNE3 | | NR4A3 |
| CDC42EP4 | | NASP |
| SAMHD1 | | EIF3F |
| PARM1 | | NEDD9 |
| OLFM2 | | ANKRD37 |

| | | |
|---|---|---|
| TGFB1 | | ABHD5 |
| DYNLT3 | | HNRNPA2B1 |
| PSIP1 | | MID1IP1 |
| MFF | | PIGA |
| CD302 | | TUBB2A |
| DCN | | PMP22 |
| DARS1 | | DDAH2 |
| ADAMTS1 | | RPS19 |
| DHX15 | | PLEKHA3 |
| NAP1L1 | | TAF9 |
| SLC9A3R2 | | PCF11 |
| GJC1 | | MEF2D |
| ZNF24 | | SLFN11 |
| CNBP | | RAB5A |
| DCXR | | ZC3HAV1 |
| MRPS36 | | NXF1 |
| 3620 | | NMB |
| COPS2 | | H2AZ2 |
| FNBP4 | | CAV2 |
| ROCK2 | | IVNS1ABP |
| RBBP4 | | RPS24 |
| MTHFD2 | | BCL3 |
| PPP1R2 | | MSX2 |
| GNG11 | | NRARP |
| GNAI1 | | PHB2 |
| INTS6 | | CHD2 |
| PIK3R1 | | COMMD6 |
| VAMP3 | | GABARAPL1 |
| PELO | | PRPH |
| SCGB1B2P | | TIPARP |
| AP3S1 | | UBE2A |
| CYP26B1 | | SAP30 |
| DEK | | TWSG1 |
| DTWD1 | | HNRNPU |
| APOD | | ZFAND2A |
| TJP1 | | EIF3D |
| TNS2 | | PRMT9 |
| ECI1 | | RPL21 |
| SPTBN1 | | CD55 |
| ADH1B | | BMP2 |
| BRI3 | | KLF4 |
| SERINC3 | | SPAG9 |
| RSL1D1 | | INSIG1 |
| PKIG | | EFNA1 |
| MEF2A | | CHMP5 |
| HNMT | | RPL7A |
| POLR1D | | MXI1 |
| ADRA2A | | RPS18 |
| SASH1 | | TAMALIN |
| SAP30BP | | CNN1 |
| DEPP1 | | HSPB1 |
| ANP32A | | RGS2 |
| FHL1 | | BCAS2 |
| OAT | | DUSP5 |
| ADGRF5 | | KLF5 |
| EPN2 | | TRIM28 |
| CD74 | | ALAS1 |
| CFD | | HSPB8 |

| | | |
|---|---|---|
| RSBN1L | | NAF1 |
| CTDSPL | | ELOVL5 |
| NPTX2 | | TMED5 |
| ABCC9 | | MRPL32 |
| FAM162B | | PPP2R2A |
| GINM1 | | GNAS |
| HP1BP3 | | HMGN2 |
| CCDC71L | | CCNG2 |
| RNF130 | | PRDX6 |
| STK24 | | |
| UBE2E1 | | |
| SCAF11 | | |

| **Cluster shared DEgenes of pericytes** | |
|---|---|
| Up in BP lesional skin | Down in BP lesional skin |
| SERPINB6 | CLK1 |
| CD151 | RPL30 |
| IGFBP7 | HES1 |
| ARPC2 | RPS27 |
| CCT3 | ZFAND5 |
| LRRFIP1 | EIF3L |
| PSMB6 | NR4A2 |
| UFD1 | PNRC1 |
| EHD2 | GLUL |
| NME2 | RPL24 |
| CALU | CLDND1 |
| TPM4 | HSP90AA1 |
| KRT6C | UBE2S |
| CAPZB | IRF1 |
| RPL13A | DDIT4 |
| SNU13 | EIF3E |
| MRPL14 | TSC22D3 |
| CSTB | JUNB |
| PLAC9 | ETS2 |
| NDUFA6 | RGS16 |
| CDV3 | KLF9 |
| MLEC | SQSTM1 |
| RPL28 | N4BP2L2 |
| UQCRC1 | RPL5 |
| NDUFA1 | TXNIP |
| TMEM256 | MCL1 |
| RAMP1 | BNIP3L |
| TNC | SNHG1 |
| MRPS34 | GADD45B |
| GRINA | NFIL3 |
| RAN | IER2 |
| MRPL3 | HNRNPH1 |
| S100A4 | PRNP |
| SCAND1 | HSP90AB1 |
| HRAS | MYC |
| COX5A | LDAF1 |
| FKBP1A | RCAN2 |
| CLPP | MIR22HG |
| SERPINF1 | RPL22 |
| MRPL24 | NBEAL1 |
| CD59 | RPL35A |
| NT5DC2 | UBC |

| | |
|---|---|
| MRPS26 | PGRMC1 |
| MSN | HBP1 |
| PSMB1 | CKB |
| CALR | NOP58 |
| TIMM13 | KMT2E |
| FN1 | SLC38A2 |
| ARF5 | GLA |
| EPRS1 | ZFP36 |
| POLR2E | EIF4A3 |
| AP2S1 | BRD2 |
| COL3A1 | ZFAS1 |
| RAD23A | ATF3 |
| C20orf27 | RSRP1 |
| GADD45GIP1 | ZFP36L2 |
| LMAN1 | EPC1 |
| S100A2 | BCL6 |
| DCTN3 | RPS10 |
| GRHPR | PPP1R10 |
| DRAP1 | FAU |
| ROMO1 | HLA-E |
| UQCRFS1 | YTHDC1 |
| GPI | IER5L |
| RAB5C | IFI16 |
| PRDX5 | ARL6IP1 |
| SRI | SERTAD3 |
| COX6A1 | H2AZ1 |
| SRSF9 | MT1A |
| TUBB6 | RPL11 |
| RHOQ | EIF1B |
| PRMT1 | SNHG29 |
| TBCB | CIRBP |
| HM13 | FOSB |
| RNF181 | H3-3B |
| VAMP5 | CAMLG |
| SNRPG | MYLIP |
| CALD1 | ARHGAP10 |
| A2M | HSPE1 |
| GNB2 | YME1L1 |
| GTF3C6 | SERTAD1 |
| KRT16 | BCLAF1 |
| COL18A1 | SAP18 |
| PPP2R1A | MAT2A |
| KRT14 | UBA2 |
| COX8A | GEM |
| NAA10 | WSB1 |
| APOE | H4C3 |
| PARK7 | YPEL5 |
| RPS19BP1 | SUGT1 |
| SMOC2 | SRSF3 |
| PSMC3 | SRSF7 |
| MAP4 | HSPA2 |
| RAB7A | SEPHS2 |
| PSMB3 | HNRNPH3 |
| MTPN | MATR3 |
| RSU1 | PDK4 |
| LAMTOR2 | ISYNA1 |
| RNASEH2C | SAT1 |
| LAMTOR5 | NR4A1 |

| | |
|---|---|
| MYLK | NDRG2 |
| EIF6 | MT1X |
| LSM4 | RSL24D1 |
| HSD17B10 | ARIH1 |
| MYO1C | DNAJB6 |
| ATP5MC1 | HSPA8 |
| GNG10 | DUSP1 |
| ISG15 | SNHG12 |
| PCOLCE | GOLGB1 |
| MRPL57 | SNHG5 |
| MIR4435-2HG | PPP1R15A |
| SF3B5 | FTL |
| ARPC5 | PPIG |
| ALYREF | TCP1 |
| NME1 | GPBP1 |
| MRPS15 | RRAD |
| RNH1 | DDX3X |
| TALDO1 | PLK2 |
| AEBP1 | EIF1 |
| ARHGDIA | CEBPD |
| POLD2 | SOCS3 |
| MICOS10 | UBB |
| EHBP1L1 | FOS |
| PPP1R14B | CDKN1A |
| ANAPC11 | MAFF |
| TMEM258 | CCNL1 |
| NDUFB2 | RPL9 |
| RTL8C | H1-10 |
| COA3 | RSRC2 |
| NNMT | EIF4A2 |
| MACROH2A1 | CCNI |
| FKBP2 | DANCR |
| TMOD3 | TAF7 |
| EBNA1BP2 | CITED2 |
| S100A9 | RASL11A |
| MPG | UBE2B |
| POLR2L | SRSF5 |
| CCT8 | EIF5 |
| COPB2 | CSRP2 |
| COL4A2 | DSTN |
| MT-ND6 | DDIT3 |
| PPIB | DDX5 |
| APRT | CHASERR |
| MT-CO3 | SNHG8 |
| CRISPLD2 | C12orf57 |
| PSMC4 | CCL2 |
| DNAJC15 | ID4 |
| TAGLN | CEBPB |
| IFITM3 | GAS5 |
| PDLIM1 | DNAJA1 |
| PLEC | BTG1 |
| NDUFA12 | FXYD1 |
| SNRPF | RPL34 |
| PA2G4 | LAPTM4A |
| CCT7 | HSPD1 |
| CALM3 | KLF10 |
| CHCHD10 | CSRNP1 |
| SEC13 | GTF2B |

| | |
|---|---|
| IFITM2 | MAP1LC3B |
| NDUFS8 | MAFB |
| TOMM22 | DST |
| MRPL11 | DNAJB4 |
| SSU72 | BTG2 |
| CCDC124 | BAG3 |
| CYB5R3 | TSC22D1 |
| SMIM12 | RPS3A |
| SHISA5 | JUN |
| SH3BGRL3 | MAPRE2 |
| COX5B | LMOD1 |
| RPN2 | |
| CTNNB1 | |
| TNFRSF1A | |
| MRPS7 | |
| PSMA5 | |
| ACTA2 | |
| EVA1B | |
| PSMD8 | |
| ARPC3 | |
| TUBB | |
| SEC61G | |
| YIF1A | |
| SNRPE | |
| PGAM1 | |
| COPS6 | |
| PPIA | |
| NDUFB11 | |
| PSMB5 | |
| ATP5F1D | |
| PALM2AKAP2 | |
| VASP | |
| PPM1G | |
| PFN1 | |
| NDUFA13 | |
| MDH1 | |
| BANF1 | |
| NUCB1 | |
| RANBP1 | |
| NDUFS7 | |
| LY6E | |
| CYC1 | |
| C1QBP | |
| CCDC85B | |
| FABP5 | |
| NDUFS5 | |
| CAPNS1 | |
| KRT5 | |
| COL1A2 | |
| COL1A1 | |
| MRPL27 | |
| COX6B1 | |
| ANXA6 | |
| AK2 | |
| CNIH4 | |
| GUK1 | |
| MYL12A | |
| SSNA1 | |

| |
|---|
| CCT5 |
| ARL2 |
| CLIC1 |
| MRPL4 |
| SERF2 |
| TIMM8B |
| MRPL21 |
| TAGLN2 |
| SRPRA |
| COPS9 |
| HSBP1 |
| HSPB6 |
| NDUFAB1 |
| TWF1 |
| PUF60 |
| COPE |
| FAM50A |
| TMEM147 |
| BST2 |
| SNF8 |
| MRPL34 |
| OST4 |
| REXO2 |
| ADCY3 |
| SRM |
| CFL1 |
| CHCHD2 |
| ZNHIT1 |
| SPARC |
| GSTO1 |
| MRPL20 |
| C1orf122 |
| COL4A1 |
| FAAP20 |
| ATP6V0B |
| RRAGA |
| ZYX |
| PDIA3 |
| ATP5F1E |
| CSNK1A1 |
| NME4 |
| MT-CYB |
| TNFRSF12A |
| CIAO2B |
| HSP90B1 |
| PDGFRB |
| ATP6V0E1 |
| SLC25A39 |
| CYTOR |
| TMSB10 |
| TGFBI |
| ATOX1 |
| TRAPPC1 |
| KRT17 |
| EDF1 |
| STUB1 |
| NHP2 |
| CD63 |

| |
|---|
| PPA1 |
| PHB |
| SNRPD1 |
| MVP |
| ITGA7 |
| COL6A1 |
| TRAPPC5 |
| IMPDH2 |
| COL6A2 |
| CDK4 |
| TMED10 |
| PPP4C |
| NDUFAF3 |
| PSMD6 |
| SSR4 |
| TMED9 |
| SNRPD3 |
| CIB1 |
| DDOST |
| RPN1 |
| KRT6A |
| TSPO |
| LY6D |
| PKM |
| TKT |
| LITAF |
| MDH2 |
| LGALS1 |
| TXNDC17 |
| MYH9 |
| TXN2 |
| CCT6A |
| PPDPF |
| MICOS13 |
| UQCRQ |
| RAC1 |
| PHLDA2 |
| EIF5A |
| CAVIN3 |
| KDELR2 |
| SSR3 |
| P4HB |
| PSMD13 |
| RBX1 |
| NDUFB3 |
| EIF4G1 |
| NDUFA4L2 |
| UFC1 |
| CDK2AP1 |
| GTF3A |
| HIF1A |
| UBE2L3 |
| COA4 |
| NEDD8 |
| NOP10 |
| LGALS7B |
| MRPL36 |
| PFDN2 |

| |
|---|
| DAD1 |
| PSMD2 |
| ACTR3 |
| PSMA6 |
| RRBP1 |
| TUFM |
| PSME2 |
| CLTB |
| SEPTIN4 |
| RAD23B |
| ENO1 |
| MRPL41 |
| POLE4 |
| RAB31 |
| CTSB |
| SELENOW |
| MRPL51 |
| PSMB2 |
| HOPX |
| RAB13 |
| S100A8 |
| AP2M1 |
| LMNA |
| TPM3 |
| GAPDH |
| MIF |
| SDF4 |
| MYDGF |
| HLA-A |
| ANXA11 |
| TPI1 |
| MTLN |
| NDUFB7 |
| LGALS3BP |
| CTSZ |
| POLD4 |
| ATP5MC3 |
| PSMA7 |
| IFI27L2 |
| C19orf53 |
| HDLBP |
| ANXA2 |
| RABAC1 |
| AURKAIP1 |
| ATP5MF |
| FLOT1 |
| BGN |
| TOMM5 |
| PDCD5 |
| NDUFB6 |
| PDIA6 |
| RPS16 |
| PRELID1 |
| PHPT1 |
| COMT |
| SFN |
| STOML2 |
| MYL6 |

| |
|---|
| RPS17 |
| VDAC1 |
| KXD1 |
| PRDX4 |
| ACTB |
| HNRNPAB |
| SH3GLB1 |
| PPP1CA |
| POLR2J |
| ACTN1 |
| RPS26 |
| ERCC1 |
| S100A10 |
| MRPL52 |
| ARPC1B |
| RHOC |
| DBI |
| TIMP1 |
| GJA4 |
| CISD3 |
| PIN1 |
| PDAP1 |
| NDUFAF8 |
| TAF10 |
| ADRM1 |
| RPS2 |
| ACTG1 |
| UBL5 |
| METTL26 |
| CYBA |
| MCAM |
| HSPA5 |
| CAPN2 |
| SEC61B |
| UQCR10 |
| PSMD7 |
| MT-CO1 |
| SDF2L1 |
| RPLP2 |
| YWHAB |
| MIEN1 |
| CAP1 |
| BEX3 |
| SLIT3 |
| MCRIP1 |
| YWHAH |
| IGFBP4 |
| TXNL4A |
| NUTF2 |
| NDUFS6 |
| C1QTNF1 |
| SELENOH |
| OSTC |
| ATP6V1F |
| S100A11 |
| SNRPC |
| EMP3 |
| GSTP1 |

| |
|---|
| ELOB |
| CCND1 |

| Cluster unique upregulated Degenes of endothelial cells in BP lesional skin | | | | | | |
|---|---|---|---|---|---|---|
| LEC | Arterioles | PAC | PVC | PCV | Venules | CV |
| EMC3 | APLP2 | EFNB1 | SEPTIN9 | TRMT10C | SAE1 | PTRHD1 |
| PSMB10 | JAG2 | MLEC | CD164 | TMEM87B | ZNF467 | CHCHD5 |
| FAM43A | MIDN | NDRG1 | MESD | TFG | STK17A | CLPP |
| TMEM256 | UQCRC1 | TMEM205 | HSF1 | RC3H1 | POSTN | MRPL24 |
| AFAP1L1 | PSMA1 | VAPA | MRPS6 | KPNB1 | LTC4S | MMADHC |
| NRDC | SRP72 | APLNR | SH3TC1 | DEDD2 | MSRB2 | GLRX5 |
| SYNM | ZMIZ1 | TMEM88 | ORAI1 | TRA2A | FAM171A1 | GRHPR |
| IGFBP5 | RAD23A | MCF2L | SELENOS | ARID4A | FBN1 | FAM110D |
| DPM2 | FES | CPD | NUCB2 | BCL2L1 | SELENON | SNRPG |
| CDC25B | FBL | SPRR1B | PWP1 | FNDC3B | SSH2 | LARS1 |
| PGM5 | NOS3 | JCAD | SPCS1 | RND1 | EHBP1L1 | RHOG |
| PTBP1 | RASIP1 | H19 | RALGDS | IFNGR2 | MRPS18B | ALYREF |
| GPR146 | BCAP29 | RBBP4 | NDUFV1 | ENSG000002 74213 | VCAN | ECHS1 |
| PSMB7 | ASRGL1 | GLTP | TMEM54 | NDUFB4 | HHEX | TRAPPC2L |
| RPL35 | NAA20 | SBSN | MOSMO | GON4L | RFTN1 | CAPRIN1 |
| SMIM10L1 | DPP7 | TWF2 | NSFL1C | SLC38A2 | FBLN2 | PYCARD |
| SCAMP2 | LPCAT2 | APBB2 | DGUOK | RPS6KA3 | STARD7 | UBALD2 |
| ACKR3 | HCFC1R1 | C1orf54 | DOC2B | CEMIP2 | DHX30 | CNIH1 |
| BCAR1 | GNPTG | TOX4 | DNAJC7 | ADGRG1 | LTBP3 | CD40 |
| TC2N | PSMC4 | PEMT | LYPLA2 | SF1 | SLC25A24 | CTHRC1 |
| C1R | PTP4A3 | IFI27L2 | ATP6AP2 | CHD1 | TGFB1 | TMED2 |
| RNASEH2C | SLC6A6 | PHACTR2 | ITGB1BP1 | FILIP1L | RAB1A | HDGFL3 |
| COMMD1 | POLR2A | SENP6 | TNFRSF14 | PER1 | G6PC3 | SEC11C |
| PDPN | ABRACL | PSMA4 | ANP32B | H4C3 | GDI1 | COX7B |
| AEBP1 | CTNNB1 | CDC42EP1 | RAB6A | NOP56 | GSE1 | DENR |
| DPYSL3 | LSM3 | | SEPTIN10 | NDUFC1 | IRAK3 | NECAP2 |
| BRD4 | MRPL9 | | ABI3 | ZFC3H1 | ELN | TMEM167A |
| C4orf48 | HEY1 | | KLF11 | VGLL4 | DBNL | MKKS |
| RCN2 | PIK3C2A | | TNIP2 | ITPKC | UBE2V1 | CYTH1 |
| SH3BP5 | SMAP1 | | SPNS2 | NXF1 | FHL2 | ZNRF1 |
| DUSP6 | JAG1 | | SERTAD2 | IVNS1ABP | LTBP2 | GTF2A2 |
| ATP5MG | PELI1 | | CITED4 | GOLGB1 | FIBP | ANXA4 |
| NINJ1 | SYNPO | | COBLL1 | NRARP | ZBTB8OS | PIN1 |
| MRPL11 | EMD | | PTRH1 | TERF2IP | NFE2L1 | TRIM28 |
| SMIM12 | RAP1B | | RHOJ | CPEB4 | LIMS2 | TMEM160 |
| SH2D3C | LTBP4 | | CDK6 | PMAIP1 | LAMA4 | CCL23 |
| RPS9 | NCKAP1 | | RNF19B | MT-ND2 | MBTPS1 | MRPL55 |
| SNRPD2 | ALPL | | SEC61A1 | GJA1 | ACADVL | FAM162A |
| CCDC25 | NOTCH4 | | PLPP3 | BAG1 | PTGDS | |
| SULF2 | SIPA1L2 | | HSD17B10 | HOXD10 | | |
| RPL36A | NPTN | | ANKLE2 | SOX4 | | |
| SEMA3A | PSMD6 | | CD2AP | FERMT2 | | |
| MRPL21 | PSMG2 | | ARF3 | RPS27L | | |
| ETFB | EFNB2 | | LSM5 | KRT18 | | |
| UQCC2 | RAPGEF5 | | SZRD1 | ADAR | | |
| CISD1 | RHOA | | ANXA7 | YPEL2 | | |
| DCTPP1 | LEPROTL1 | | SLAMF1 | ZNF267 | | |
| RPLP1 | LYL1 | | GOLPH3 | PNN | | |
| DCXR | ATN1 | | UBQLN1 | RIOK3 | | |
| DNPH1 | RAB14 | | HK1 | RGS2 | | |
| UGP2 | B4GALT1 | | PDCD6IP | GLUD1 | | |
| MFAP2 | ATP13A3 | | RNF145 | HP1BP3 | | |

| | | | | |
|---|---|---|---|---|
| MZT2A | SHE | | PFKFB3 | PHACTR4 |
| SSB | GJA4 | | STX4 | TRIB1 |
| TMED3 | EMP3 | | AIP | ENSG00000289152 |
| DBN1 | | | RPN2 | MRPL32 |
| PFKL | | | FEZ2 | CCNL2 |
| KRT7 | | | UCK2 | |
| ISOC2 | | | VASP | |
| MRPL40 | | | PANK2 | |
| KCTD17 | | | VPS28 | |
| PSME2 | | | SLC25A37 | |
| TNXB | | | ACP1 | |
| EFEMP1 | | | PLCB4 | |
| STON2 | | | YPEL5 | |
| TRAPPC3 | | | NFKB1 | |
| CCL21 | | | HAPLN3 | |
| FLOT1 | | | PLAU | |
| CEBPB | | | TWF1 | |
| POLR2J | | | MCUR1 | |
| ETHE1 | | | SNHG6 | |
| RPL38 | | | CLEC14A | |
| UBTD1 | | | UBE2Z | |
| PTMA | | | SNF8 | |
| OLA1 | | | DNM2 | |
| CYBA | | | MTCH2 | |
| PAK2 | | | IRAK2 | |
| NUDT16 | | | LPP | |
| ATP5MJ | | | BATF3 | |
| MARCHF3 | | | ELMO1 | |
| | | | NFIC | |
| | | | CDC42BPB | |
| | | | TNIP1 | |
| | | | PIK3R1 | |
| | | | STUB1 | |
| | | | PDLIM4 | |
| | | | GGCT | |
| | | | PSENEN | |
| | | | NRP2 | |
| | | | CFI | |
| | | | PAFAH1B1 | |
| | | | RRP7A | |
| | | | LSM6 | |
| | | | CSF2RB | |
| | | | SETD5 | |
| | | | SHANK3 | |
| | | | SNX6 | |
| | | | CNKSR3 | |
| | | | PSMD2 | |
| | | | ADSS2 | |
| | | | BMP2 | |
| | | | MPRIP | |
| | | | ATXN2L | |
| | | | ARL6IP4 | |
| | | | MGAT4A | |
| | | | FKBP9 | |
| | | | RABAC1 | |
| | | | STK25 | |
| | | | MAP1LC3A | |

| | | | | | | |
|---|---|---|---|---|---|---|
| | | | PAIP1 | | | |
| | | | COMT | | | |
| | | | DAZAP1 | | | |
| | | | ABCD4 | | | |
| | | | ANP32A | | | |
| | | | ATP11A | | | |
| | | | KXD1 | | | |
| | | | ETFA | | | |
| | | | SBNO2 | | | |
| | | | PGM2 | | | |
| | | | MAP2K2 | | | |
| | | | TSPAN3 | | | |
| | | | CD44 | | | |
| | | | TSPAN4 | | | |
| | | | ARFGAP3 | | | |
| | | | FBXW5 | | | |
| | | | ACTR1A | | | |
| | | | EMC7 | | | |
| | | | EGFL7 | | | |
| | | | CYB5A | | | |
| | | | HECW2 | | | |
| | | | SSRP1 | | | |
| | | | EIF2S3 | | | |
| | | | SLC12A7 | | | |
| | | | ITGAV | | | |
| | | | ZNF326 | | | |
| | | | EIF4EBP1 | | | |
| | | | ARL4C | | | |
| | | | NUDCD1 | | | |
| | | | MAPK3 | | | |
| | | | TIMMDC1 | | | |
| | | | R3HDM4 | | | |
| | | | PSMD1 | | | |
| | | | ATP6V1F | | | |
| | | | UQCRB | | | |
| **Cluster unique downregulated Degenes of endothelial cells in BP lesional skin** | | | | | | |
| LEC | Arterioles | PAC | PVC | PCV | Venules | CV |
| CRACR2B | STARD13 | TXNDC12 | RPL7L1 | UBE2R2 | CRK | TM9SF2 |
| C11orf58 | PCM1 | SIVA1 | AHI1 | NDUFV2 | ZNF292 | ZBTB20 |
| PTGES3 | HEXIM1 | TCEAL4 | CDC42EP2 | RPL15 | EPSTI1 | ELK3 |
| DCTN2 | MAGED2 | GPIHBP1 | DNAJB9 | RPL17 | FKBP5 | MOB2 |
| GPM6A | FOXC1 | GNG12 | CNOT4 | STAT1 | ITSN2 | FNTA |
| PPP1CC | FAM124B | TMEM123 | DENND4A | SF3B4 | APH1A | ANKRD29 |
| FXYD6 | CX3CL1 | RPL37 | RGL2 | RIT1 | PARP12 | PLD3 |
| ADD3 | KMT2C | IGFBP3 | BCCIP | UXT | PRPF8 | LAMB2 |
| RBM39 | NARF | ENSA | PTPRE | CYSTM1 | BBX | RRAS |
| CALM1 | PLLP | YIPF3 | NIP7 | SECTM1 | PCMTD1 | TPM1 |
| MPG | EPC1 | MRPL22 | ERG | RPS3 | SEMA6A | KRAS |
| NT5E | BCL6 | TPM2 | YRDC | RPL13 | AKR1C1 | LFNG |
| PKHD1L1 | H2AX | PHC2 | CDC34 | DAB2 | SULF1 | SEC62 |
| TMEM35B | ZNF706 | FBXO31 | MAX | SENCR | LIMS1 | SERTAD4-AS1 |
| SNX2 | SERPINB1 | TIMP4 | SS18L2 | RPS25 | PLA1A | OXA1L |
| NTAN1 | SUGT1 | NAPA | ZNF644 | EMCN | MYRIP | BFAR |
| MAF | RTF1 | RBM17 | PRNP | ESD | SOD1 | UFM1 |
| STXBP6 | UAP1 | UQCRH | POLR2K | PRPF40A | CRTAC1 | PJA2 |
| LYVE1 | MXD4 | KRBA2 | NOL10 | XRN2 | BRI3 | VSIR |
| REEP3 | GNAQ | | CCDC59 | RNF167 | TMOD1 | DARS1 |

| | | | | | | |
|---|---|---|---|---|---|---|
| CHRDL1 | PRDX1 | | TEAD1 | CHTOP | CPE | PERP |
| ADAM10 | USP53 | | PRRC2B | TRIM69 | RNF10 | ENSG00000263620 |
| LAPTM5 | LSM2 | | EIF4A1 | RBM42 | AKR1C2 | AFDN |
| FGL2 | TMEM219 | | SLTM | LRATD2 | FNBP1L | MYCBP2 |
| PROX1 | ING1 | | NDUFAF4 | PLAUR | PRKCH | OGA |
| SGCE | TACR1 | | KLHL21 | CCDC9 | IL15RA | AKAP9 |
| RCSD1 | SNAPIN | | NOP58 | MYL12A | SLC40A1 | C1orf115 |
| HES4 | BTNL9 | | RPS10-NUDT3 | RPS12 | PTGIS | HLA-F |
| VPS35L | ADGRF5 | | ENSG00000270571 | FUNDC2 | SELENOK | FABP4 |
| AP3S1 | BUD31 | | GLA | IK | ANXA3 | EGLN1 |
| CHURC1 | TRIM16 | | GNA13 | BORCS7 | APOL3 | SLC2A4RG |
| H1-0 | | | DKC1 | RRAGC | | MTIF3 |
| LRPAP1 | | | SLC43A3 | LINC00963 | | NDUFB5 |
| TFF3 | | | MAP2K3 | ANAPC16 | | TPD52L1 |
| MBNL2 | | | CCDC86 | RTRAF | | COX7A1 |
| ST13 | | | DDX46 | FKBPL | | SNTG2 |
| S1PR1 | | | AKIRIN2 | CFAP36 | | CMPK1 |
| DSTN | | | CNP | SDHD | | NME3 |
| TECR | | | CDK12 | PPM1K | | MT-ND3 |
| EPHX1 | | | RAB21 | RPL3 | | LSM8 |
| FAM3C | | | WDR43 | HADHA | | |
| DKK3 | | | DTX3L | CFDP1 | | |
| SMIM19 | | | EDNRB | MYNN | | |
| SRSF2 | | | MPLKIP | RAD21 | | |
| OCIAD1 | | | TRIP10 | IGBP1 | | |
| PCBP1 | | | ANKRD17 | HAX1 | | |
| CDH13 | | | SRSF1 | NR1D1 | | |
| BAG3 | | | GCA | KPNA2 | | |
| RNF130 | | | BDKRB2 | RAB24 | | |
| TBX1 | | | DCAF13 | COX7A2L | | |
| CAPN2 | | | ARID5A | INSIG1 | | |
| MMRN1 | | | PPTC7 | CXCL10 | | |
| SKP1 | | | MAFG | RPL7A | | |
| PEPD | | | CWC25 | SEC11A | | |
| | | | SCML1 | MCUB | | |
| | | | MIR222HG | ARMCX1 | | |
| | | | ZEB1 | ENSG00000277969 | | |
| | | | SF3A3 | RABGGTB | | |
| | | | STAG2 | SRSF11 | | |
| | | | ZPR1 | AHR | | |
| | | | COPS3 | CCNH | | |
| | | | RBM22 | NFATC3 | | |
| | | | UTP4 | PLEKHG1 | | |
| | | | ZNF800 | RAB29 | | |
| | | | ENSG00000278607 | PPP2R2A | | |
| | | | TXNRD1 | HIKESHI | | |
| | | | RB1CC1 | BTF3 | | |
| | | | PHLDA1-AS1 | UBE2D2 | | |
| | | | TFB2M | | | |
| | | | TNFRSF12A | | | |
| | | | ETF1 | | | |
| | | | MED19 | | | |
| | | | ABT1 | | | |
| | | | RYBP | | | |

| | | | |
|---|---|---|---|
| | | | CDK17 |
| | | | CAMSAP2 |
| | | | RAPGEF4 |
| | | | GNL3 |
| | | | TASOR2 |
| | | | TNKS2 |
| | | | TFAM |
| | | | LUCAT1 |
| | | | UBE2W |
| | | | HTRA2 |
| | | | CRTC2 |
| | | | SDF2 |
| | | | IST1 |
| | | | TP53BP2 |
| | | | POLR1C |
| | | | RASSF1 |
| | | | FBXO34 |
| | | | TNFRSF10A |
| | | | SUPT4H1 |
| | | | BCL9L |
| | | | WDR74 |
| | | | CHMP4B |
| | | | HMGA1 |
| | | | SASH1 |
| | | | PNO1 |
| | | | NGRN |
| | | | TRIR |
| | | | ERF |
| | | | GSK3A |
| | | | GNAI3 |
| | | | NTMT1 |
| | | | ADAMTS4 |
| | | | ATP6V1C1 |
| | | | PRPF4B |
| | | | PSMG1 |
| | | | GTF2B |
| | | | FAM91A1 |
| | | | FUBP1 |
| | | | RNF114 |
| | | | SLC30A7 |
| | | | PPP2CA |
| | | | RNF149 |
| | | | ATF4 |

| **Cluster shared DEgenes of endothelial cells** | |
|---|---|
| Up in BP lesional skin | Down in BP lesional skin |
| S100A16 | RPL30 |
| TPM4 | RPS27 |
| KRT6C | CLU |
| RPL13A | PNRC1 |
| FKBP1A | CD9 |
| CALR | N4BP2L2 |
| AP2S1 | TXNIP |
| GADD45GIP1 | SNHG1 |
| S100A2 | GADD45B |
| KRT16 | DCD |
| KRT14 | RPL35A |

| | |
|---|---|
| NDUFB2 | ZFAS1 |
| S100A9 | HLA-E |
| COL4A2 | ARL6IP1 |
| PPIB | SCGB2A2 |
| ENG | RPL11 |
| IFITM2 | EEF1D |
| SH3BGRL3 | SNHG29 |
| PPIA | H3-3B |
| PFN1 | NACA |
| CCDC85B | PDK4 |
| NDUFS5 | FTL |
| KRT5 | CEBPD |
| HSBP1 | CDKN1A |
| CFL1 | MAFF |
| COL4A1 | KLF4 |
| PDIA3 | CCL2 |
| HSP90B1 | DNAJA1 |
| KRT17 | PCAT19 |
| ADAMTS9 | MAP1LC3B |
| KRT6A | JUN |
| MYH9 | |
| CAVIN3 | |
| LGALS7B | |
| ENO1 | |
| ADAM15 | |
| SELENOW | |
| RAB13 | |
| S100A8 | |
| TPM3 | |
| MIF | |
| HLA-A | |
| TPI1 | |
| PRELID1 | |
| SFN | |
| ACTN1 | |
| RPS26 | |
| HSPA5 | |
| SELENOH | |
| S100A11 | |

| Table S4 \| BP Blood pDC top10% DEgenes | | | | | |
|---|---|---|---|---|---|
| **Top10% Degenes of blood pDCs between healthy controls and BP patients in active stage** | | | | | |
| GeneName | HC | BP active | log2FC | p_value | p_val_adjust |
| CD14 | 0.027400864 | 1.563624995 | 5.83452937 | 2.17E-13 | 8.33E-09 |
| CD36 | 0.018937672 | 0.782113773 | 5.36804757 | 4.21E-12 | 1.61E-07 |
| HLA-DQA2 | 0.025814979 | 0.734318799 | 4.83012621 | 1.18E-15 | 4.51E-11 |
| BST1 | 0.018563621 | 0.499580599 | 4.75016738 | 1.25E-10 | 4.78E-06 |
| ASGR1 | 0.048124377 | 1.037229002 | 4.42982277 | 9.60E-15 | 3.68E-10 |
| SGK1 | 0.05788602 | 1.040488754 | 4.1679026 | 2.46E-12 | 9.44E-08 |
| PLOD3 | 0.01997555 | 0.333700432 | 4.06224643 | 1.15E-06 | 0.044035029 |
| GSTM1 | 0.023819505 | 0.392667473 | 4.04309274 | 2.01E-09 | 7.71E-05 |
| ENSG00000255639 | 0.020365106 | 0.313664744 | 3.94505218 | 8.34E-08 | 0.003197687 |
| ALDH2 | 0.069939535 | 1.072615911 | 3.93888155 | 2.54E-19 | 9.75E-15 |
| BHLHE40 | 0.045750677 | 0.693994956 | 3.92306018 | 1.57E-12 | 6.02E-08 |
| ENSG00000288819 | 0.03628763 | 0.544692421 | 3.90789204 | 4.78E-08 | 0.001831965 |
| GNG10 | 0.042383127 | 0.597891956 | 3.81832288 | 3.95E-10 | 1.51E-05 |
| S100A9 | 2.026414365 | 27.94426979 | 3.78555136 | 3.83E-36 | 1.47E-31 |
| GAS5 | 0.114663631 | 1.5168679 | 3.72561568 | 4.34E-28 | 1.67E-23 |
| FCGR1A | 0.037074936 | 0.457665865 | 3.62577859 | 3.04E-09 | 1.17E-04 |
| SNAPIN | 0.023819505 | 0.28770992 | 3.59439961 | 7.30E-07 | 0.02799038 |
| FUCA1 | 0.033651905 | 0.397643591 | 3.56271585 | 5.54E-10 | 2.13E-05 |
| HLA-DRB5 | 0.922491571 | 10.09555798 | 3.45204111 | 6.12E-42 | 2.35E-37 |
| CAPG | 0.118053691 | 1.245369551 | 3.39905886 | 1.39E-14 | 5.33E-10 |
| AGA | 0.02640027 | 0.263946539 | 3.32162114 | 1.30E-06 | 0.049885848 |
| S100A8 | 1.405709658 | 14.01790728 | 3.31790044 | 3.97E-24 | 1.52E-19 |
| ATP6V1C1 | 0.027759272 | 0.271774995 | 3.29137111 | 1.27E-06 | 0.048694377 |
| BCKDHA | 0.057676737 | 0.540092528 | 3.22714513 | 1.18E-11 | 4.53E-07 |
| ENSG00000267120 | 0.038539171 | 0.356318183 | 3.20876866 | 2.38E-07 | 0.009123209 |
| JUN | 0.522036586 | 4.781527265 | 3.19524868 | 2.41E-22 | 9.25E-18 |
| CD63 | 0.326414939 | 2.941504866 | 3.17177543 | 5.43E-23 | 2.08E-18 |
| DNTTIP1 | 0.056676144 | 0.500060837 | 3.14129011 | 1.71E-11 | 6.54E-07 |
| BASP1 | 0.036815598 | 0.301431002 | 3.03343876 | 8.15E-07 | 0.031264778 |
| TSC22D2 | 0.05878937 | 0.478565065 | 3.02508786 | 2.77E-10 | 1.06E-05 |
| PSMB5 | 0.038539171 | 0.299606409 | 2.95867104 | 7.56E-07 | 0.02897376 |
| SNHG3 | 0.051793053 | 0.386649075 | 2.90019424 | 1.76E-08 | 6.73E-04 |
| C3AR1 | 0.10864899 | 0.800225499 | 2.88073183 | 1.43E-16 | 5.48E-12 |
| RPIA | 0.061342556 | 0.450613838 | 2.87693142 | 7.17E-09 | 2.75E-04 |
| LINC02432 | 0.21645628 | 1.566348607 | 2.85525777 | 2.09E-20 | 8.02E-16 |
| RETN | 0.391766671 | 2.833496568 | 2.85451688 | 3.85E-27 | 1.48E-22 |
| ARID5A | 0.043918207 | 0.316570253 | 2.84963465 | 1.39E-07 | 0.005327327 |
| NFIL3 | 0.057608996 | 0.412842333 | 2.84122489 | 1.58E-10 | 6.07E-06 |
| MARCO | 0.144040997 | 1.021523162 | 2.82617052 | 9.02E-11 | 3.46E-06 |
| TAF3 | 0.0399511 | 0.277581533 | 2.79660447 | 8.45E-08 | 0.003240577 |
| TCN2 | 0.053683781 | 0.357969257 | 2.7372775 | 1.64E-08 | 6.29E-04 |
| ADSL | 0.066703642 | 0.443154658 | 2.73197284 | 2.22E-09 | 8.53E-05 |
| HLA-DQA1 | 1.049894929 | 6.877468926 | 2.71163276 | 1.81E-26 | 6.92E-22 |
| HAUS4 | 0.048678856 | 0.298393548 | 2.61584918 | 2.97E-09 | 1.14E-04 |
| TNFAIP8L2 | 0.095128904 | 0.56743233 | 2.57649269 | 1.99E-09 | 7.65E-05 |
| TMEM176A | 0.347816383 | 2.052925308 | 2.56128334 | 5.52E-22 | 2.12E-17 |
| ENSG00000289341 | 0.219047329 | 1.226638837 | 2.48539601 | 1.46E-14 | 5.59E-10 |
| TAX1BP3 | 0.093723794 | 0.518674396 | 2.46834189 | 7.49E-13 | 2.87E-08 |
| ELOA | 0.058261402 | 0.316947939 | 2.44363355 | 4.38E-07 | 0.016798724 |

| | | | | | |
|---|---|---|---|---|---|
| ENSG00000261222 | 0.136659584 | 0.728410286 | 2.41416466 | 8.25E-13 | 3.16E-08 |
| ATP6V1A | 0.05829816 | 0.309902444 | 2.41029188 | 3.38E-10 | 1.30E-05 |
| EIF4EBP3 | 0.068131075 | 0.36032174 | 2.40290083 | 2.47E-08 | 9.48E-04 |
| SPINT2 | 0.090173012 | 0.47516083 | 2.39764829 | 5.83E-08 | 0.002237598 |
| FOSB | 0.337534216 | 1.772837479 | 2.39295462 | 1.36E-18 | 5.21E-14 |
| IER5L | 0.086791004 | 0.453156783 | 2.38439286 | 1.18E-08 | 4.54E-04 |
| TMEM35B | 0.081469276 | 0.416692342 | 2.35465459 | 2.03E-09 | 7.79E-05 |
| HSPA1A | 0.089719276 | 0.458855 | 2.35454845 | 1.54E-09 | 5.92E-05 |
| COMMD10 | 0.077599449 | 0.39602739 | 2.3514819 | 1.65E-07 | 0.00632478 |
| FAM20A | 0.071908856 | 0.364967483 | 2.34352656 | 5.35E-07 | 0.020505812 |
| MS4A6A | 0.380250052 | 1.927393948 | 2.34163113 | 1.80E-14 | 6.92E-10 |
| SDHAF2 | 0.048678856 | 0.245493084 | 2.33431522 | 1.47E-07 | 0.005638951 |
| GOLGA2 | 0.062300638 | 0.312821334 | 2.32802007 | 1.94E-07 | 0.007449676 |
| MTX1 | 0.07622052 | 0.373842421 | 2.29417893 | 5.50E-08 | 0.00210774 |
| FBP1 | 0.355060444 | 1.730790581 | 2.28529463 | 1.41E-16 | 5.41E-12 |
| NOP56 | 0.062300638 | 0.294326494 | 2.24009857 | 8.25E-07 | 0.031637896 |
| ID2 | 0.693916266 | 3.276063186 | 2.23912969 | 1.28E-26 | 4.89E-22 |
| IL18 | 0.064046902 | 0.300150381 | 2.22848481 | 4.38E-07 | 0.016814198 |
| JAGN1 | 0.087777817 | 0.409454352 | 2.22177433 | 4.87E-12 | 1.87E-07 |
| EIF3E | 0.567741842 | 2.645526183 | 2.22024772 | 7.99E-26 | 3.06E-21 |
| IMPA2 | 0.135728417 | 0.625931162 | 2.2052812 | 1.43E-10 | 5.48E-06 |
| SCP2 | 0.236165704 | 1.084880994 | 2.19966542 | 1.48E-18 | 5.68E-14 |
| RPS20 | 1.174703858 | 5.361361926 | 2.19030243 | 2.43E-31 | 9.32E-27 |
| ETHE1 | 0.092119423 | 0.419963366 | 2.1886862 | 1.27E-10 | 4.88E-06 |
| PNP | 0.090732194 | 0.409560421 | 2.17438985 | 5.28E-08 | 0.002026622 |
| TNFSF13 | 0.382136567 | 1.720053539 | 2.17029325 | 2.71E-23 | 1.04E-18 |
| LINC00963 | 0.060246771 | 0.271046942 | 2.16958691 | 1.15E-08 | 4.41E-04 |
| CYB561A3 | 0.071736269 | 0.322591418 | 2.16893344 | 5.12E-08 | 0.001961587 |
| HERC1 | 1.166294159 | 0.261217423 | -2.1586087 | 4.62E-14 | 1.77E-09 |
| GXYLT1 | 0.572412998 | 0.127979791 | -2.1611404 | 2.13E-10 | 8.16E-06 |
| HES4 | 5.721364315 | 1.26514392 | -2.1770577 | 2.43E-20 | 9.32E-16 |
| COMMD3 | 0.72227909 | 0.15905942 | -2.1829906 | 6.60E-10 | 2.53E-05 |
| FRS2 | 0.737878086 | 0.161569579 | -2.1912269 | 1.27E-06 | 0.048775808 |
| SLC9A8 | 0.84694665 | 0.183674104 | -2.2051229 | 2.26E-08 | 8.65E-04 |
| FAM120B | 0.490899256 | 0.106189566 | -2.208785 | 1.59E-07 | 0.006116033 |
| PPP6R2 | 1.112688048 | 0.236973283 | -2.2312529 | 1.75E-10 | 6.70E-06 |
| MBD3 | 0.614885785 | 0.129953823 | -2.2423194 | 1.30E-07 | 0.004972834 |
| INPPL1 | 0.474701815 | 0.100289591 | -2.2428497 | 6.83E-07 | 0.026182004 |
| KHSRP | 0.633495694 | 0.132863807 | -2.2533867 | 8.35E-08 | 0.003202656 |
| CKB | 3.763631685 | 0.787629937 | -2.2565356 | 1.34E-12 | 5.12E-08 |
| MYO15B | 1.683886839 | 0.35050944 | -2.26427 | 7.78E-13 | 2.98E-08 |
| NBPF10 | 0.950729879 | 0.197659071 | -2.2660213 | 2.16E-10 | 8.28E-06 |
| SUGP2 | 0.938035525 | 0.194725107 | -2.2682036 | 5.81E-07 | 0.022267345 |
| PLAA | 0.828575374 | 0.171543761 | -2.2720563 | 3.47E-09 | 1.33E-04 |
| ATXN2 | 0.666492798 | 0.137288624 | -2.2793772 | 5.72E-10 | 2.19E-05 |
| SFMBT2 | 1.243893898 | 0.254590174 | -2.2886148 | 7.99E-14 | 3.06E-09 |
| TNRC18 | 0.850444902 | 0.170105521 | -2.3217878 | 3.19E-07 | 0.012237851 |
| FAM168B | 0.962266661 | 0.191048827 | -2.3324953 | 1.99E-09 | 7.63E-05 |
| ZNF333 | 0.655103757 | 0.129212163 | -2.3419815 | 1.59E-08 | 6.10E-04 |
| SCAF4 | 1.963891101 | 0.386997376 | -2.3433192 | 1.69E-11 | 6.47E-07 |
| CDK16 | 0.753162347 | 0.147918683 | -2.3481566 | 1.50E-07 | 0.005747498 |
| GARS1-DT | 0.899026922 | 0.174379477 | -2.3661341 | 6.07E-12 | 2.33E-07 |
| FBRSL1 | 1.247396226 | 0.238785204 | -2.3851344 | 6.54E-09 | 2.51E-04 |
| SPRED1 | 0.66135338 | 0.126387927 | -2.3875627 | 2.27E-08 | 8.69E-04 |
| MHENCR | 0.963590278 | 0.177119686 | -2.4436953 | 1.51E-07 | 0.005787309 |
| SMPD4 | 0.631586359 | 0.1158921 | -2.4461978 | 8.48E-07 | 0.03250525 |
| MTMR3 | 1.586256487 | 0.285842544 | -2.4723335 | 2.00E-12 | 7.67E-08 |

| | | | | | |
|---|---|---|---|---|---|
| REV3L | 0.63554434 | 0.113542959 | -2.4847545 | 4.02E-07 | 0.015406049 |
| AP1G1 | 0.720795148 | 0.127629214 | -2.4976307 | 5.93E-07 | 0.022722913 |
| USP38 | 0.593684465 | 0.104393781 | -2.5076606 | 8.40E-07 | 0.032219121 |
| KCTD15 | 0.673333674 | 0.117083287 | -2.5237865 | 2.64E-07 | 0.010121825 |
| UICLM | 4.19120426 | 0.725925886 | -2.5294707 | 7.75E-23 | 2.97E-18 |
| LMBR1 | 0.832666026 | 0.14247971 | -2.5469815 | 1.11E-11 | 4.24E-07 |
| PREP | 0.933668588 | 0.158550908 | -2.5579644 | 2.33E-10 | 8.95E-06 |
| RYK | 2.04312932 | 0.332084917 | -2.6211564 | 3.41E-18 | 1.31E-13 |
| ZNF641 | 0.605341969 | 0.095019923 | -2.6714484 | 3.13E-07 | 0.01201581 |
| PHACTR4 | 1.209744899 | 0.176972651 | -2.7731045 | 2.17E-09 | 8.33E-05 |
| RPS6KA5 | 0.680730672 | 0.098024183 | -2.7958745 | 6.47E-07 | 0.024809799 |
| LPAR2 | 0.829222022 | 0.109157297 | -2.9253499 | 1.35E-07 | 0.005165303 |
| Note: HC, healthy control; BP active, BP patients in active stage. | | | | | |

| **Top10% Degenes of blood pDCs between healthy controls and BP patients in remission stage** | | | | | |
|---|---|---|---|---|---|
| GeneName | HC | BP remission | log2FC | p_value | p_val_adjust |
| CD14 | 0.027400864 | 3.299538234 | 6.91189896 | 6.45E-28 | 2.47E-23 |
| CD36 | 0.018937672 | 2.174008449 | 6.84295472 | 3.37E-29 | 1.29E-24 |
| FCGR1A | 0.037074936 | 1.865436828 | 5.6529255 | 6.09E-25 | 2.33E-20 |
| BST1 | 0.018563621 | 0.781706397 | 5.39607677 | 4.04E-15 | 1.55E-10 |
| ALDH2 | 0.069939535 | 2.457965627 | 5.13521273 | 1.65E-32 | 6.33E-28 |
| S100A9 | 2.026414365 | 66.67385332 | 5.04011999 | 9.25E-43 | 3.55E-38 |
| S100A8 | 1.405709658 | 44.1944215 | 4.97449373 | 2.09E-32 | 8.03E-28 |
| ASGR1 | 0.048124377 | 1.380795404 | 4.84258789 | 2.82E-21 | 1.08E-16 |
| HLA-DQA2 | 0.025814979 | 0.695328854 | 4.75141513 | 1.11E-13 | 4.25E-09 |
| CAPG | 0.118053691 | 2.514182447 | 4.4125743 | 4.74E-24 | 1.82E-19 |
| GSTM1 | 0.023819505 | 0.484924843 | 4.34754581 | 2.89E-11 | 1.11E-06 |
| SERPING1 | 0.035940712 | 0.696354494 | 4.27613103 | 9.27E-13 | 3.55E-08 |
| ATP6V1C1 | 0.027759272 | 0.533722674 | 4.26504869 | 1.46E-13 | 5.59E-09 |
| GNG10 | 0.042383127 | 0.803971107 | 4.24558173 | 2.83E-16 | 1.09E-11 |
| SNAPIN | 0.023819505 | 0.440977415 | 4.21048941 | 6.94E-13 | 2.66E-08 |
| NCF1 | 0.749338377 | 13.49958629 | 4.17115405 | 1.32E-26 | 5.08E-22 |
| CD63 | 0.326414939 | 5.634497734 | 4.10950802 | 3.71E-31 | 1.42E-26 |
| JUN | 0.522036586 | 9.00468223 | 4.10845254 | 1.02E-29 | 3.90E-25 |
| PLOD3 | 0.01997555 | 0.336383239 | 4.07379869 | 9.50E-07 | 0.036444583 |
| ENSG00000255639 | 0.020365106 | 0.331213418 | 4.02358992 | 1.12E-09 | 4.30E-05 |
| TIMM9 | 0.018563621 | 0.301820364 | 4.02314008 | 1.10E-07 | 0.004216311 |
| HSPA1A | 0.089719276 | 1.440090848 | 4.00459804 | 1.42E-21 | 5.44E-17 |
| BHLHE40 | 0.045750677 | 0.715058548 | 3.96619637 | 8.14E-14 | 3.12E-09 |
| ENSG00000288819 | 0.03628763 | 0.544437694 | 3.9072172 | 5.02E-09 | 1.92E-04 |
| FCGR1B | 0.045997681 | 0.683657114 | 3.8936399 | 1.30E-10 | 4.98E-06 |
| ABCE1 | 0.027759272 | 0.403662651 | 3.86210849 | 1.17E-11 | 4.51E-07 |
| CIAPIN1 | 0.018563621 | 0.269503017 | 3.85975135 | 3.91E-07 | 0.015013248 |
| MRPL17 | 0.018937672 | 0.267098597 | 3.81804147 | 3.86E-07 | 0.014812356 |
| DBNDD2 | 0.02914908 | 0.397878367 | 3.7708052 | 1.47E-09 | 5.64E-05 |
| HLA-DRB5 | 0.922491571 | 12.27848809 | 3.73445339 | 1.10E-42 | 4.23E-38 |
| GM2A | 0.040250519 | 0.532216353 | 3.72493354 | 8.67E-12 | 3.33E-07 |
| MS4A6A | 0.380250052 | 5.016828233 | 3.72175519 | 2.37E-26 | 9.10E-22 |
| ENSG00000267120 | 0.038539171 | 0.502510174 | 3.70475536 | 4.84E-11 | 1.86E-06 |
| ATIC | 0.023819505 | 0.310057909 | 3.70232234 | 1.60E-08 | 6.13E-04 |
| SETD5 | 0.018563621 | 0.230420024 | 3.63371602 | 2.89E-07 | 0.011092417 |
| ENSG00000257764 | 0.10430594 | 1.199872737 | 3.52398817 | 1.58E-13 | 6.07E-09 |
| RETN | 0.391766671 | 4.40155852 | 3.48994787 | 6.95E-28 | 2.66E-23 |

| | | | | | |
|---|---|---|---|---|---|
| TLR8 | 0.025978075 | 0.290976415 | 3.48553581 | 4.61E-08 | 0.00176798 |
| SGK1 | 0.05788602 | 0.647857802 | 3.48439032 | 1.75E-07 | 0.006720169 |
| RPE | 0.01997555 | 0.214093304 | 3.42193255 | 4.25E-07 | 0.01628731 |
| SMIM15 | 0.0443786 | 0.467032839 | 3.39558793 | 1.04E-11 | 3.98E-07 |
| PSMB5 | 0.038539171 | 0.404963042 | 3.3933928 | 3.94E-08 | 0.001509441 |
| IER5L | 0.086791004 | 0.896590607 | 3.36883196 | 4.84E-16 | 1.86E-11 |
| BASP1 | 0.036815598 | 0.360211144 | 3.29045377 | 8.26E-08 | 0.003167222 |
| FPR2 | 0.039751948 | 0.381896448 | 3.26408404 | 1.52E-08 | 5.83E-04 |
| DNTTIP1 | 0.056676144 | 0.527205046 | 3.21755066 | 2.78E-09 | 1.06E-04 |
| DYNLT3 | 0.029856807 | 0.275827157 | 3.20763273 | 2.40E-08 | 9.22E-04 |
| GAS5 | 0.114663631 | 1.042957577 | 3.1852007 | 5.76E-17 | 2.21E-12 |
| MARCO | 0.144040997 | 1.293157128 | 3.16634619 | 2.92E-16 | 1.12E-11 |
| FUCA1 | 0.033651905 | 0.300487989 | 3.15854725 | 1.03E-08 | 3.96E-04 |
| BCKDHA | 0.057676737 | 0.51188331 | 3.1497535 | 7.58E-14 | 2.90E-09 |
| LYZ | 4.250869887 | 37.48040322 | 3.14030647 | 1.01E-22 | 3.87E-18 |
| GTF2E2 | 0.03148575 | 0.275675128 | 3.1301982 | 6.89E-09 | 2.64E-04 |
| ALAS1 | 0.027400864 | 0.238525593 | 3.1218508 | 1.21E-06 | 0.046561097 |
| ATP6V1A | 0.05829816 | 0.505950547 | 3.11747412 | 1.21E-12 | 4.63E-08 |
| HLA-DMB | 0.2377372 | 2.059411523 | 3.11479258 | 4.07E-24 | 1.56E-19 |
| DIS3 | 0.027759272 | 0.239156922 | 3.10691593 | 1.28E-06 | 0.049074286 |
| VCAN | 0.279933753 | 2.371786724 | 3.08281693 | 6.75E-14 | 2.59E-09 |
| PRKD3 | 0.03628763 | 0.305323345 | 3.07278815 | 5.45E-07 | 0.020904617 |
| HLA-DQA1 | 1.049894929 | 8.822942183 | 3.07101488 | 2.02E-26 | 7.73E-22 |
| FOSB | 0.337534216 | 2.701158613 | 3.0004727 | 2.56E-22 | 9.83E-18 |
| IL18 | 0.064046902 | 0.511053431 | 2.99627344 | 4.03E-07 | 0.015468287 |
| TCN2 | 0.053683781 | 0.425357055 | 2.98611619 | 4.08E-10 | 1.57E-05 |
| ENSG00000289341 | 0.219047329 | 1.729018792 | 2.98063902 | 4.69E-19 | 1.80E-14 |
| TAF3 | 0.0399511 | 0.308422045 | 2.94859876 | 1.39E-08 | 5.34E-04 |
| TNFAIP8L2 | 0.095128904 | 0.733482755 | 2.94680739 | 1.28E-15 | 4.92E-11 |
| EXOSC5 | 0.06735081 | 0.506792526 | 2.91162806 | 2.08E-14 | 7.99E-10 |
| C14orf119 | 0.060635103 | 0.449741071 | 2.89086948 | 2.47E-11 | 9.47E-07 |
| CIDEB | 0.035862346 | 0.261926696 | 2.86862134 | 2.42E-07 | 0.00929781 |
| SELL | 0.097160616 | 0.708750666 | 2.86683464 | 3.51E-12 | 1.35E-07 |
| MGST2 | 0.047951847 | 0.348259246 | 2.86050338 | 3.12E-07 | 0.011981791 |
| GCA | 0.213476353 | 1.541678225 | 2.85235351 | 1.80E-24 | 6.91E-20 |
| TMEM176A | 0.347816383 | 2.492743291 | 2.84133652 | 2.45E-17 | 9.40E-13 |
| TOR4A | 0.039751948 | 0.283153477 | 2.83248679 | 7.20E-07 | 0.027623418 |
| DCTN6 | 0.030275874 | 0.214049468 | 2.82170375 | 5.31E-07 | 0.020345976 |
| SAP30 | 0.091121155 | 0.634736402 | 2.80029965 | 8.43E-16 | 3.23E-11 |
| CYFIP1 | 0.107008483 | 0.739946249 | 2.78969531 | 3.74E-12 | 1.44E-07 |
| STING1 | 0.09385089 | 0.6289064 | 2.74440299 | 3.09E-16 | 1.18E-11 |
| C3AR1 | 0.10864899 | 0.717912812 | 2.72413388 | 1.37E-15 | 5.27E-11 |
| FH | 0.0443786 | 0.292197711 | 2.71900881 | 1.01E-06 | 0.03865224 |
| SQOR | 0.293717354 | 1.924386516 | 2.71189818 | 2.62E-26 | 1.00E-21 |
| ETHE1 | 0.092119423 | 0.595909888 | 2.6935169 | 1.78E-11 | 6.84E-07 |
| ACADM | 0.051793053 | 0.327794615 | 2.66196164 | 3.68E-07 | 0.014100557 |
| LINC00963 | 0.060246771 | 0.380303289 | 2.6581946 | 5.11E-11 | 1.96E-06 |
| TNFSF12 | 0.088626114 | 0.55871639 | 2.65631238 | 1.06E-11 | 4.06E-07 |
| CRLS1 | 0.039751948 | 0.249841802 | 2.65191742 | 5.02E-08 | 0.001926487 |
| GIMAP4 | 0.440516565 | 2.760625979 | 2.64772726 | 6.64E-24 | 2.55E-19 |
| EVI2A | 0.067426462 | 0.422256377 | 2.64673241 | 6.75E-09 | 2.59E-04 |
| LIPA | 0.287647519 | 1.797563955 | 2.64366917 | 4.09E-20 | 1.57E-15 |
| WSB2 | 0.051461299 | 0.321255822 | 2.64216281 | 3.55E-07 | 0.013629022 |
| MNDA | 1.28033672 | 7.91885828 | 2.62876916 | 8.58E-24 | 3.29E-19 |
| ITGAM | 0.086509543 | 0.535039756 | 2.62871491 | 9.19E-11 | 3.52E-06 |
| PLBD1 | 0.285509548 | 1.751992058 | 2.61738534 | 1.19E-22 | 4.57E-18 |
| MIF4GD | 0.074048887 | 0.454107436 | 2.61648371 | 3.88E-09 | 1.49E-04 |

| | | | | | |
|---|---|---|---|---|---|
| SLC35A4 | 0.065009361 | 0.39457952 | 2.60159669 | 2.51E-12 | 9.64E-08 |
| GCDH | 0.037074936 | 0.223864387 | 2.59410893 | 8.06E-07 | 0.030909535 |
| IL27RA | 0.099138604 | 0.594203654 | 2.58343862 | 5.71E-12 | 2.19E-07 |
| KRCC1 | 0.052215526 | 0.312951352 | 2.58338766 | 3.36E-07 | 0.0128996 |
| PNP | 0.090732194 | 0.543159253 | 2.5816888 | 4.09E-12 | 1.57E-07 |
| NOLC1 | 0.063690324 | 0.379658931 | 2.57555784 | 1.12E-12 | 4.31E-08 |
| GPN3 | 0.127965887 | 0.756165866 | 2.56294346 | 5.53E-16 | 2.12E-11 |
| NOP56 | 0.062300638 | 0.363775878 | 2.54573104 | 5.12E-10 | 1.96E-05 |
| GSN | 0.093217684 | 0.54372417 | 2.54419938 | 1.36E-11 | 5.22E-07 |
| NFIL3 | 0.057608996 | 0.331952097 | 2.52660904 | 5.62E-11 | 2.16E-06 |
| MSRB1 | 0.18494218 | 1.06561837 | 2.52654466 | 1.26E-16 | 4.84E-12 |
| PLIN3 | 0.166612252 | 0.955886825 | 2.52034532 | 1.58E-16 | 6.07E-12 |
| FOS | 4.827537243 | 27.596939 | 2.51514896 | 3.21E-27 | 1.23E-22 |
| PTCH2 | 0.09005991 | 0.506103973 | 2.49047687 | 2.67E-09 | 1.03E-04 |
| COMMD10 | 0.077599449 | 0.435038325 | 2.48702419 | 1.10E-11 | 4.22E-07 |
| HSD17B4 | 0.082257821 | 0.457892012 | 2.47678263 | 8.25E-12 | 3.16E-07 |
| AHSA1 | 0.097181049 | 0.538707924 | 2.47075638 | 9.82E-13 | 3.77E-08 |
| IFIT2 | 0.128877932 | 0.71330597 | 2.4685158 | 3.99E-09 | 1.53E-04 |
| HAUS4 | 0.048678856 | 0.268480545 | 2.46345039 | 6.47E-10 | 2.48E-05 |
| ADSL | 0.066703642 | 0.3660636 | 2.45625689 | 2.05E-07 | 0.007850265 |
| SLAMF7 | 0.081084645 | 0.444238281 | 2.45383308 | 6.32E-08 | 0.002423814 |
| GK | 0.065921316 | 0.359548598 | 2.44736983 | 2.47E-08 | 9.48E-04 |
| TXNDC15 | 0.04912463 | 0.267079698 | 2.44275187 | 1.03E-06 | 0.039562015 |
| HSPA1B | 0.060551748 | 0.328523468 | 2.43975593 | 1.13E-07 | 0.004348268 |
| PLAC8 | 1.100957196 | 5.968028661 | 2.43849608 | 1.65E-19 | 6.34E-15 |
| SCAF4 | 1.963891101 | 0.36244779 | -2.4378698 | 5.54E-11 | 2.12E-06 |
| MPPE1 | 1.048531957 | 0.191633438 | -2.4519496 | 1.07E-08 | 4.09E-04 |
| GGA1 | 1.041230783 | 0.19012211 | -2.4532916 | 2.22E-08 | 8.51E-04 |
| KIF22 | 1.258513039 | 0.229652217 | -2.4541976 | 8.30E-12 | 3.18E-07 |
| NSMF | 1.200633841 | 0.216708815 | -2.4699665 | 4.47E-09 | 1.71E-04 |
| KLF7 | 0.798902352 | 0.142755752 | -2.4844703 | 5.51E-09 | 2.11E-04 |
| TPPP3 | 1.333058559 | 0.237135313 | -2.4909577 | 1.65E-08 | 6.33E-04 |
| L3MBTL3 | 1.020861647 | 0.180467065 | -2.4999799 | 2.82E-08 | 0.001079927 |
| TAMALIN | 1.075054561 | 0.18989546 | -2.5011326 | 6.55E-08 | 0.002511947 |
| HECTD1 | 1.972951766 | 0.346867442 | -2.5078994 | 1.02E-14 | 3.93E-10 |
| NUDT4 | 1.045834443 | 0.181521283 | -2.5264439 | 1.50E-08 | 5.76E-04 |
| WASH6P | 0.845696187 | 0.145966492 | -2.5345022 | 2.16E-10 | 8.27E-06 |
| ENSG00000 272449 | 1.469146536 | 0.251395903 | -2.5469453 | 4.44E-13 | 1.70E-08 |
| UBXN6 | 0.5221263 | 0.089179579 | -2.5496135 | 1.11E-06 | 0.042517055 |
| TBC1D8 | 2.205819845 | 0.375820133 | -2.5532007 | 6.01E-13 | 2.30E-08 |
| ZNF292 | 1.130075898 | 0.192533944 | -2.553235 | 9.25E-09 | 3.55E-04 |
| HIPK2 | 1.569998138 | 0.265448394 | -2.5642595 | 9.35E-14 | 3.59E-09 |
| COMMD3 | 0.72227909 | 0.118434025 | -2.6084728 | 3.52E-07 | 0.013508678 |
| CACUL1 | 2.885309293 | 0.466862387 | -2.6276567 | 5.94E-17 | 2.28E-12 |
| ACVR1B | 0.857462414 | 0.13717605 | -2.6440448 | 3.61E-08 | 0.001383269 |
| LMBR1 | 0.832666026 | 0.132298894 | -2.653937 | 4.69E-08 | 0.00179917 |
| MYO15B | 1.683886839 | 0.266407825 | -2.6600868 | 1.42E-11 | 5.46E-07 |
| REV3L | 0.63554434 | 0.099313572 | -2.67793 | 3.26E-07 | 0.012505119 |
| FBRSL1 | 1.247396226 | 0.190906806 | -2.7079794 | 2.42E-09 | 9.29E-05 |
| ZDHHC1 | 1.48717053 | 0.227335215 | -2.709677 | 6.21E-10 | 2.38E-05 |
| KCTD15 | 0.673333674 | 0.100504465 | -2.744062 | 3.50E-07 | 0.013430827 |
| SOX4 | 1.238710903 | 0.182512342 | -2.7627736 | 6.22E-09 | 2.38E-04 |
| RNF145 | 1.94269342 | 0.281483146 | -2.7869378 | 1.74E-15 | 6.66E-11 |
| FRS2 | 0.737878086 | 0.104852665 | -2.8150189 | 3.02E-07 | 0.011572025 |
| SNX9 | 2.797915999 | 0.382245328 | -2.8717819 | 1.91E-14 | 7.31E-10 |
| MTMR3 | 1.586256487 | 0.212471959 | -2.9002817 | 1.27E-13 | 4.88E-09 |
| ADGRE1 | 1.363312792 | 0.16487179 | -3.0477001 | 6.06E-10 | 2.32E-05 |

| | | | | | |
|---|---|---|---|---|---|
| ADK | 2.39730671 | 0.28373679 | -3.0787894 | 1.88E-12 | 7.20E-08 |
| CRIP1 | 27.7137351 | 3.146343173 | -3.1388532 | 8.79E-31 | 3.37E-26 |
| RYK | 2.04312932 | 0.228317537 | -3.1616669 | 1.65E-12 | 6.32E-08 |
| PRR12 | 0.860844161 | 0.089998158 | -3.2577847 | 8.32E-07 | 0.031889143 |
| ICAM4 | 1.332191769 | 0.135650894 | -3.2958313 | 5.26E-09 | 2.02E-04 |
| HES4 | 5.721364315 | 0.513515156 | -3.4778804 | 7.82E-19 | 3.00E-14 |
| LIMD1 | 0.853446835 | 0.075614146 | -3.4965732 | 1.02E-06 | 0.039124327 |
| CKB | 3.763631685 | 0.209253615 | -4.168801 | 9.14E-14 | 3.51E-09 |
| SFMBT2 | 1.243893898 | 0.059323161 | -4.3901241 | 2.59E-07 | 0.009928096 |
| UICLM | 4.19120426 | 0.168141253 | -4.6396192 | 2.12E-19 | 8.14E-15 |
| Note: HC, healthy control; BP remission, BP patients in remission stage. | | | | | |

| **Top10% Degenes of blood pDCs between BP patients in active and remission stages** | | | | | |
|---|---|---|---|---|---|
| GeneName | BP active | BP remission | log2FC | p_value | p_val_adjust |
| ANKRD22 | 0.023846585 | 0.535921727 | 4.49016774 | 4.39E-12 | 1.68E-07 |
| CD163 | 0.038351915 | 0.457666307 | 3.57692558 | 5.51E-08 | 0.002112527 |
| SERPING1 | 0.066980582 | 0.696354494 | 3.37800711 | 1.65E-13 | 6.32E-09 |
| MCEMP1 | 0.04252406 | 0.381797515 | 3.16645646 | 1.01E-07 | 0.003870088 |
| ALDH1A1 | 0.105674695 | 0.900365023 | 3.09088006 | 3.01E-14 | 1.16E-09 |
| NCF1 | 1.980556343 | 13.49958629 | 2.76893755 | 2.97E-27 | 1.14E-22 |
| LAP3 | 0.827091503 | 4.542872014 | 2.45748581 | 2.67E-26 | 1.02E-21 |
| Note: BP active, BP patients in active stage; BP remission, BP patients in remission stage. | | | | | |

| Table S5 \| Gene set reference | | | | |
|---|---|---|---|---|
| Inflammatory response genes | Proliferation genes | TCR signaling | Cytotoxicity | IFN response |
| ABCF1 | ABL1 | CALM1 | GZMA | STAT1 |
| ADGRE5 | AGER | CALM2 | GZMB | STAT3 |
| ADORA1 | AIF1 | CALM3 | GZMH | MX1 |
| ADORA2A | ANXA1 | CD4 | GZMK | IRF1 |
| ADORA3 | ARG1 | CAST | GZMM | ISG15 |
| AFAP1L2 | ARG2 | CD247 | GNLY | ISG20 |
| AGER | ARMC5 | CD3D | PRF1 | IFITM1 |
| AHSG | BAX | CD3E | IFNG | IFITM2 |
| AIF1 | BCL6 | CD3G | TNF | IFITM3 |
| AIMP1 | BID | CSK | SERPINB9 | OAS1 |
| ALOX15 | BMI1 | DOK2 | CTSA | OAS2 |
| ALOX5AP | BMP4 | FYN | CTSB | OASL |
| ANXA1 | BTN2A2 | LCK | CTSC | SOCS1 |
| AOAH | BTN3A1 | NFATC1 | CTSD | SOCS3 |
| AOC3 | CADM1 | NFATC2 | CTSH | TRIM22 |
| AOX1 | CARD11 | PLEK | CTSW | APOL6 |
| APCS | CASP3 | PTPN11 | CST7 | IFNAR2 |
| APOL3 | CBLB | PTPN13 | CAPN2 | IFNGR1 |
| BLNK | CCDC88B | PTPN2 | PLEK | GBP1 |
| C2 | CCL19 | PTPN22 | | GBP2 |
| C3AR1 | CCL5 | PTPN4 | | GBP4 |
| C5 | CCND3 | PTPN6 | | GBP5 |
| CCL11 | CCR2 | PTPN7 | | BST2 |
| CCL13 | CD151 | PTPRC | | IFI16 |
| CCL20 | CD1D | PTPRCAP | | IFI35 |
| CCL21 | CD209 | S100A10 | | IFI44L |
| CCL22 | CD24 | S100A11 | | IFI6 |
| CCL23 | CD274 | S100A4 | | PARP8 |
| CCL24 | CD276 | S100A6 | | PARP9 |
| CCL26 | CD28 | ZAP70 | | |
| CCL3 | CD3E | DUSP1 | | |
| CCL3L3 | CD40LG | DUSP2 | | |
| CCL4 | CD46 | DUSP4 | | |
| CCL5 | CD55 | DUSP16 | | |
| CCR1 | CD6 | LAT | | |
| CCR2 | CD70 | FOS | | |
| CCR3 | CD80 | FOSB | | |
| CCR4 | CD81 | FOSL2 | | |
| CCR5 | CD86 | JUN | | |
| CCR7 | CEBPB | JUNB | | |
| CD40 | CLC | JUND | | |
| CD40LG | CLEC4G | NR4A1 | | |
| CDO1 | CLECL1P | NR4A2 | | |
| CEBPB | CORO1A | BATF | | |
| CFHR1 | CR1 | IRF1 | | |
| CHRNA7 | CRTAM | SH2D1A | | |
| CHST2 | CTLA4 | SH2D2A | | |
| CRP | CTNNB1 | MAP2K3 | | |
| CX3CL1 | CTPS1 | MAP3K4 | | |
| CXCL1 | DHPS | MAP3K8 | | |
| CXCL10 | DLG1 | MAP4K1 | | |
| CXCL11 | DLG5 | NFKB2 | | |
| CXCL2 | DNAJA3 | NFKBIA | | |
| CXCL6 | DOCK2 | NFKBIZ | | |
| CXCL8 | DOCK8 | REL | | |

| | | |
|---|---|---|
| CXCL9 | EBI3 | RELB |
| CXCR1 | EFNB1 | |
| CXCR2 | ELF4 | |
| CXCR4 | EPO | |
| CYBB | ERBB2 | |
| CYP4F11 | FADD | |
| ELF3 | FKBP1B | |
| F11R | FOXJ1 | |
| FOS | FOXP3 | |
| FPR2 | FYN | |
| GHRL | GLMN | |
| GHSR | GPAM | |
| GPR68 | GPNMB | |
| HDAC4 | HAVCR2 | |
| HDAC5 | HES1 | |
| HDAC7 | HHLA2 | |
| HDAC9 | HLA-A | |
| HRH1 | HLA-DMB | |
| IFNA2 | HLA-DPA1 | |
| IL10RB | HLA-DPB1 | |
| IL13 | HLA-DRB1 | |
| IL17C | HLA-E | |
| IL18RAP | HLA-G | |
| IL1A | HMGB1 | |
| IL1RAP | ICOSLG | |
| IL20 | IDO1 | |
| IL5 | IGF1 | |
| IL9 | IGF2 | |
| IRAK2 | IGFBP2 | |
| KLRG1 | IHH | |
| KNG1 | IL10 | |
| KRT1 | IL12B | |
| LBP | IL12RB1 | |
| LTB4R | IL15 | |
| LY75 | IL18 | |
| LYZ | IL1A | |
| MBL2 | IL1B | |
| MEFV | IL2 | |
| MGLL | IL20RB | |
| NFATC3 | IL21 | |
| NFATC4 | IL23A | |
| NFE2L1 | IL23R | |
| NFKB1 | IL27 | |
| NFRKB | IL2RA | |
| NFX1 | IL4 | |
| NLRP3 | IL4I1 | |
| NMI | IL6 | |
| NOD1 | IL6ST | |
| NOX4 | IRF1 | |
| ORM1 | ITCH | |
| ORM2 | JAK2 | |
| PARP4 | JAK3 | |
| PLA2G2D | KITLG | |
| PLA2G2E | LEP | |
| PLA2G7 | LGALS3 | |
| PRDX5 | LGALS9 | |
| PTAFR | LGALS9B | |
| PTX3 | LGALS9C | |

| | |
|---|---|
| RAC1 | LILRB1 |
| RELA | LILRB2 |
| RIPK2 | LILRB4 |
| S100A12 | LMBR1L |
| S100A8 | LMO1 |
| S100A9 | LRRC32 |
| S1PR3 | MAD1L1 |
| SCG2 | MALT1 |
| SELE | MAPK8IP1 |
| SIGIRR | MARCHF7 |
| TACR1 | MIR181C |
| TGFB1 | MIR21 |
| TNFAIP6 | MIR30B |
| TNFRSF1A | MSN |
| TPST1 | NCK1 |
| VPS45 | NCK2 |
| XCR1 | NCKAP1L |
| | NCSTN |
| | NDFIP1 |
| | P2RX7 |
| | PAWR |
| | PDCD1LG2 |
| | PDE5A |
| | PELI1 |
| | PIK3CG |
| | PLA2G2A |
| | PLA2G2D |
| | PLA2G2E |
| | PLA2G2F |
| | PLA2G5 |
| | PNP |
| | PPP3CA |
| | PPP3CB |
| | PRDX2 |
| | PRKAR1A |
| | PRKCQ |
| | PRNP |
| | PSMB10 |
| | PTPN22 |
| | PTPN6 |
| | PTPRC |
| | PYCARD |
| | RAC2 |
| | RASAL3 |
| | RASGRP1 |
| | RC3H1 |
| | RC3H2 |
| | RIPK2 |
| | RIPK3 |
| | RPS3 |
| | RPS6 |
| | SASH3 |
| | SCGB1A1 |
| | SCRIB |
| | SDC4 |
| | SELENOK |
| | SFTPD |
| | SH2D2A |

| | |
|---|---|
| | SH3RF1 |
| | SHH |
| | SLAMF1 |
| | SLC11A1 |
| | SLC4A2 |
| | SLC7A1 |
| | SOS1 |
| | SOS2 |
| | SPN |
| | SPTA1 |
| | STAT5B |
| | SYK |
| | TFRC |
| | TGFBR2 |
| | TMEM131L |
| | TMIGD2 |
| | TNFRSF13C |
| | TNFRSF14 |
| | TNFRSF1B |
| | TNFRSF21 |
| | TNFRSF4 |
| | TNFRSF9 |
| | TNFSF13B |
| | TNFSF14 |
| | TNFSF18 |
| | TNFSF4 |
| | TNFSF8 |
| | TNFSF9 |
| | TP53 |
| | TRAF6 |
| | TSPAN32 |
| | TWSG1 |
| | TYK2 |
| | VCAM1 |
| | VSIG4 |
| | VSIR |
| | VTCN1 |
| | WNT4 |
| | XCL1 |
| | ZAP70 |
| | ZBTB7B |
| | ZP3 |
| | ZP4 |
